# The adaptation front equation explains diversification hotspots and living-fossilization

**DOI:** 10.1101/2021.07.17.452749

**Authors:** Hiroshi C. Ito, Akira Sasaki

**Author notes:** Corresponding author: Hiroshi C. Ito, **Email:**.

## Abstract

Taxonomic turnovers are common in the evolutionary histories of biological communities. Such turnovers are often associated with the emergence and diversification of groups that have achieved fundamental innovations beneficial in various ecological niches. In the present study, we show that such innovation-driven turnovers could be analyzed using an equation that describes the dynamics of zero-fitness isoclines in a two-dimensional trait space comprising a “fundamental trait” (describing fundamental innovation) and a “niche trait” (describing niche position) or with its higher-dimensional extensions. Our equation allows analytical prediction of evolutionary source–sink dynamics along the niche axis for an arbitrary unimodal (or multimodal with weak separation) carrying capacity distribution. The prediction was confirmed by numerical simulation under different assumptions for resource competition, reproduction, and mutation. In the simulated evolution, biodiversity sources are the central niches having higher carrying capacities than the outer niches, allowing species there the faster evolutionary advancement in fundamental traits and their repeated diversification into outer niches, which outcompete the indigenous less advanced species. The outcompeted species go extinct or evolve directionally toward the far outer niches of the far slower advancement because of the far lower carrying capacities. In consequence of this globally acting process over niches, species occupying peripheral (i.e., the outermost) niches can have significantly primitive fundamental traits and deep divergence times from their closest relatives, and thus, they correspond to living fossils. The extension of this analysis for multiple geographic regions showed that living fossils are also expected in geographically peripheral regions for the focal species group.

**Significance Statement:** We developed a new equation for analyzing the long-term coevolution of many species through their directional evolution, evolutionary branching, and extinction in an arbitrary trait space comprising traits describing niche positions and traits describing fundamental innovations. This equation describes the many coevolving species as a fluid, as in the study of galaxy dynamics in astronomical physics. This equation may be used to explain how innovation-driven taxonomic turnovers generate diversification hotspots and coldspots among niches or geographical regions, accompanied by continuous production of “living fossil” species in peripheries, in a logic equivalent to that deduced by Darwin and Darlington from empirical observations.

## Introduction

Taxonomic turnovers are common in evolutionary histories of biological communities at various taxonomic levels (1). These turnovers are often associated with the emergence and diversification of groups that seem to have achieved fundamental innovations beneficial in various ecological niches, called “new organic forms” by Darwin (1859) (2), “general adaptation” by Peter (1871) (3), or “key innovations” by Miller (1949) (4). Examples of such innovation-driven turnovers include the diversification of angiosperms followed by the decline of gymnosperms (5) and the diversification of eutherian mammals replacing metatherian mammals (6).

For convenience, we refer to evolutionary traits related to fundamental innovations and those related to niche positions as “fundamental traits” and “niche traits,” respectively. Theoretical studies with numerical simulations have shown that joint evolution of a fundamental trait and a niche trait typically induces repeated taxonomic turnover through evolutionary branching of the more innovated groups and extinction of the less innovated groups (7, 8). Interestingly, through this turnover dynamics, species occupying the central niches (of the higher carrying capacities) tended to be more advanced in the fundamental trait than those occupying the outer niches (of the lower carrying capacities), having induced almost unidirectional evolutionary flow of species from the central to outer niches (7). This is concordant with the well-known empirical tendency, which states that older (or less advanced) groups in the same higher-level taxon occupy the outer positions in the niche space or morphospace for that taxon (9–12), and also with a long-lasting expectation that such an empirical tendency is generated through the evolutionary flow of species from optimum to suboptimum environments (or geographical regions) (2, 13–18).

Hence, by further analyzing the deterministic properties of the simulated taxonomic turnovers, we may obtain deeper insights into the mechanisms underlying taxonomic turnovers of real biological communities and resulting biodiversity patterns observed in existing and extinct biological communities. However, obtaining deterministic properties of multispecies coevolution involving evolutionary branching and extinction is difficult, because analytical tools for such dynamics are missing in current theoretical frameworks (19–22).

Combining species packing theory (23–26) and adaptive dynamics theory (22, 27–30), we developed a partial differential equation for describing many species’ coevolution involving evolutionary branching and extinction for arbitrarily dimensional trait spaces comprising fundamental traits and niche traits. Here we show an application of our equation for a simple resource competition model for the joint evolution of a fundamental trait and a niche trait. The obtained results, confirmed by numerical simulation, can explain the generation mechanism of “living fossils,” i.e., taxa characterized with significantly primitive morphologies, slow morphological evolution, low diversification and extinction rates, and taxonomical distinctness (31), such as ginkgo (*Ginkgo biloba*), tuatara lizard (*Sphenodon punctatus*), and horseshoe crabs (Limulidae). We also show that our analysis can be extended for describing geographical source–sink dynamics, which is related to taxon cycles (32, 33), taxon pulses (17), and the out-of-tropic hypothesis (i.e., latitudinal biodiversity source–sink dynamics) (34).

## Model for innovation-driven taxonomic turnovers

### Population dynamics

For analytical tractability, we assume asexual reproduction though obtained analytical results are examined by numerical simulation under both sexual and asexual reproduction. We define niche traits as evolutionary traits under negative frequency-dependent selection (i.e., the differentiation in niche traits facilitates species coexistence) and define fundamental traits as evolutionary traits under monotonic directional selection (i.e., the improvement in fundamental traits increases the fitness of species occupying any niches). We consider a two-dimensional trait space comprising a niche trait and a fundamental trait, denoted by two scalars *x* and *y*, respectively. For arbitrary *N* coexisting phenotypes, (*x*_1_, *y*_1_), …, (*x_N_, y_N_*), with their population sizes, *n*_1_, …, *n_N_*, we describe the *i*th phenotype’s per capita growth rate in the form of the classical MacArthur–Levins resource competition model (23, 35):

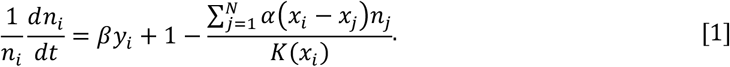

Here, *K*(*x_i_*) describes the carrying capacity for the *i*th phenotype, which is a Gaussian function with standard deviation *σ_K_* and peak *K*_0_ at *x* = 0:

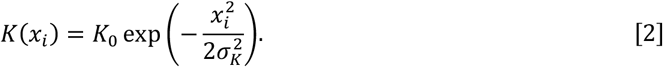

For a simple presentation of our analysis, we assume linear scaling of population size so that the maximum carrying capacity is equal to 1, i.e., *K*_0_ = 1. The *α*(*x_i_, – x_j_*) in Eq. 1, known as the competition kernel, describes the competition strength between the *i*th and *j*th phenotypes and is a Gaussian function with standard deviation *σ_α_* and peak 1 at (*x_i_, – x_j_*) = 0:

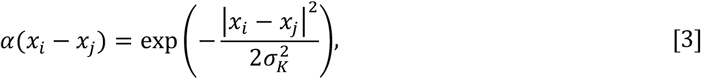

i.e., the strength of the competition between phenotypes *i* and *j* decreases with their niche difference |*x_i_ – x_j_*|. Note that increment of the īth phenotype’s fundamental trait, *y_i_*, always increases its fitness (Eq. 1), and that differentiation of its niche trait, *x_i_*, from other existing phenotypes’ niche traits also increases its fitness. When two phenotypes share the same *x*, the phenotype with a larger *y* competitively excludes the other. Hence, their coexistence requires their differentiation in *x*.

### Evolutionary dynamics

Evolutionary dynamics is induced by the repeated emergence of mutants produced from resident phenotypes and by selection on them through the population dynamics defined by Eqs. 1–3. For simplicity, we consider a trait space scaled beforehand so that the mutational covariance matrix is given by the identity matrix multiplied by 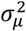, i.e., the mutation is isotropic with its average size (standard deviation) given by *σ_μ_*. We also assume that *σ_μ_* is sufficiently small so that in many cases evolutionary dynamics proceeds as directional evolution, evolutionary branching, and extinction of well-separated phenotypic clusters (27, 36). For convenience, we treat each phenotypic cluster as a species (although it could be a subspecies or strain). For analytical tractability, we also assume that the mutation rate *μ* per generation and per unit population size is low so that coexisting phenotypes are almost at their population-dynamical equilibrium whenever a mutant emerges. In this case, each phenotypic cluster is almost monomorphic (27).

For a mutant with phenotype (*x′*, *y′*) emerging from one of the coexisting species, its initial growth rate, called the invasion fitness, is expressed in a form similar to Eq. 1 as

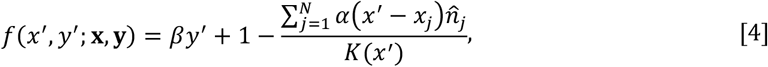

where **x** = (*x*_1_, …, *x_N_*) and **y** = (*y*_1_, …, *y_N_*) describe coexisting species’ niche and fundamental traits, respectively, and 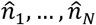 describe their equilibrium population sizes, which are functions of **x** and **y**. Here, the linear dependency of *f*(*x′*, *y′*; **x, y**) on *y′* gives the constant fitness gradient *β* along the *y*-direction at any position in the trait space. An initial ancestral species is expected to evolve directionally toward the optimum niche *x* = 0 as well as toward a larger *y*. Then, its evolutionary branching into two species along the *x*-direction is highly expected if the competition kernel has a narrower width than the carrying capacity, i.e., *σ_α_* < *σ_K_*, and if the fitness gradient *β* along the *y*-direction is sufficiently weak so that the branching line condition (37), 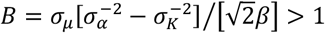 holds (see SI Appendix, section 1 for the derivation). Numerical simulation shows that the initial evolutionary branching is always followed by repeated branching and extinction (tree-like trajectories in Fig. 1; see SI Appendix, section 2A for the simulation algorithm [implemented in R]), analogously to (7) analyzing a model similar to Eqs. 1–3 in this study. Here, species occupying central niches tend to be more advanced in trait *y* than those occupying the outer niches, which seem to induce repeated species diversification from the central niches into the outer niches. Such dynamics can also arise under sexual reproduction with nonrare mutation (Fig. S1; see SI Appendix, section 2B for the simulation algorithm [implemented in C++]), analogously to (7).

**Figure 1.**
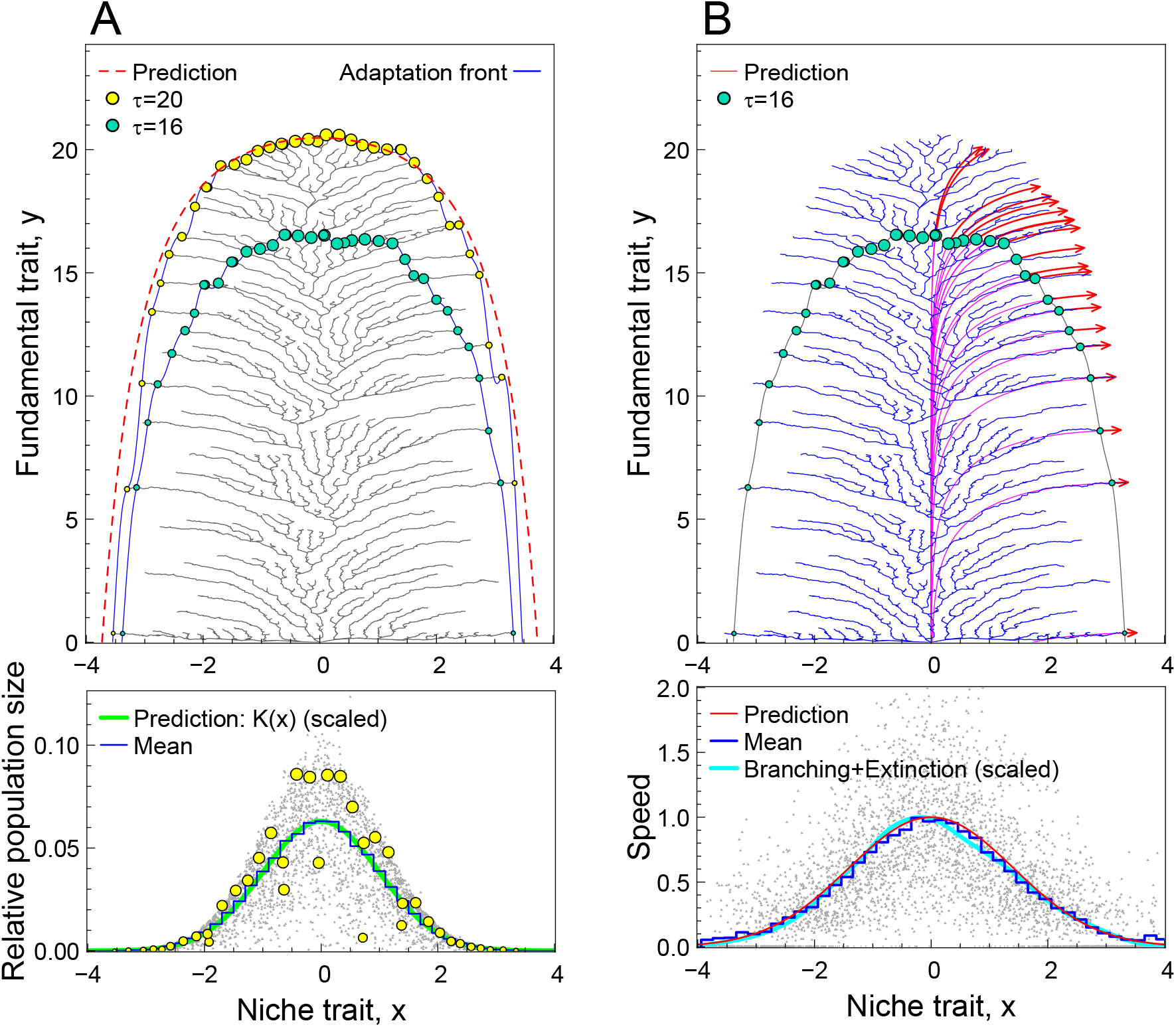
Predictions by the adaptation front equation for shape of community distribution and for speed and direction of evolutionary trajectories in simulated evolution. Upper panels in (A) and (B) show the same evolutionary dynamics (tree-like trajectories indicated with gray and blue, respectively) simulated for the resource competition model defined by Eqs. 1–3. The upper panel in (A) shows the adaptation front (blue curves), i.e., zero-fitness isocline, and coexisting species (colored circles) at time *τ* = 16 and *τ* = 20, where their population sizes are indicated with sizes of circles. The red dashed curve shows the predicted shape of the adaptation front by Eq. 6. (see the end of this legend for details). In the lower panel of (A), yellow circles indicate relative population sizes 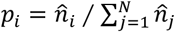 for coexisting species at *τ* = 20, light gray dots indicate those sampled over the simulation with their average for each *x* shown with the blue histogram. The green curve illustrates the prediction by the carrying capacity distribution, *K*(*x*) (Eq. 2), with its height scaled to attain the same peak height with the blue histogram. In the upper panel of (B), blue-green circles indicate coexisting species at *τ* = 16. Red arrows indicate the prediction for their evolutionary trajectories from *τ* = 16 to *τ* = 20 given by time integration of Eq. 7. Purple curves represent the backward prediction for the species’ trajectories from *τ* = 16. In the lower panel of (B), light gray dots indicate speeds of species’ directional evolution sampled over the simulation, with their average for each *x* indicated with the blue histogram. The red curve illustrates the prediction, 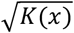 (Eq. 8). Model parameters: *β* = 0.1, *σ_K_* = 1, *σ_α_* = 0.15, *μ* = 1, *σ_μ_* = 0.01. The phenotype for the initial species was (*x, y*) = (−0.2, 0). Sampling was every 100 mutant invasions from *τ* = 0 to *τ* = 60. The prediction formula for the shape of the adaptation front is *y* = *y_H_* + *H*(*x*) with *H*(*x*) given by Eq. 6 and *y_H_* chosen so that the estimated community average 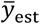 for trait *y* coincides with 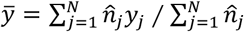 at *τ* = 20, where 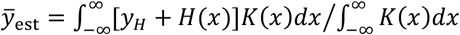. See SI Appendix, section 2A for the simulation algorithm.

## Adaptation front equation

In the adaptive dynamics theory originated from (27) (among several different formulations for adaptive dynamics, including (38–40)), directional evolutions of coexisting species are described with simultaneous ordinary differential equations, called the canonical equations (22, 30, 41). However, for many coexisting species, their analytical treatment is often difficult, as easily imagined from the complex tree-like trajectories in Fig. 1. Alternatively, the zero-isocline of invasion fitness referred to as the “adaptation front,” shows monotonic progress keeping roughly the same and simple shape (blue curves in Fig. 1A), suggesting its analytical tractability. Indeed, as explained in “Derivation of adaptation front equation” in Materials and Methods, we can approximately convert the canonical equations for many species into an analytically tractable partial differential equation that describes the expected dynamics of adaptation front, under the assumption of dense and uniform species packing along the niche and time axes. Such a species packing can be expected, according to the species packing theories (25, 26), if the competition kernel has a much narrower width than the carrying capacity, i.e., *σ_α_* ≪ *σ_K_*, and if the fitness gradient along the *y*-direction is weak, i.e., *β* ≪ 1, so that the much faster evolution of niche trait than that of fundamental trait keeps coexisting species’ niche traits close to evolutionary equilibrium corresponding to the ideal-free distribution (42). Under these assumptions, the whole community in the trait space may be treated as a kind of fluid consisting of the coexisting species as its molecules. The converted equation, referred to as the (rescaled) adaptation front equation, is given by

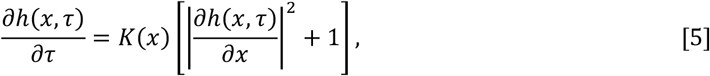

where *y* = *h*(*x,τ*) describes the adaptation front position in the fundamental trait at niche *x* at time *τ*. Here, the time *t* is rescaled into *τ*, so that the progress speed of adaptation front at *x* = 0 along the *y*-direction becomes equal to 1, i.e., *∂h*(0, *τ*)/*∂τ* = *K*(0) = 1, for sufficiently large *τ* (see “Time rescaling” in Materials and Methods). In this equation, terms in the square bracket mean that the progress of adaptation front, realized by the evolution of coexisting species around there, is locally orthogonal to the front, whereas the *K*(*x*) gives its speed. The *K*(*x*) comes from that the expected population size for a species occupying niche *x* is approximately proportional to *K*(*x*), as shown in the lower panel of Fig. 1A (see also Eq. 23 in Materials and Methods). Since all coexisting species must be located on the adaptation front at their population-dynamical equilibria, the shape of the adaptation front characterizes that of community distribution in the trait space (upper panel of Fig. 1A).

Eq. 5 can be compared to the Kardar–Parisi–Zhang universality class equations for describing the growing interfaces of bacterial colonies, crystals, paper wetting, etc. (43, 44). These equations are characterized by interface growth in the direction locally orthogonal to the interface (corresponding to |*∂h*(*x, τ*)/*∂x*^2^| in Eq. 5).

Strictly speaking, the adaptation front equation applies only when the dynamics of the adaptation front are driven solely by directional evolution of coexisting species, provided that evolutionary branching and extinction are negligibly rare. Nevertheless, even under nonrare evolutionary branching and extinction, as shown below, the equation has a good prediction performance not only for the shape of adaptation front but also for the corresponding source–sink dynamics along the *x*-axis, composed of their directional evolution, evolutionary branching, and extinction (see also SI Appendix, sections 3–7).

## Prediction by adaptation front equation

### Community distribution (adaptation front)

As long as *K*(*x*) is unimodal and peaked at *x* = 0, we expect that the adaptation front for a sufficiently large *τ* has a steady shape (i.e., |*∂h*(*x, τ*)/*∂x* = 1 for any finite *x*, from which we get *∂h*(*x, τ*)/*∂x*|^2^ = 1/*K*(*x*) – 1). Therefore, *h*(*x, τ*) is approximately given by

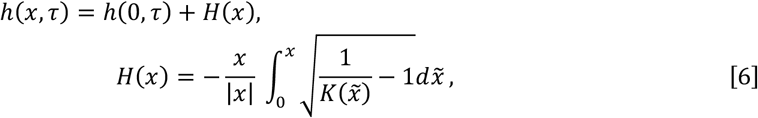

(see “Adaptation horizon” in Materials and Methods). We refer to *H*(*x*) as the “adaptation horizon.” Even for a nonisotropic mutation with average mutation sizes *σ_μxx_* and *σ_μyy_* along the *x*- and *y*-directions, respectively, the adaptation horizon has a similar form, given by [*σ_μyy_ / σ_μxx_*]*H*(*x*) (see “Adaptation horizon” in Materials and Methods). Note that *H*(*x*) is negative except for its maximum *H*(0) = 0.

As shown in Fig. 1A, the adaptation horizon (red dashed curve) successfully predicts the global shapes of adaptation front (blue solid curves) and community distribution (colored circles) in the simulated evolution, where the outer species, i.e., species occupying the outer niches, have less advanced fundamental traits.

### Directional evolution

Since the adaptation front describes the zero-fitness isocline, directional selection for each species is always orthogonal to the adaptation front. Hence, under the isotropic mutation assumed here, the expected directional evolution for each species is orthogonal to the adaptation front at its position. From this relationship, we derive the expected velocity of directional evolution for a species occupying niche *x* for sufficiently large *τ* (see “Directional evolution” in Materials and Methods) as

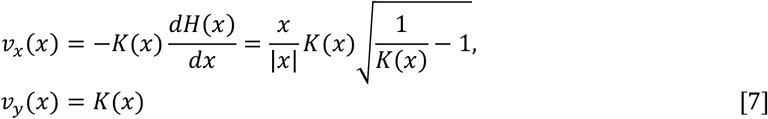

(red arrows and purple curves in the upper panel of Fig. 1B) with its speed and direction calculated as

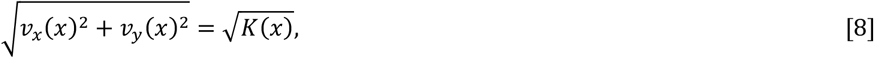

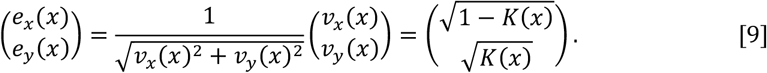

Eq. 8 indicates that the outer species evolve more slowly with the speed 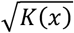 than do species at the center with the speed 1 (lower panel of Fig. 1B). Note that 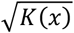 in Eq. 8 also coincides with the frequency distribution for either evolutionary branching or extinction, indicated with the cyan curve in the lower panel of Fig. 1B (with height scaling so that its maximum height becomes equal to 1). Thus, species occupying the outer niches have lower rates not only for their directional evolution but also for their evolutionary branching and extinction.

### Net diversification

Since the expected velocity of evolution *v_x_*(*x*) along the *x*-direction varies over *x*, the niche distance between neighboring species may increase or decrease, depending on their relative velocities. More specifically, the niche distance Δ*x* between a species *i* and its neighboring species *j*, occupying *x_i_* = *x* and *x_j_* = *x* + Δ*x*, respectively, is expected to increase when [*v_x_*(*x* + Δ*x*) – *v_x_*(*x*))]/Δ*x* ≃ *dv_x_*(*x*) / *dx* > 0 or decrease when *dv_x_*(*x*) / *dx* < 0. However, under the assumption of uniform species packing (25, 26), the expected between-species niche distance should be approximately kept constant over niches *x* through *τ*. This gap can be filled with the counterbalancing nonuniform rates of evolutionary branching and extinction over *x* (top panel of Fig. 2). From this perspective, we obtain the following relationship by applying the continuity equation in fluid dynamics (45):

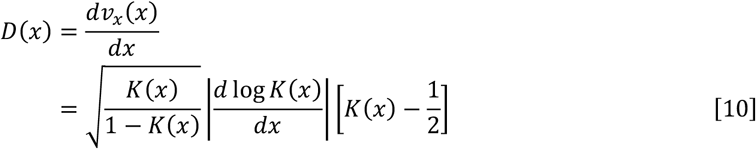

(see “Net diversification” in Materials and Methods), where *D*(*x*) describes the net diversification rate, defined as the number of branching subtracted by that of extinction, expected for species occupying niches around *x*, per species, and per unit *τ*. Eq. 10 effectively predicts the net diversification rate in the simulated evolution (middle panel of Fig. 2). Eq. 10 shows that the outer niches satisfying *K*(*x*) < 1 / 2 are always expected to be sinks (i.e., *D*(*x*) < 0), under any unimodal *K*(*x*) with its peak height scaled to 1 (bottom panel of Fig. 2). Moreover, we see from Eq. 10 that the magnitude of the maximum net diversification rate, attained at *x* = 0, is estimated from the curvature of carrying capacity at *x* = 0: 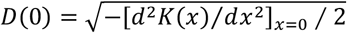, e.g., 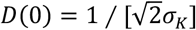 for the Gaussian carrying capacity defined by Eq. 2.

**Figure 2.**
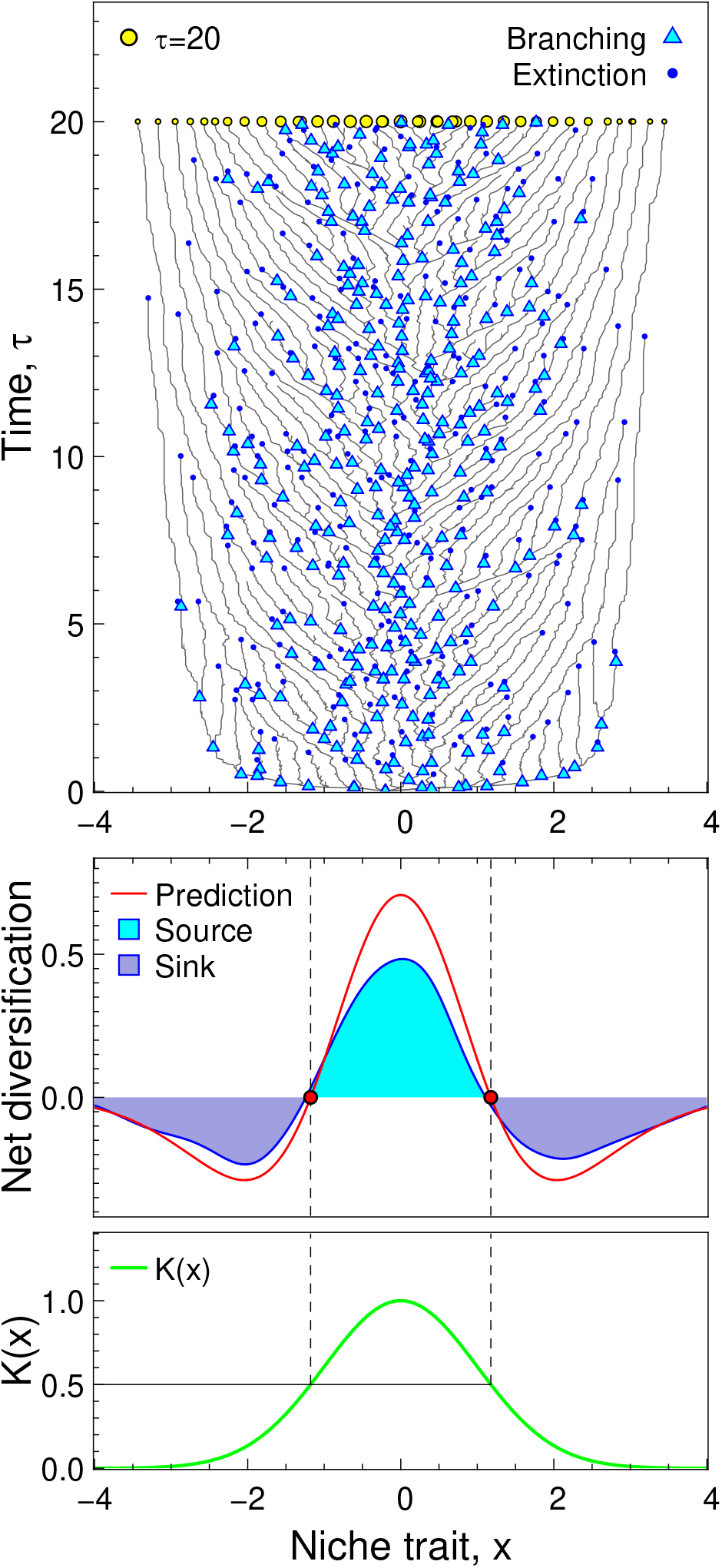
Prediction by the adaptation front equation for net diversification rate (for the simulated evolution shown in Fig. 1). The top panel shows species’ trajectories (gray curves), their evolutionary branchings (cyan triangles), and their extinctions (blue dots), in the “niche–time” (*x-τ*) plane. Coexisting species at *τ* = 20, shown in Fig. 1A, are indicated with yellow circles. The bottom panel shows carrying capacity distribution, defined by Eq. 2. The middle panel shows net diversification rate (number of evolutionary branching subtracted by that of extinction expected for species occupying niches around *x* per species and per unit time in *τ*) indicated with a blue curve filled with cyan or blue, corresponding to positive (i.e., source) and negative (sink) net diversification rates, respectively (smoothed with Gaussian-moving-average filter with standard deviation *2σ_α_*). The red curve is the prediction by Eq. 10 for the net diversification rate. The source–sink boundaries are always expected at the niche positions where their carrying capacities are equal to 1 / 2 (red circles) for any unimodal carrying capacity distribution with its peak scaled to 1.

### Phenotypic distance and divergence time

In the simulated evolution, the outer species tend to be more phenotypically distant from their closest relatives (Fig. 3A) and have deeper divergence times from their closest ones (Fig. 3B). Concerning the phenotypic distance, we can predict the magnitude of phenotypic distance *p_d_* in the *x-y* space for a species occupying niche *x* from its closest relative, as the corresponding length of the slope of adaptation horizon (upper panel of Fig. 3A), under the assumption of constant between-species niche distance, *σ_α_*, for the Gaussian competition kernel (25, 26). Specifically, the prediction formula for *p_d_* is given by

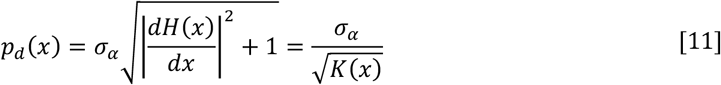

(red curve in the lower panel of Fig. 3A). Moreover, as explained in “Divergence time” in Materials and Methods, the divergence time *τ_d_*(*x*) for a species occupying a sufficiently outer niche *x* from its closest relative coincides approximately with the difference of its fundamental trait (i.e., *h*(*x, τ*)) from the front top (i.e., *h*(0, *τ*)), so that the following relationship approximately holds:

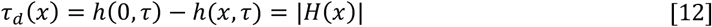

(red curve in the lower panel of Fig. 3B).

**Figure 3.**
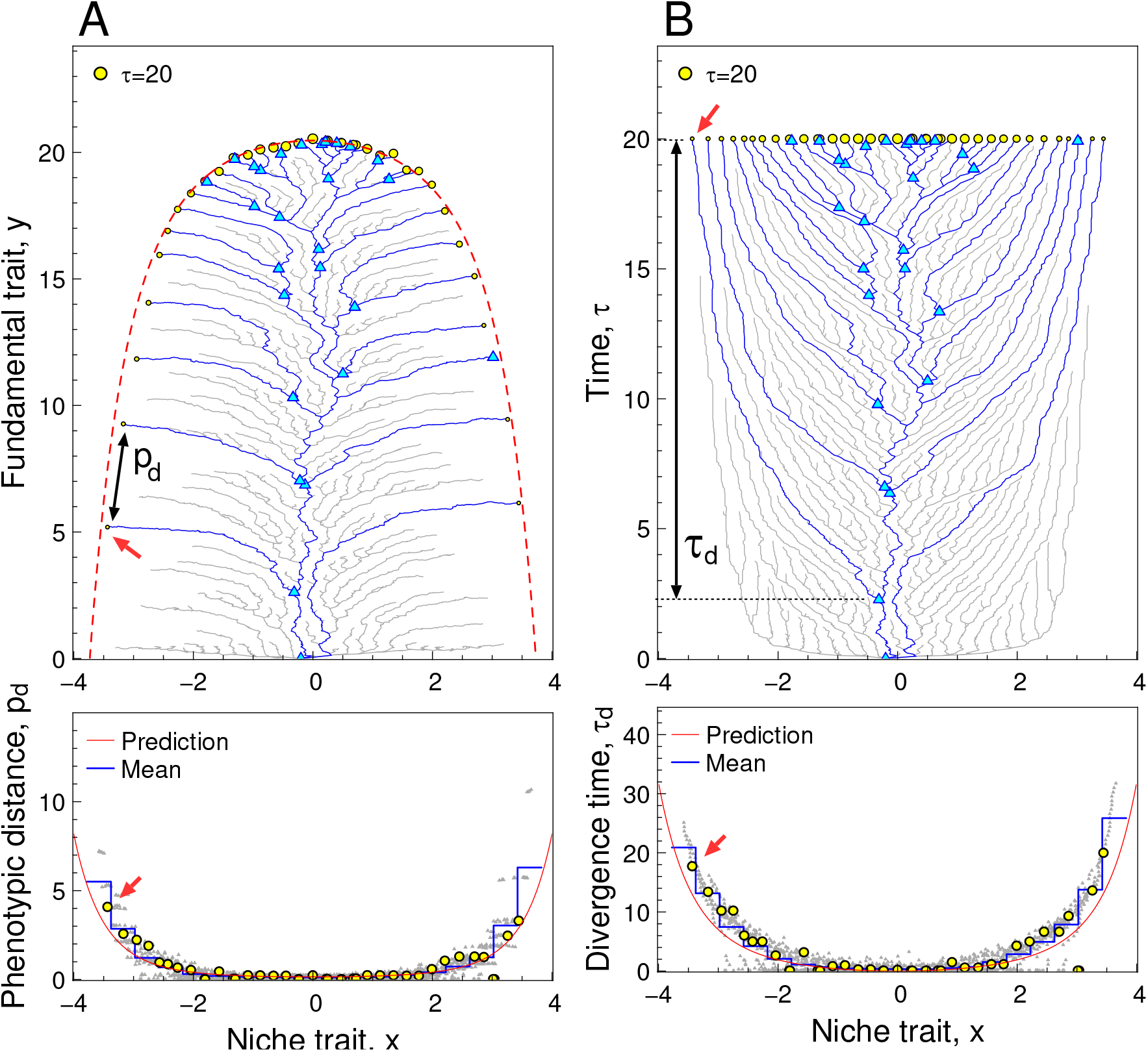
Prediction by the adaptation front equation for phenotypic distances and divergence times among coexisting species (for the simulated evolution shown in Fig. 1). In the upper panel of (A), blue curves show trajectories (in the trait space) of coexisting species at *τ* = 20 shown with yellow circles. Gray curves show trajectories of species extinct at *τ* = 20. Cyan triangles indicate branching points among the coexisting species at *τ* = 20. The red dashed curve shows the adaptation horizon, *y* = *y_H_* + *H*(*x*) (see caption of Fig. 1 for *y_H_*). In the lower panel of (A), each yellow circle indicates the phenotypic distance for each species from its closest relative, denoted by *p_d_*, and light gray dots indicate those sampled over the simulation with their average for each *x* shown with the blue histogram (equally spaced 40 samplings from *τ* = 0 to *τ* = 60). The red curve shows the prediction for *p_d_* as a function of *x* (Eq. 11). The upper and lower panels of (B) show the relationship between *x* and *τ*, instead of *x* and *y* shown in (A), in a manner analogous to (A), where *τ_d_* denotes the divergence time for each species from its closest relative. The red curve shows the prediction for *τ_d_* as a function of *x* (Eq. 12).

## Living-fossilization

In the above results, the outermost species (i.e., species occupying the outermost niches) are characterized by their (i) primitive fundamental traits (Fig. 1A), (ii) slow adaptive evolution (Fig. 1B), (iii) low diversification and extinction rates (Fig. 1B), and (iv) large phenotypic distances and deep divergence times from their closest relatives (Fig. 3). Hence, the outermost species seem to correspond to “living fossils,” which are extant or extinct taxa characterized with properties including (I) primitive morphologies, (II) slow morphological evolution, (III) low diversification and extinction rates, and (IV) outstanding taxonomical distinctness (31, 46, 47). For convenience, we denote by “living-fossilization” the evolutionary process of getting relatively more primitive in fundamental traits due to slowing of their adaptive evolution, in comparison with the most advanced species in the focal species group. The globally convex adaptation horizon (dashed red curve in the upper panel of Fig. 3A) implies that most species that once deviated from the optimal niche (*x* = 0) undergo living-fossilization and are to go extinct, and some of which that are locally stronger than their neighbors may live extremely long, in exchange for their significant living-fossilization (e.g., a species highlighted with red arrows in Fig. 3).

Although living-fossilized species occupying the outermost niches are far more primitive than the advanced ones occupying the central niches, the advanced ones do not easily exclude the primitive ones. This is because the advanced ones inevitably experience slower adaptive evolution when they evolutionarily shift toward the outer niches. Consequently, the advanced ones’ reaching the outermost niches take very long, inevitably accompanied by their living-fossilization.

## Robustness of living-fossilization

Here, we explain how robust the predictions by the adaptation front equation are (i.e., Eqs. 6–12) against more general ecological and genetic settings. In all cases that we have examined below (except for one of the two sides of the carrying capacity distribution under the asymmetric competition kernel), the outermost species are characterized by their least advanced fundamental traits, slow adaptive evolution, low diversification and extinction rates (with negative net diversification rates), and deep divergence times (and large phenotypical distances) from their closest relatives. In this sense, the process of living-fossilization seems qualitatively robust, as far as we have examined.

### Sexual reproduction

Our predictions still work to some extent even under sexual reproduction and nonrare mutation, as long as the evolution of assortative mating required for evolutionary branching is not difficult (SI Appendix, section 3), analogously to (7).

### Varied shapes of carrying capacity distribution and competition kernel

Our predictions have good performance even when the carrying capacity has a different shape from the Gaussian, as long as it is unimodal or multimodal but with weak separation (SI Appendix, section 4), or when the fitness gradient for the fundamental trait is not constant but depends on niche trait (SI Appendix, section 5). In contrast, prediction performance qualitatively decreases when the competition kernel is platykurtic, i.e., its tail is thinner than the Gaussian (SI Appendix, section 6A), possibly because platykurtic competition kernels can cause excess stabilizing selection on coexisting species’ niche traits (26). Qualitative decrease in prediction performance also arises when the competition kernel is strongly asymmetric (SI Appendix, section 6B). In such a case, coexisting species experience excess directional selection toward the larger (or smaller) values for the niche trait. The low prediction performances under the platykurtic and strongly asymmetric competition kernels seem to be related to the violation of the main assumption for adaptation front equation that the community state is kept close to the evolutionary ideal-free distribution (Figs. S10D and S11D).

### Rare mutation for fundamental trait

When mutation for the fundamental trait, *y*, is significantly rarer but larger than that for the niche trait, *x*, each invading mutant toward more advanced *y*, referred to as the innovated mutant, triggers repeated diversification along the *x*-direction to form a species clade sharing the same y, excluding the other non-innovated species, until it is invaded by another innovated mutant (Fig. 4). This turnover process through diversification of the innovated species and extinction of non-innovated species acts as species-level selection (48–50). For the simulated evolution shown in Fig. 4, for example, we can estimate from model parameters that the advancement speed of the whole community in the *y*-direction becomes faster than the prediction (by the adaptation front equation) by factor *η* = 9.9 (see SI Appendix, section 7B and Fig. S14). The faster advancement speed causes a steeper adaptation front and is effectively predicted by a corrected adaptation horizon, 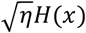 (red dashed curve in Fig. 4A). The corrected adaptation horizon successfully predicts that the outermost species are characterized by their least advanced fundamental traits, slow adaptive evolution, low diversification and extinction rates, and deep divergence times (and large phenotypical distances) from their closest relatives (Fig. S12), similarly to the case of isotropic mutation (Figs. 1–3). Moreover, the corrected adaptation horizon allows us to predict clade-level tendencies that the outer clades have smaller numbers of species (Fig. 4C) and older ages (Fig. 4D) than those occupying the central niches.

**Figure 4.**
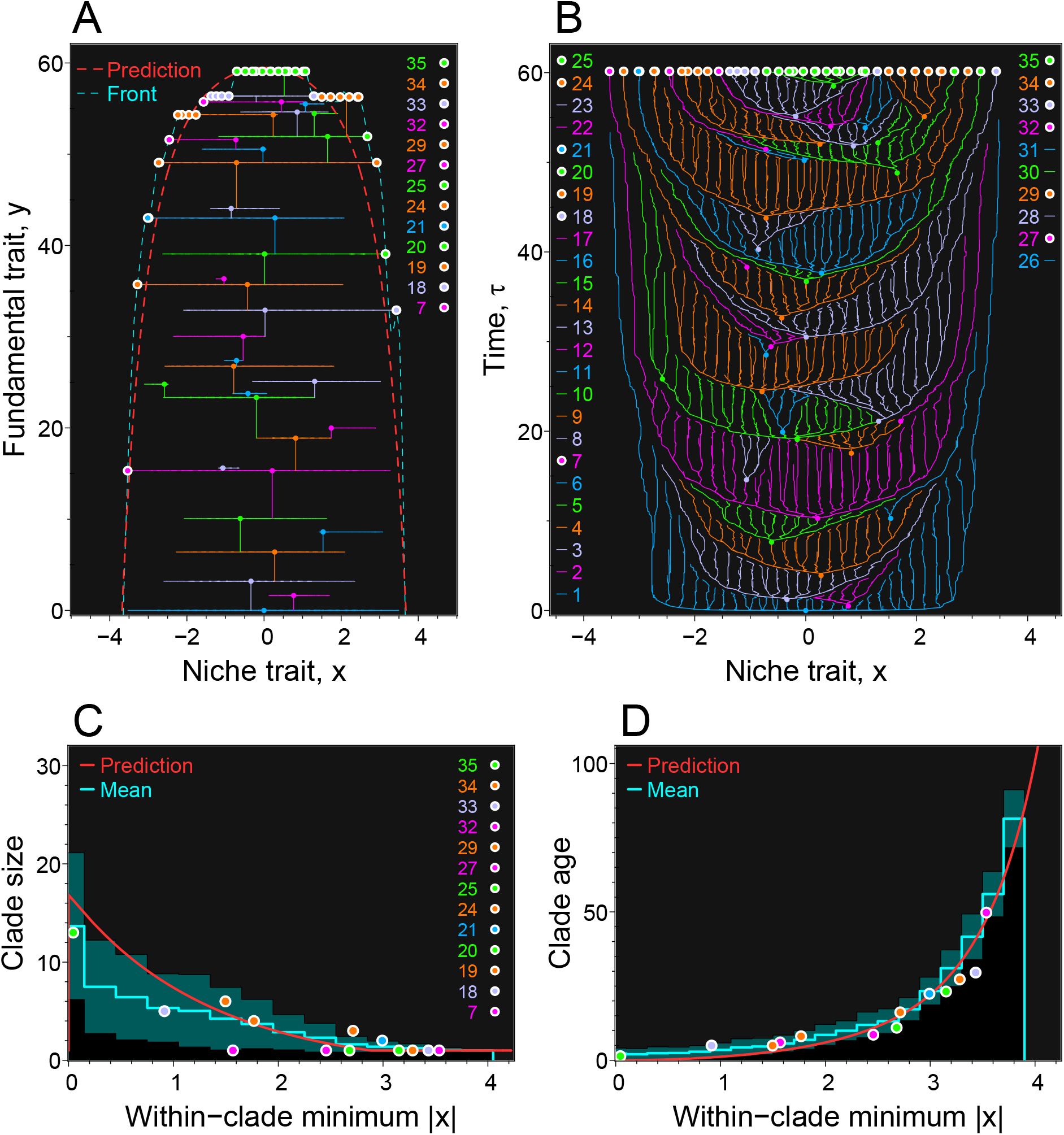
Clade-level dynamics in simulated evolution under much rarer and larger mutation for fundamental traits than for niche traits. Panels (A) and (B) show the simulated evolution in the *x-y* plane (trait space) and *x-τ* plane, respectively, where each subtree in (B) with one of five colors indicates a clade containing species sharing the same value for *y*. Integers assigned to these clades indicate their orders of origination, and colored tiny dots in (A) and (B) indicate their founders’ phenotypes and emergence times, respectively. Colored circles bordered with white indicate coexisting species at time *τ* = 60. The cyan and red dashed curves in (A) show the adaptation front and the corrected adaptation horizon 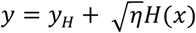 with the estimated effect of species-level selection, *η* = 9.92 (see caption of Fig. 1 for *y_H_*). In panel (C), colored circles bordered with white indicate clade size (number of species) and “within-clade minimum |*x*|” at *τ* = 60, where the within-clade minimum |*x*| for each clade was calculated as the minimum absolute value for the niche trait, *x*, among its component species. The cyan histogram with dark blue bands shows average clade sizes with standard deviations among clades sampled over the simulation (every 40 invasions from *τ* = 0 to *τ* = 150). Panel (D) plots the relationship between within-clade minimum |*x*| and clade age in a manner analogous to panel (C). The prediction shown in (C) (red curve) was numerically obtained from the corrected adaptation horizon, whereas the prediction shown in (D) is directly given from the corrected horizon by 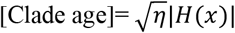 with *x* = [within-clade minimum |*x*|] (see SI Appendix, section 7C for details).

## Extension for higher-dimensional trait spaces

From an arbitrary fitness function for an arbitrary trait space of arbitrary dimension, the adaptation front equation can be derived in a form analogous to Eq. 5, provided that the trait space can be decomposed into a subspace for niche traits with narrow competition range and into that for fundamental traits undergoing monotonic and weak directional selection, and that population dynamics is kept close to equilibrium (see SI Appendix, section 8). The effect of niche dimensionality on evolutionary dynamics can be shown clearly by extending the original resource competition model defined by Eqs. 1–3 for an arbitrary isotropic *L*-dimensional niche space **x** = (*x*_1_, …, *x_L_*)^T^ with isotropic average mutation size, isotropic competition kernel, and isotropic carrying capacity distribution denoted by *K*(*r*) with *r* = |**x**|. In this case, the adaptation horizon and expected directional evolution are equivalent to Eqs. 6–9 with replacement of *x* with *r* (see SI Appendix, section 8F). However, the net diversification rate

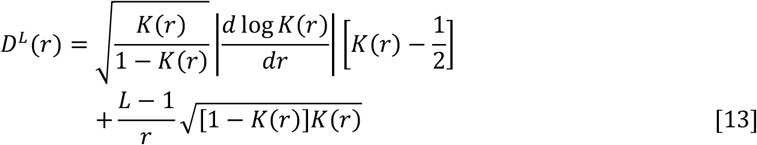

is different from that for the original model (Eq. 10), by the second term at the right-hand side, which is nonnegative and increases with the dimension *L* of the niche space. Hence, the maximum net diversification rate, *D^L^*(0). grows with the niche dimensionality. For example, if *K*(*r*) is the Gaussian distribution with a standard deviation *σ_K_*, Eq. 13 gives 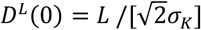.

## Extension for multiple geographic regions

Our model defined by Eqs. 1–3 corresponds to a single geographic region such that migration within the region is much faster than evolution. Here, we extend our model for multiple regions, among which migration is difficult (see SI Appendix, section 9A). For simplicity, we assume that these regions are aligned along a one-dimensional spatial axis *z*, and their positions are denoted by integer values (Fig. 5A). Carrying capacity distributions in these regions share the same shape with the single region model but differ in their heights so that the central region (*z* = 0) has the highest peak equal to 1 at *x* = 0, whereas the outer regions have lower peaks. With the continuous approximation of *z*, we can derive the spatially extended adaptation front equation (see SI Appendix, section 9B).

**Figure 5.**
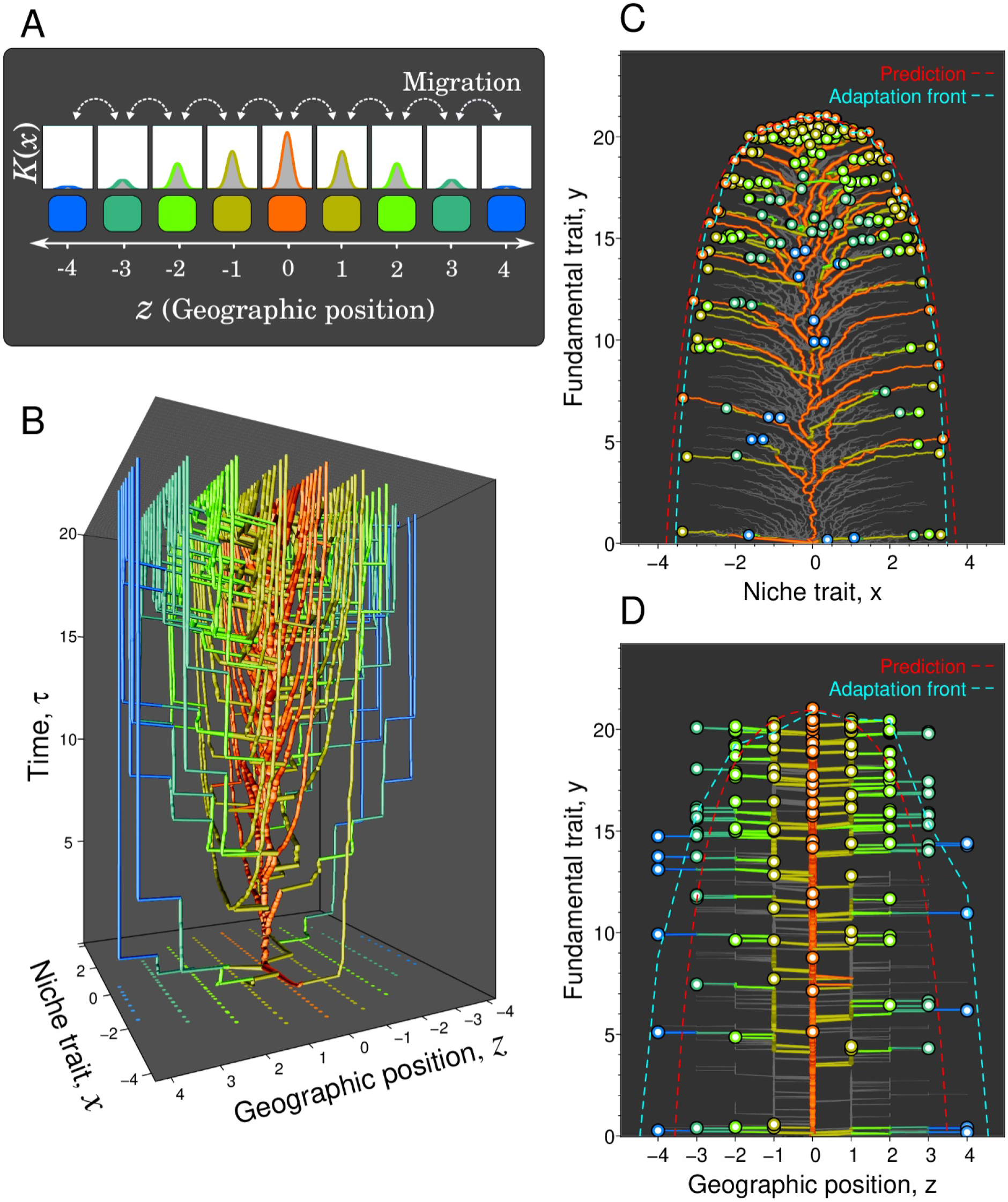
Simulated evolution and prediction under multiple geographic regions. Panel (A) illustrates the extension of the resource competition model defined by Eqs. 1–3, so that the system has multiple geographic regions (colored rectangles) aligned along the *z*-axis. These regions have the same carrying capacity as in Eq. 2, but their heights are lower in the outer regions (colored curves in white boxes). Panel (B) projects the simulated evolutionary trajectories in four-dimensional space, (*x, z, y, τ*) onto the *x-z-τ* space, where tubes indicate evolutionary trajectories of coexisting species at *τ* = 20, with their widths and colors indicating their population sizes and geographical positions. Filled circles on the bottom plane indicate phenotypes of the coexisting species at *τ* = 20. Panel (C) projects the simulated evolutionary trajectories onto the *x-y* plane, where the blue and red dashed curves show the adaptation front and adaptation horizon for the central region (*z* = 0). Colored circles with their inside white indicate coexisting species at the final time point. Trajectories of species that were extinct at *τ* = 20 are indicated with a light gray color. Panel (D) projects the simulated evolutionary trajectories onto the *z-y* plane similarly as in panel (C). The blue and red dashed curves show adaptation front and adaptation horizon as functions of *z* with fixed *x* at the optimum niche (*x* = 0). The carrying capacity distribution is defined by 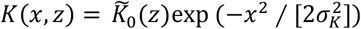 with 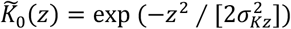. Model parameters: *σ_Kz_* = 1, *μ_m_* (successful migration rate between neighboring regions) = 5 × 10^-5^, and other parameters are the same as in Fig. 1. The initial species has phenotype (*x, z, y*) = (0,0,0). See SI Appendix, section 9B for details of predictions by the adaptation front equation.

The simulated evolution forms evolutionary trajectories in the four-dimensional space (*x, z, y, τ*), projection of which onto the *x- z- τ* space, the *z-y* plane, and the *x- y* plane are shown in Fig. 5B, 5C, and 5D, respectively. Although continuous approximation of the discrete *z* affects the prediction performance of the adaptation front equation, the adaptation horizon (Fig. S15A) effectively characterizes the adaptation front in the simulation (Fig. 5C and 5D). Without migration among regions, evolutionary dynamics in each region must be equivalent to that in the single region model (Figs. 1-3), except that dynamics in the outer regions proceed more slowly because of their lower carrying capacities, resulting in their slower advancement speeds in trait *y*. Hence, with migration, successful migrations tend to occur asymmetrically from the central regions to the outer regions (Fig. 5B and 5D). Consequently, only the central region keeps evolutionary dynamics similar to that of the single region model (Fig. 5C), whereas species in the outer regions tend to be excluded repeatedly (before their niche diversification) by migrators from the more central regions (Fig. 5D). This tendency is quantitatively visualized in Fig. 6A, as the time-averaged directional evolution and net diversification rate for each position on the *x-z* plane, which is well characterized by the prediction from the adaptation front equation (Fig. 6B). Note that living-fossilization acts over geographical regions (Fig. 5D) as well as over niches (Fig. 5C) so that all peripheral species near the boundary of existence area on the *x-z* plane experience high degrees of living-fossilization (Fig. S15B). Although some peripheral species have shallow divergence times, species having significantly deep divergence times are almost always restricted to peripheries characterized by significantly low carrying capacities (Fig. S16).

**Figure 6.**
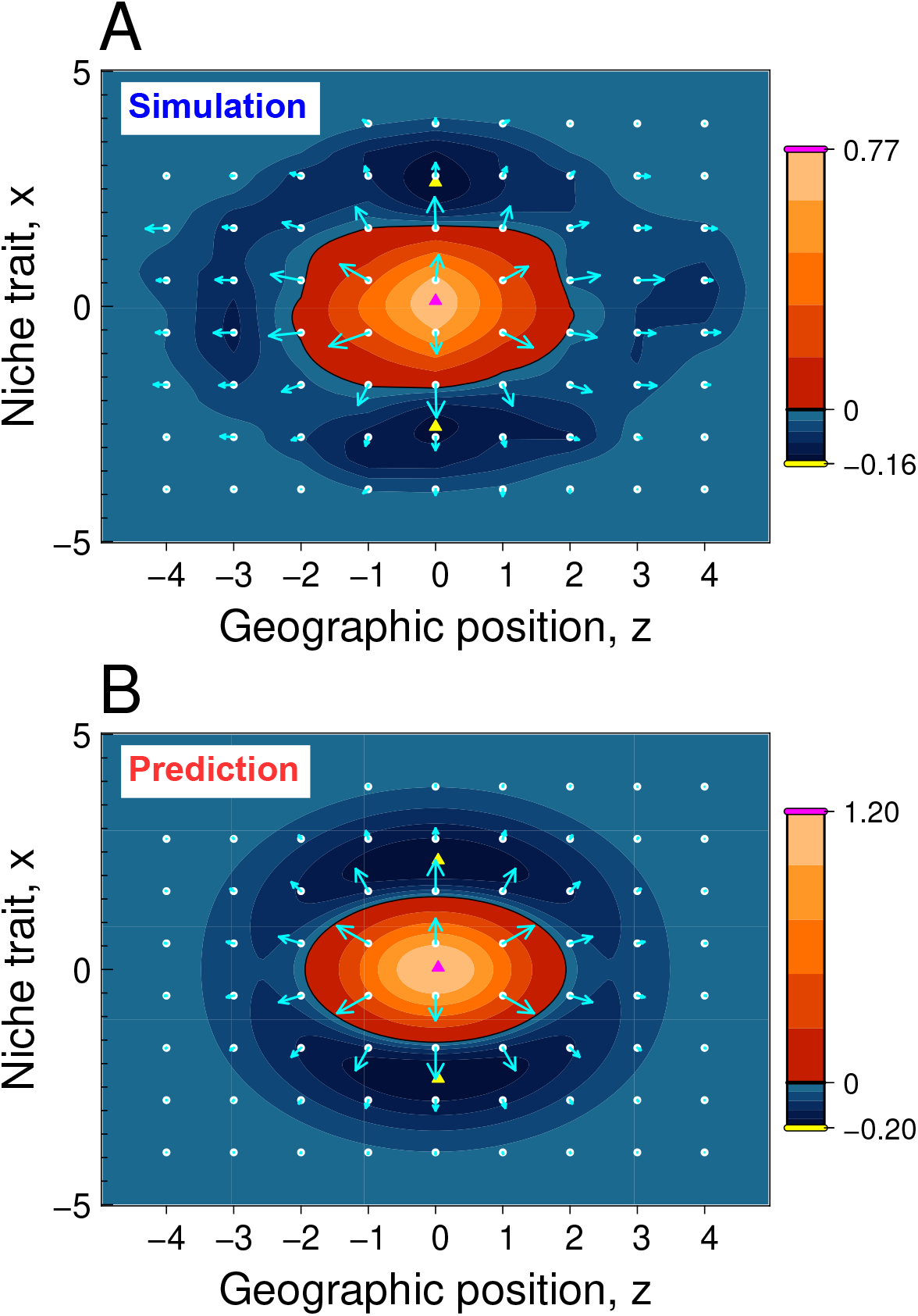
Prediction by the adaptation front equation for diversification hotspots and coldspots. Panels (A) and (B) compare simulation and prediction on the time average of directional evolution (cyan arrows) and that of net diversification rate (filled contours) for the simulated evolution shown in Fig. 5. In each panel, lengths of cyan arrows indicate speeds of directional evolution. The black closed curve indicates the boundary between positive (reddish gradient) and negative (blueish gradient) net diversification rates, corresponding to sources and sinks. The maximum and minimum values for the net diversification rate are indicated with numbers at the top and bottom, respectively, of the color bar on the right side. A hotspot and two coldspots corresponding to the highest and the two lowest values for net diversification rate are indicated with purple and yellow triangles. Sampling was every 4000 invasions by mutants or migrators from *τ* = 0 to *τ* = 50. See SI Appendix, section 9B for details of predictions made in this figure.

## Discussion

### Summary

The biological populations are thought to have been evolving in both fundamental and niche traits simultaneously. Using the adaptation front equation and numerical simulation, we have shown that the joint evolution is likely to induce repeated diversification around the central niches or the central geographic regions (that attain the fastest advancement in fundamental trait) and species flow from there toward the outer niches or regions (of slower advancement), accompanied by their living-fossilization, provided that the variation in the carrying capacity primarily characterizes the variation among niches or geographical regions. Below, we discuss our results in connection with relevant empirical and theoretical studies.

### Living fossils

Ever since Darwin (1859) (2), living fossils, characterized by their significantly primitive morphologies, slow morphological evolution, low diversification and extinction rates, and taxonomic distinctness (31, 46, 47), have attracted significant attention. As Darwin wrote, “Species and groups of species, which are called aberrant, and which may fancifully be called living fossils, will aid us in forming a picture of the ancient forms of life” (2), living fossils might provide information regarding organisms that lived in ancient eras. However, the scientific plausibility of the concept of living fossils has been in a long controversy (31). Thus, it is crucial to examine whether living fossils may be generated through plausible ecoevolutionary processes. A few theoretical studies are relevant to this perspective (7, 8, 51), all of which are, however, based only on numerical simulation. To our knowledge, this analysis is the first to show that living fossils can be generated through deterministic coevolutionary dynamics induced by competition. Furthermore, the results obtained from both analytical prediction and numerical simulation showed that living fossilization is a globally acting process over niches and geographic regions rather than a local process acting on particular types of niches or lineages.

The formula for adaptation horizon (*H*(*x*) in Eq. 6) indicates that a high degree of living-fossilization is expected for peripheral species along a niche or a spatial axis. Moreover, according to the adaptation horizon for nonisotropic mutation, given by [*σ_μyy_ / σ_μxx_*]*H*(*x*), the degree of living fossilization for a peripheral species is enhanced if (1) mutation is not easy but still possible along the axis (corresponding to small but nonzero *σ_μxx_*). Additionally, as we assumed in our model, (2) outer niches should stably remain outside, having lower carrying capacities than central, and (3) the environments in outer niches should sufficiently be stable for small populations to avoid extinction by environmental fluctuation, for living-fossilization to be expected robustly.

Among various niche axes, the sea depth may fit with the above three conditions. As for condition 1, adaptation to high water pressure seems difficult (52, 53), yet hundreds of deep-dwelling species (e.g., microbes, protists, worms, Porifera, Mollusca, Echinodermata, Crustacea, Cnidaria, and fishes) are found in the deepest areas of the ocean between 6,000 and 11,000 m, known as the hadal zone (54, 55). As for condition 2, the abundance and biomass of most phyla generally decrease with increasing depth (56). As for condition 3, although deep sea environments may be less stable than previously thought (57), some areas in deep sea provide stable environments (58), including cold seeps with the steady release of fluids from the seafloor for thousands of years at a single seep (59, 60).

### Darwin–Darlington principle and related patterns

The mechanism for innovation-driven evolutionary source–sink dynamics among niches and geographical regions, clarified in our present analysis, would have a strong connection with the hypothesis called the Darlington’s principle (35), originated from Darwin (2) and developed by Darlington (15, 16, 48) (related to (13, 14, 18)). Darlington defined “general adaptation” as an adaptation involving all improvements of structure and function that increase fitness in many “environments” (in many niches in the present terminology) (15). Then, for a focal taxonomic group, faster general adaptation was expected to be attained in more favorable environments (such that species of larger populations are maintained) and in larger geographic regions (such that larger numbers of species are maintained), through individual-level selection and species-level selection. In consequence, repeated evolutionary diversification was expected in the optimal environments and geographic regions (attaining the fastest general adaptation), inducing species flows to suboptimal ones with repeated exclusion of old species there (16, 48). In this dynamics, the complete adaptive and geographic history of a lineage from its monophyletic beginning to its end (i.e., extinction of all its progeny) was called taxon pulse (17).

Since Darwin’s contribution to the principle described above seems important (61), we here refer to the principle as the “Darwin–Darlington principle” or “DD principle.” The DD principle is related to the recent Darwinian framework of invasion biology (62), showing that floristic geographic regions of higher phylogenetic diversity are more likely to be source areas of invasive plants, whereas regions of lower phylogenetic diversity are more likely to be invaded. Moreover, since the “environments” and the “general adaptation” in the DD principle respectively correspond to the niches and the advancement in fundamental traits in our present study, the mechanism for innovation-driven evolutionary source–sink dynamics clarified in our present study corresponds to the DD principle, especially when mutation for the fundamental trait is rare (in which case geographical regions with the larger species diversity may attain faster potential speeds for community advancement, as derived in SI Appendix, section 7D). In this sense, our present analytical and numerical results underpin the plausibility of the DD principle. It is notable that Darwin (2) already expressed species flows among geographical regions as flows of “living waters,” implying the applicability of the fluid approximation conducted in our present study. Below, we refer to innovation-driven source–sink dynamics of species (or higher taxonomic groups) among niches or geographic regions or both as “Darwin–Darlington flows” or “DD flows.”

The DD flows may be expected along with any niche or spatial axis, as long as that axis satisfies condition two listed in the previous subsection: a stable cline in carrying capacity along the axis. Because lower temperatures are always expected at higher altitudes and latitudes, as well as at deeper depths in water, and because too low temperatures inevitably limit distributions of organisms, the DD flows may be common along with altitude, latitude, and water depth, and the former two of which are described in the literature for the DD principle (63) or taxon pulse (17). Empirical reports related to such species flows come from various taxonomic groups, along the habitat elevation [in ants (33, 64), carabids (17), and corvoid birds (65)], along latitude [in marine bivalves (66, 34, 67)], and along onshore–offshore, which may correspond to the sea depth [in marine bivalves (68, 69) and fishes (70)].

Regarding the ecological and geographical periphery of small and old clades, which is an expected outcome of DD flows (17, 48) underpinned by this study, recent empirical supports come from birds (10, 12) and bivalves (11, 67); species-poor clades are associated with their old ages and morphological, environmental, or geographical peripheries.

### Mechanisms for evolutionary taxonomic turnovers

The DD principle is one of the mechanisms that can induce taxonomic turnovers through repeated evolutionary branching and extinction. The other mechanisms proposed so far for repeated branching and extinction include environmental stochasticity (71), asymmetric competition (72–74), and predator–prey interaction (42, 75–77). The DD principle is not exclusive to environmental stochasticity (analyzed by (8)) nor with asymmetric competition (SI Appendix, section 6B). Regarding predator–prey interaction, the adaptation front equation may be extended for multiple trophic levels, provided that traits of predators and prey undergoing frequency-dependent selection (i.e., niche traits) are all kept close to evolutionary equilibria that correspond to evolutionary ideal-free distributions (42). Thus, the DD principle has a high affinity with other mechanisms for explaining evolutionary taxonomic turnovers through repeated evolutionary branching and extinction.

### Taxon cycle

In a broad sense, the taxon cycle hypothesis (32, 33) argued that taxa at all levels but most noticeably and accessibly at the species and subspecies level in island settings (archipelagos). Such species or subspecies go through continuous shifts in geographic range, followed by a slow decline toward extinction, with associated evolutionary shifts in their ecology and morphology (65, 78). Thus, the taxon cycle describes a phenomenon similar to the DD principle (63). Taxon cycles have been explained by asymmetric competition (79), interaction with enemies (predators, parasites, or pathogens) (80), or the ontogeny of the islands themselves (81). In the well-known case of the Lesser Antilles (79) bird groups, some species show deterministic declines toward extinction, but each monopolizes an island. Thus, direct competition among related species is unlikely to be the driving force of this taxon cycle, suggesting the interaction with predators, parasites, or pathogens as potential drivers (78, 80). If these enemies have higher dispersal abilities than the focal taxonomic group undergoing the cycle, the enemies can induce apparent competition among species inhabiting different islands. In this case, even a species monopolizing an island may go extinct because the smaller islands sustain only the smaller populations, which can cause their slower adaptive evolution in resistance abilities against enemies than species inhabiting the larger islands. We may translate these anti-enemy resistance abilities into fundamental traits, which allow us to analyze the stage shifts of species in the taxon cycle by our adaptation front equation.

### Applicability of adaptation front equation

In principle, the adaptation front equation can describe many species’ coevolution in an arbitrary trait space that can be decomposed into a subspace for niche traits (which may include spatial axes) and a subspace for fundamental traits, under the assumption of dense species packing in the niche space. This assumption corresponds to the fluid approximation of the whole coexisting species, like a fluid approximation of power flows, queues, communications in complex networks (82), and galactic dynamics in astronomy (83). As a prototypic descriptor for Darwin–Darlington flows, the adaptation front equation may provide a basis for a stronger connection among evolutionary community ecology, invasion biology, geographical ecology, and paleontology.

## Materials and Methods

### Derivation of adaptation front equation

Under a sufficiently small average mutational step size *σ_μ_* and a sufficiently low mutation rate *μ*, as assumed in the main text, the expected directional evolution of the *i*th species is described with the canonical equation of adaptive dynamics theory (22):

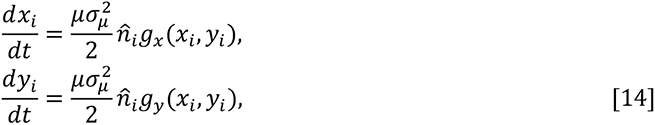

(see (30) for its extension for sexual reproduction), where *g_x_*(*x, y*) = *∂f*(*x, y*; **x, y**) / *∂x* and *g_y_*(*x, y*) = *∂f*(*x, y*; **x, y**)/*∂y* = *β* describe fitness gradients at (*x,y*) along the *x-* and *y*-directions, respectively. Under the isotropic mutation assumed here, the directional evolution described by Eq. 14 is always orthogonal to the adaptation front. From Eq. 14, we derive the velocity of adaptation front at each position (*x, y*) = (*x, h*(*x, t*)) on it (i.e., *f*(*x, y*; **x, y**) = *f*(*x, h*(*x, t*); **x, y**) = 0) as a vector (*u_x_*(*x, y*), *u_y_*(*x, y*)) with

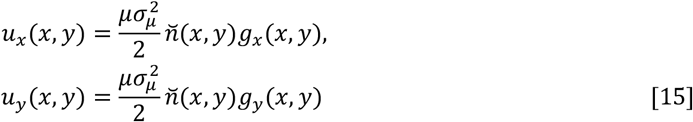

(derived from 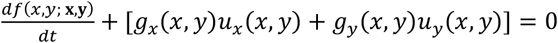 with 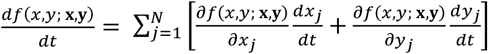 and *u_x_*(*x, y*) / *u_y_*(*x, y*) = *g_x_*(*x, y*) / *g_y_*(*x, y*) in SI Appendix, section 10A), where 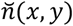 is a weighted sum of coexisting species’ population sizes, given by

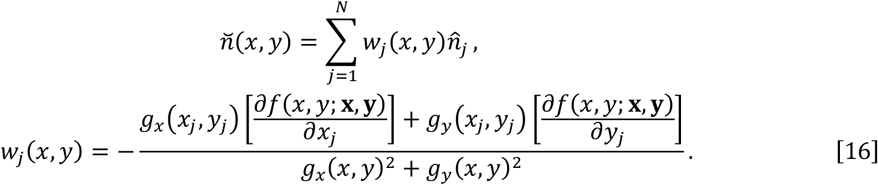

Here, *w_i_*(*x_j_, y_j_*) is equal to 1 for *i* = *j* and 0 otherwise, resulting in 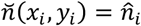 for all *i* = 1, …, *N*. Thus, Eq. 15 for (*x, y*) = (*x_i_, y_i_*) is equivalent to Eq. 14.

For a mutant (*x′*, *y′*) located close to the adaptation front *y* = *h*(*x, t*), we can expand its invasion fitness *f*(*x′*, *y′*; **x, y**) around (*x′*, *h*(*x′*, t)) with respect to *y′* as

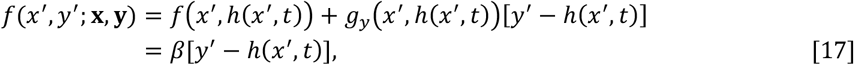

where *f*(*x′*, *h*(*x′*, *t*)) = 0 and *g_y_*(*x′*, *h*(*x′*, *t*)) = *β* are used (derived from Eq. 4). Hence, the fitness gradients along the *x*- and *y*-directions at each position (*x*, *y*) = (*x*, *h*(*x*, *t*)) on the adaptation front are expressed as

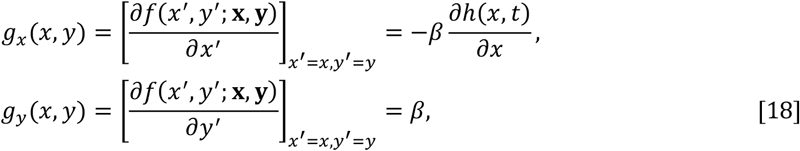

which, upon substitution into Eq.15, gives

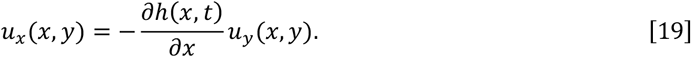

Additionally, the following geometrical relationship always holds:

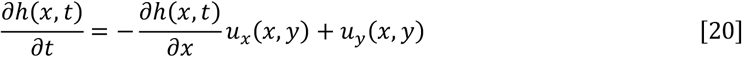

(derived from *h*(*x, t*) + Δ*y* ≃ *h*(*x* + Δ*x, t* + Δ*t*) for Δ*x* = *u_x_*(*x, h*(*x, t*))Δ*t* and Δ*y* = *u_y_*(*x, h*(*x, t*))Δ*t* for small Δ*t*, in SI Appendix, section 10C). Substitution of Eqs. 15 and 19 into Eq. 20 gives

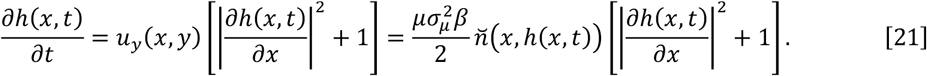

Since analytical treatment of 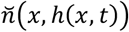 is difficult, we approximate Eq. 21 into a form without 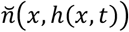, as follows. Under assumptions of *σ_α_* ≪ *σ_K_* and *β* ≪ 1, we can expect that species are densely and uniformly packed along the *x*-axis through the evolutionary dynamics, so that the expected density of species that compete with each other around niche *x* is kept approximately constant, denoted by 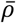. In this case, the expected dynamics of the adaptation front is approximately given by Eq. 21 with substitution of

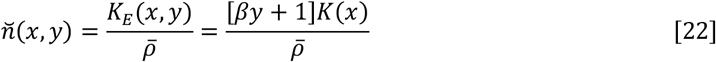

(see SI Appendix, section 10D for the derivation), where *K_E_*(*x, y*) = [*βy* + 1]*K*(*x*) is the equilibrium population size for a species of phenotype (*x, y*) in the absence of competitors, referred to as the effective carrying capacity. 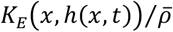 gives the expected population size for a species occupying niche *x* (with competitors) at time *τ*. 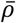 is derived as 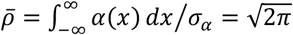 for the Gaussian competition kernel defined by Eq. 2 (see SI Appendix, 10D). Note that different shapes of competition kernel may result in different values for 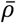.

### Time rescaling

To facilitate the analysis, we introduce a rescaled time *τ* = *h*(0, *t*) – *h*(0, 0) so that its unit progress corresponds to the unit progress of adaptation front at *x* = 0 along the *y*-direction. We also introduce the corresponding notation for the adaptation front, 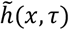, that satisfies 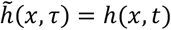. Then, provided that the front top is located at *x* = 0 for sufficiently large *τ*, so that 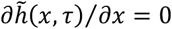 holds for *x* = 0, we see that

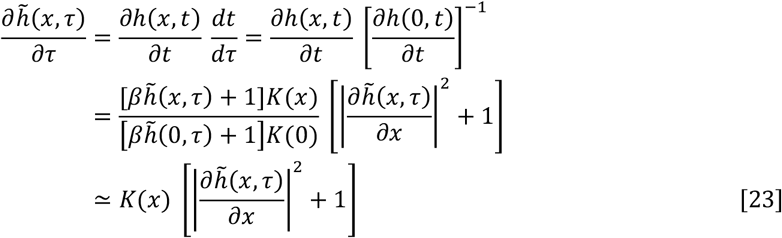

for finite |*x*|. Note that the approximation of the second row into the third row indicates that the ratio of the expected population size for a species occupying niche *x* to that for a species occupying *x* = 0 is approximately given by *K*(*x*). For notational simplicity, we rename 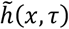 to *h*(*x, τ*) in this equation, which results in Eq. 5 in the main text.

Since the progress of the front top in the simulated evolution had stochastic noises. We calculated the rescaled time *τ* as the progress of the community average for *y*, i.e., 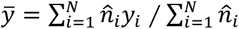, because it differed little from the value for *y* at the front top and had smaller stochastic noises than the front top.

### Adaptation horizon

From *∂h*(*x, τ*)/*∂τ* = *∂h*(0, *τ*)/*∂τ* = 1 and from *h*(*x, τ*) = *h*(0, *τ*) + *H*(*x*) for sufficiently large *τ*, we see that

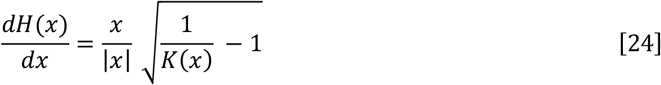

Therefore, *H*(*x*) is obtained by integrating Eq. 24 from the optimum niche *x* = 0 to niche *x*, resulting in Eq. (6). When average mutation sizes along the *x*- and *y*-directions are given by *σ_μxx_* and *σ_μyy_*, respectively, the rescaled adaptation front equation is given by

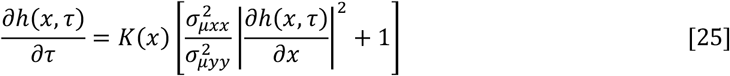

(see SI Appendix, section 8E for the derivation), from which the adaptation horizon is derived as 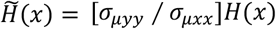 with *H*(*x*) given by Eq. 6.

### Directional evolution

Since Eqs. 19 and 20 trivially hold even when *t* is replaced with *τ*, we obtain from these equations and Eq. 5 the expected directional evolution for a species occupying niche *x* along the *x*- and *y*-directions in *τ*, denoted respectively by *v_x_*(*x, τ*) and *v_y_*(*x, τ*), as

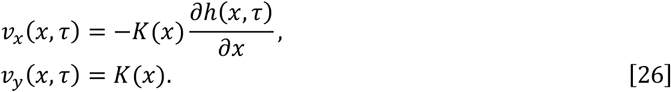

We obtain from Eqs. 24 and 26 the expected directional evolution for *τ* → ∞, i.e., Eq. 7.

### Net diversification rate

By introducing an expected species density 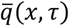 (i.e., the expected number of species existing within a unit niche range) around *x* at time *τ*, we derived that the following equation, known as the continuity equation in fluid dynamics (45), approximately holds,

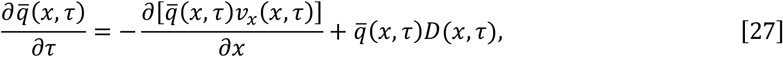

(SI Appendix, section 10E for the derivation), where the first and second terms on the right-hand side describe the divergence (through directional evolution) and source/sink (through branching/extinction), respectively. *D*(*x, τ*) represents the net diversification rate around *x* (i.e., the expected number of branching subtracted by that of extinction around *x* at time *τ* per unit species density and per unit time. Under uniform species packing kept through the evolutionary dynamics, corresponding to the constantness of the expected competing species density (i.e., 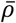), we can expect that 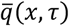 is also constant along *x* and *τ*, under the constant shape of the competition kernel defined by Eq. 2 (i.e., 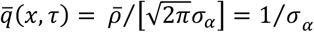 is derived in SI Appendix, section 10D). Then, Eq. 27 with 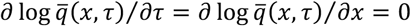 for *τ* → ∞ gives Eq. 10 in the main text.

### Divergence time

We consider a species of phenotype (*x*_1_, *y*_1_) = (*x*(*τ*_1_), *y*(*τ*_1_)) that exists at present *τ* = *τ*_1_ and shares the common ancestral phenotype (*x*_0_, *y*_0_) = (*x*(*τ*_0_), *y*(*τ*_0_)) at time *τ* = *τ*_0_ with its closest relative existing at *τ* = *τ*_1_. Note that *y*_1_ = *h*(*x*_1_, *τ*_1_) and *y*_0_ = *h*(*x*_0_, *τ*_0_) hold. We assume *x*_0_ > 0 without loss of generality. Provided that the adaptation front is given by *h*(*x, τ*) = *h*(0, *τ*) + *H*(*x*) = *h*(0, *τ*) – |*H*(*x*)|, we transform the divergence time for the focal species at time *τ* = *τ*_1_, denoted by *d_τ_* (*τ*_1_) = *τ*_1_ – *τ*_0_, as

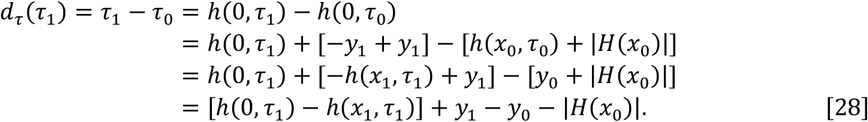

Since the expected evolutionary trajectory starting from (*x*_0_, *y*_0_) is given by Eq. 7, the following relationships hold (on average):

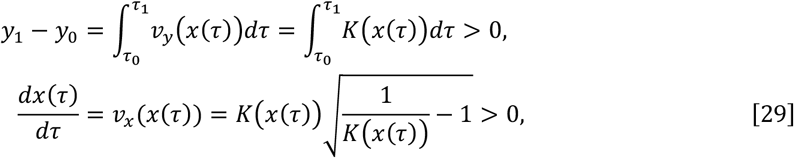

from which we see that

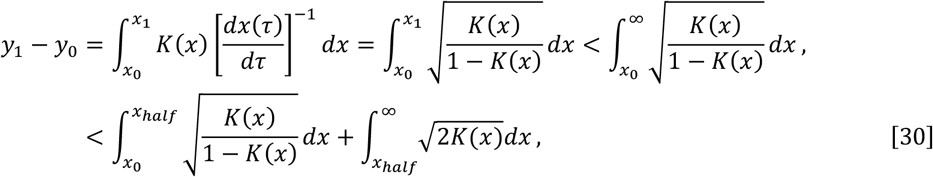

where *x_half_* satisfies *K*(*x_half_*) = 1/2 and *x_half_* > 0 (or 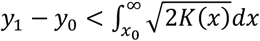 for *x*_0_ > *x_half_*). Thus, provided that 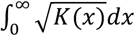 is finite, we get

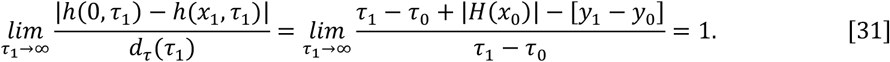

Additionally, from *h*(*x*_1_, *τ*_1_) = *h*(0, *τ*_1_) – |*H*(*x*_1_|, we see that |*h*(0, *τ*_1_) – *h*(*x*_1_, *τ*_1_)| = |*H*(*x*_1_)|. Therefore, we expect that Eq. 12 approximately holds for sufficiently large [*τ*_1_ – *τ*_0_]. The above derivation also applies for a clade consisting of species similar to each other such that their trajectories are approximately described with their average phenotype’s trajectory from (*x*_0_, *y*_0_) = (*x*(*τ*_0_), *y*(*τ*_0_)) to (*x*_1_, *y*_1_) = (*x*(*τ*_1_), *y*(*τ*_1_)).

