## Supplementary material for "The adaptation front equation explains diversification hotspots and living-fossilization": SI Appendix

**This PDF file includes:**

- SI Appendix
- Figs. S1 to S17

Throughout the text, we use italic for denoting scalars, bold lower case for column vectors, and bold upper case for matrices.

### 1 Condition for evolutionary branching line

We consider evolutionary dynamics starting from a single ancestral species of an arbitrary phenotype $(x, y)$ with $y \geqslant 0$. In this case, the invasion fitness for a mutant $(x', y')$, defined by Eq. 4 in the main text, is simplified into

$$f(x', y'; x, y) \;=\; \beta y' + 1 - \frac{\alpha(x'-x)[\beta y + 1]K(x)}{K(x')}. \qquad \text{[S1.1]}$$

The fitness gradients for $(x, y)$ are derived as

$$\begin{aligned} g_x(x,y) &= \left[\frac{\partial f(x',y';x,y)}{\partial x'}\right]_{x'=x,y'=y} = -\frac{[\beta y+1]x}{\sigma_K^2}, \\ g_y(x,y) &= \left[\frac{\partial f(x',y';x,y)}{\partial y'}\right]_{x'=x,y'=y} = \beta. \end{aligned} \qquad \text{[S1.2]}$$

We see that $g_x(x, y)$ is equal to zero for $x = 0$, positive for $x < 0$, and negative for $x > 0$. Hence, this species directionally evolves in trait $x$ toward $x = 0$, i.e., the line $x = 0$ is convergence stable along the $x$-axis. At the same time, this species evolves in trait $y$ toward larger values under the constant fitness gradient, $\beta$. To analyze the likelihood of evolutionary branching when the species comes close to the convergence stable line, $x = 0$, we calculate the first derivatives of $g_x(x, y)$ and $g_y(x, y)$ (with respect to $x$ and $y$) and the second derivatives of the invasion fitness (with respect to $x'$ and $y'$) for a species of phenotype $(0, y)$:

$$\begin{pmatrix} g_{xx} & g_{xy} \\ g_{yx} & g_{yy} \end{pmatrix} = \begin{pmatrix} \frac{\partial g_x(x,y)}{\partial x} & \frac{\partial g_x(x,y)}{\partial y} \\ \frac{\partial g_y(x,y)}{\partial x} & \frac{\partial g_y(x,y)}{\partial y} \end{pmatrix}_{x=0} = \begin{pmatrix} -\frac{[\beta y+1]}{\sigma_K^2} & 0 \\ 0 & 0 \end{pmatrix}, \qquad \text{[S1.3]}$$

$$\begin{aligned} \begin{pmatrix} f_{x'x'} & f_{x'y'} \\ f_{x'y'} & f_{y'y'} \end{pmatrix} &= \begin{pmatrix} \frac{\partial^2 f(x',y';x,y)}{\partial x'^2} & \frac{\partial^2 f(x',y';x,y)}{\partial x' \partial y'} \\ \frac{\partial^2 f(x',y';x,y)}{\partial x' \partial y'} & \frac{\partial^2 f(x',y';x,y)}{\partial y'^2} \end{pmatrix}_{x'=x=0,y'=y} \\ &= \begin{pmatrix} [\beta y+1]\left[\frac{1}{\sigma_\alpha^2} - \frac{1}{\sigma_K^2}\right] & 0 \\ 0 & 0 \end{pmatrix}. \end{aligned} \qquad \text{[S1.4]}$$

According to (1), evolutionary branching in the neighborhood of the convergence stable line, $x = 0$, is highly expected, if the following two conditions are satisfied: (i) the fitness function is significantly less sensitive to trait $y$ than to trait $x$, satisfying

$$\frac{|g_y(0,y)| + |g_{xy}| + |g_{yx}| + |g_{yy}| + |f_{x'y'}| + |f_{y'y'}|}{|g_x(0,y)| + |g_{xx}| + |f_{x'x'}|} \;=\; O(\sigma_\mu), \qquad \text{[S1.5]}$$

and (ii) a species of phenotype $(0, y)$ experiences a sufficiently strong disruptive selection along the $x$-direction (measured by $\sigma_\mu^2 f_{x'x'}$) in comparison with directional selection along the $y$-direction (measured by $\sigma_\mu |g_y(0, y)|$), satisfying

$$\frac{\sigma_\mu^2 f_{x'x'}}{\sigma_\mu |g_y(0, y)|} \quad > \quad \sqrt{2}. \qquad \text{[S1.6]}$$

By exploiting Eqs. S1.3 and S1.4, we obtain a sufficient condition for Eqs. S1.5 and S1.6 as

$$\frac{\sigma_\mu}{\beta\sqrt{2}} \left[ \frac{1}{\sigma_\alpha^2} - \frac{1}{\sigma_K^2} \right] \quad > \quad 1. \qquad \text{[S1.7]}$$

### 2 Simulation algorithm

#### 2.A Asexual reproduction

All evolutionary dynamics shown in figures in the main text were simulated with the same algorithm called the oligomorphic stochastic model (OSM) (2; 1; 3) that assumes asexual reproduction. The OSM in (3) is a slightly simplified version of the original OSM (2). The simulation in our present analysis followed (3). In the OSM in (3), the probability density for a mutant emergence with phenotype $(x', y')$ is proportional to $\mu \sum_{j=1}^{N} \xi(x', y'; x_j, y_j) \hat{n}_j$, where $\xi(x', y'; x_j, y_j)$ is a mutation distribution describing the probability density distribution for emergence of a mutant $(x', y')$ from its parental phenotype $(x_j, y_j)$ when a mutant emerges. The waiting time for a mutant emergence follows the exponential distribution with its expected value $1/\left[\mu \sum_{j=1}^{N} \hat{n}_j\right]$. The probability of successful invasion is given by the invasion fitness of the mutant (4). When a mutant has successfully invaded into the system, the next population-dynamical equilibrium is calculated by time integration of the equation for population dynamics (e.g., Eq. 1 in the main text for Fig. 1-4), with the initial population size for the mutant being set at a positive constant $\varepsilon_{\text{new}}$. During the population dynamics, a resident phenotype (or the mutant) is removed when its population size becomes lower than an extinction threshold $\varepsilon_{\text{ext}}$ (see (3) for details). $\varepsilon_{\text{ext}} = 1.0 \times 10^{-6}$ and $\varepsilon_{\text{new}} = 10\varepsilon_{\text{ext}}$ were used for all simulations with the OSM shown in this paper.

Regarding the mutation distribution, the isotropic mutation (with average mutation size $\sigma_\mu$) was described with the bivariate Gaussian distribution

$$\xi(x', y'; x, y) \quad = \quad \frac{1}{2\pi\sigma_\mu^2} \exp\left( -\frac{[x' - x]^2 + [y' - y]^2}{2\sigma_\mu^2} \right). \qquad \text{[S2.1]}$$

The "separate mutation," which occurs in traits $x$ and $y$ separately (with mutation rates $\mu_x$ and $\mu_y$, and average mutation sizes $\sigma_{\mu x}$ and $\sigma_{\mu y}$ ), was described with

$$\begin{aligned} \xi(x', y'; x, y) \quad = \quad & \frac{\mu_x}{\mu} \delta(y' - y) \frac{1}{\sqrt{2\pi}\sigma_{\mu x}} \exp\left( -\frac{[x' - x]^2}{2\sigma_{\mu x}^2} \right) \\ & + \frac{\mu_y}{\mu} \delta(x' - x) \frac{1}{\sqrt{2\pi}\sigma_{\mu y}} \exp\left( -\frac{[y' - y]^2}{2\sigma_{\mu y}^2} \right) \end{aligned} \qquad \text{[S2.2]}$$

with $\mu = \mu_x + \mu_y$, where $\delta(z)$ is the Dirac's delta function (i.e., $\delta(z) = 0$ for $z \neq 0$, $\delta(z) = \infty$ for $z = 0$, and $\int_{-\infty}^{\infty} \delta(z) \mathrm{dz} = 1$).

The simulation was conducted with software R (version 3.4.4) and its package deSolve-1.28 (5).

#### 2.B Sexual reproduction

We simulated the evolutionary dynamics under sexual reproduction and small but nonrare mutation (Figs. S1 and S3). This algorithm considers male and female individuals, diploid inheritance, and quantitative characters coded for by multilocus genetics. Specifically, $L_x$ and $L_y$ loci are considered for traits $x$ and $y$, respectively, with integer allelic values and with the value of $x$ and $y$ given by the averages of allelic values across loci. For recombination, $C_x$ and $C_y$ linkage clusters are considered for $x$ and $y$, respectively, with no recombination within such a cluster and with free recombination between the clusters. Individual birth and death rates are defined so that the expected population dynamics is described with Eq. 1 in the main text. Specifically, the $k$th female individual produces offspring individuals at rate $b_k = 2[\beta y_k + 1]$, and dies at rate $\sum_{j=1}^{N_I} \alpha(x_k, x_j)/K(x_k)$, where $N_I$ describes the total number of individuals in the system. The male individuals do not produce offsprings, and the $l$th male individual dies at rate $\sum_{j=1}^{N_I} \alpha(x_l, x_j)/K(x_l)$. For offspring production, a female individual $k$ chooses a male partner $l$ for mating with probability

$$\tilde{P}_{kl} = \frac{P_{kl} b_l}{\sum_{l'} P_{kl'} b_{l'}}, \qquad \text{[S2.3]}$$

where $b_l = [\beta y_l + 1]$ denotes the potential of male individual $l$ for contribution to offspring production. $P_{kl}$ describes the mating likelihood between the female $k$ and the male $l$, depending on a $L_m$-dimensional display trait $\mathbf{m}_l = (m_{l,1}, \ldots, m_{l,L_m})$ in male $l$ and on a corresponding preference trait $\mathbf{p}_k = (p_{k,1}, \ldots, p_{k,L_m})$ in female $k$. Each of $m_{l,1}, \ldots, m_{l,L_m}$ and $p_{k,1}, \ldots, p_{k,L_m}$ has a locus containing an integer value, and those loci are organized into $C_m$ linkage clusters. The mating likelihood $P_{kl}$ is then given by

$$P_{kl} = \exp\left(-\frac{d_{m,kl}^2}{2\sigma_{\text{mating},m}^2}\right) \exp\left(-\frac{d_{x,kl}^2}{2\sigma_{\text{mating},x}^2}\right), \qquad \text{[S2.4]}$$

with $d_{m,kl}^2 = |\mathbf{p}_k - \mathbf{m}_l|^2$ and $d_{x,kl}^2 = [x_k - x_l]^2$, so that $d_{m,kl}$ measures the distance between the preference trait of female $k$ and the display trait of male $l$, and $d_{x,kl}$ measures their distance in the niche trait, $x$. Hence, the mating probability is an increasing function with the similarity between a male's display trait and a female's preference trait, and between their niches. Although $\sigma_{\text{mating},x} = \infty$ is assumed in (2), we here assume $\sigma_{\text{mating},x} = \sigma_\alpha$ because accumulating empirical studies show that assortative mating depending on niche similarities may be a plausible and important factor for sympatric and parapatric speciation (6). Allelic mutations that increase or decrease allelic values by 1 (which happens with equal probability) occur with per locus probabilities of $\tilde{\mu}_x$, $\tilde{\mu}_y$, $\tilde{\mu}_m$, and $\tilde{\mu}_p$ for each birth of an individual with its sex assigned randomly. For the simulation, the above algorithm was implemented in C++ (5).

At each sampling time point in the simulation, species were identified by clustering individuals of similar phenotypes, so that the phenotypic difference between two individuals chosen from different clusters are always more than two mutational steps.

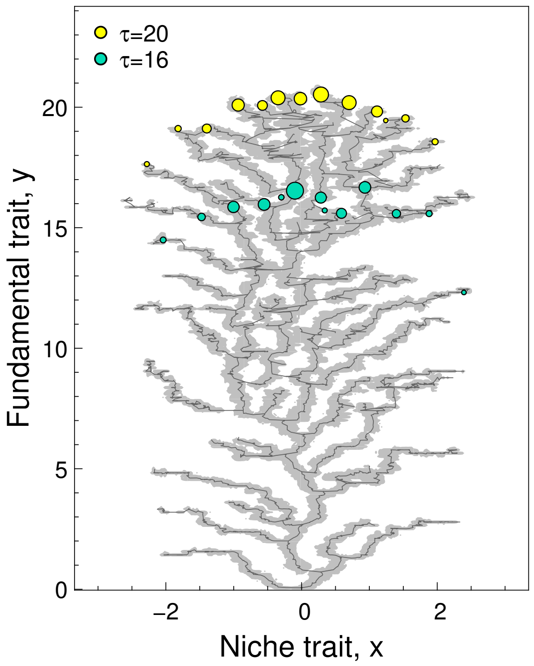

Fig. S1. Simulated evolution under sexual reproduction for the model defined by Eqs. 1-3 in the main text. The gray thin curves indicate trajectories of mean phenotypes of species. The light gray color indicates the region where at least one individual was observed in equally-spaced time samplings ($\Delta t = 100$) from $t = 0$ ($\tau = 0$) to $t = 17904.8$ ($\tau = 20$). Mean phenotypes of coexisting species at $\tau = 16$ and $\tau = 20$ are plotted with blue-green and yellow circles, respectively, where their population sizes are indicated with the sizes of circles. The original model parameters are $K_0 = 500$, $\sigma_K = 0.15$, $\sigma_\alpha = 0.04$, $\beta = 2$, $(L_x, L_y, L_m, L_p) = (120, 50, 20, 20)$, $(C_x, C_y, C_m, C_p) = (12, 5, 2, 2)$, $(\tilde{\mu}_x, \tilde{\mu}_y, \tilde{\mu}_m, \tilde{\mu}_p) = (2.5 \times 10^{-5}, 1.0 \times 10^{-5}, 5.0 \times 10^{-3}, 5.0 \times 10^{-3})$. $\mu_x$ and $\mu_y$ (mutation rates per unit population size) are given by $\mu_x = 2L_x\tilde{\mu}_x K_0$ and $\mu_y = 2L_y\tilde{\mu}_y K_0$, respectively, after population sizes are linearly scaled so that $K_0$ becomes to 1 . The initial ancestral phenotype was $(x, y, \mathbf{m}, \mathbf{p}) = (0, 0, \mathbf{0}, \mathbf{0})$. To facilitate comparison with the simulated evolution under asexual reproduction (Figs. 1-3 in the main text), the the $x$- and $y$-axes were rescaled by multiplying $\sigma_K^{-1}$ and $\sigma_K^{-1}\sqrt{\mu_x \Delta x_\mu^2 / \mu_y \Delta y_\mu^2}$, respectively, with $\Delta x_\mu = 1/[2L_x]$ and $\Delta y_\mu = 1/[2L_y]$ , so that $\sigma_K = 1$ and isotropic mutation (in its effect on the canonical equation) were attained in the rescaled trait space, where $\beta = 0.29$, and $\sigma_\alpha = 0.27$.

### 3 Robustness: sexual reproduction

To facilitate comparison of simulation results among different eco-evolutionary settings, we present these results in the same format as shown in Fig. S2, which shows the simulated evolution for the same parameter set with that shown in Figs. 1-3 in the main text in a more informative way (but the initial phenotype for Fig. S2 is $(x_a, y_a) = (0, 0)$, differently from $(x_a, y_a) = (-0.2, 0)$ for Figs. 1-3). The prediction (Eqs. 6-12 in the main text) and simulated evolution under sexual reproduction is shown in Fig. S3 with the same format.

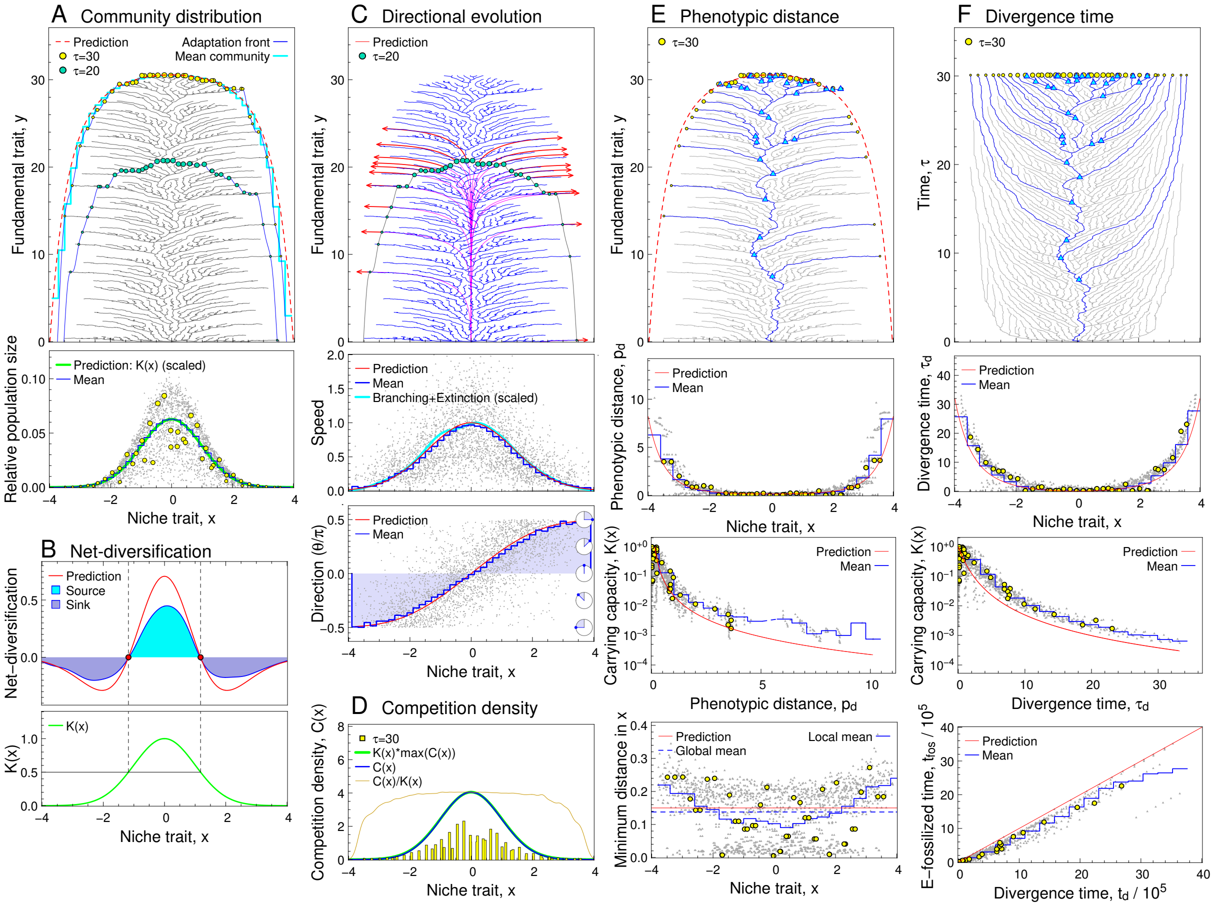

Fig. S2. Simulated evolution for the same eco-evolutionary setting with that shown in Figs. 1-3 in the main text, plotted in a more informative format.

#### Caption for Fig. S2

- A : Community distribution. Plottings are the same as in Fig. 1A in the main text, except that the cyan histogram shows the time-averaged shape of community distribution $y = y_{\rm com}(x) + \bar{y}_{\tau=30}$, where $y_{\rm com}(x)$ is the time-averaged value for $y - \bar{y}$ for each $x$ among coexisting species, where $\bar{y} = \sum_{j=1}^{N} y_j \hat{n}_j / \sum_{j=1}^{N} \hat{n}_j$ (sampled every 100 mutant invasions until $\tau = 60$). $\bar{y}_{\tau=30}$ describes the values for $\bar{y}$ at $\tau = 30$.

- B: Net diversification. Plottings are the same as in the middle and bottom panels of Fig. 2.

- C: Directional evolution. Plottings in the first and second panels are the same as in Fig 1B. The third panel shows directions of directional evolution of species occupying each $x$, in a manner analogous to the second panel. The red curve shows the prediction by Eq. 9 in the main text, i.e., $\theta = \tan^{-1}(v_y(x)/v_x(x))$ .

- D: Competition density distribution. The blue, green, and dark-yellow curves show $C(x) = \sum_{j=1}^{N} \alpha(x, x_j)\hat{n}_j$ (competition density distribution), $K(x)\max_x(C(x))$ and $C(x)/K(x)$, respectively, at $\tau = 30$. The yellow histogram shows the phenotype distribution along the $x$-axis at $\tau = 30$.

- E: Phenotypic distance. Plottings in the first and second panels are the same as in Fig. 3A. The third panel converts the plottings in the second panel for $x$ and $p_d$ into those for $p_d$ and $K(x)$, where the prediction curve (red) shows a set of points $(p_d, K(x))$ generated from a set of points $(p_d, x)$ on the prediction curve in the second panel. The bottom panel shows the minimum niche distance for each species from the others, in a manner analogous to the second panel. The global mean (blue dashed line) is compared with the prediction, $\sigma_\alpha = 0.15$ (red line).

- F: Divergence time. Plottings in the first and second panels are the same as in Fig. 3B. The third panel converts the plottings in the second panel for $x$ and $\tau_d$ into those for $\tau_d$ and $K(x)$. The bottom panel shows the relationship between $t_d$ and the "effectively-fossilized time," denoted by $t_{\mathrm{fos}}$. The $t_{\mathrm{fos}}$ was calculated as the non-rescaled divergence time $t_d$ for each species subtracted by the time period converted from evolutionary progress of the species in $y$ (based on the speed of adaptation front top). Specifically, $t_{\mathrm{foss},i}(t)$ for the $i$th species at $t$ is defined by $t_{\mathrm{foss},i}(t) = t_{d,i}(t) - \Delta t_h(y_{d,i}(t), y_i(t))$, where $t_{d,i}(t)$ describes the non-rescaled divergence time for the $i$-th species at $t$ from its closest relative, while $\Delta t_h(y_i(t), y_{d,i}(t))$ describes the non-rescaled time period taken for the adaptation front top, $y = h(0, t)$, to progress from $y = y_{d,i}(t)$ (i.e., trait $y$ for the $i$th species at its divergence time point $t - t_{d,i}$) to $y = y_i(t)$ (i.e., trait $y$ for the $i$th species at time point $t$).

The middle panel of Fig. S3C shows that evolution of the species occupying the outermost niches might not be the slowest, possibly due to the large effect of genetic drift caused by their small population sizes. However, the mean community distribution (cyan histogram in the top panel of Fig. S3A) shows that the outer most species are the least advanced in the fundamental trait, implying that the outer most species is the slowest in adaptive evolution.

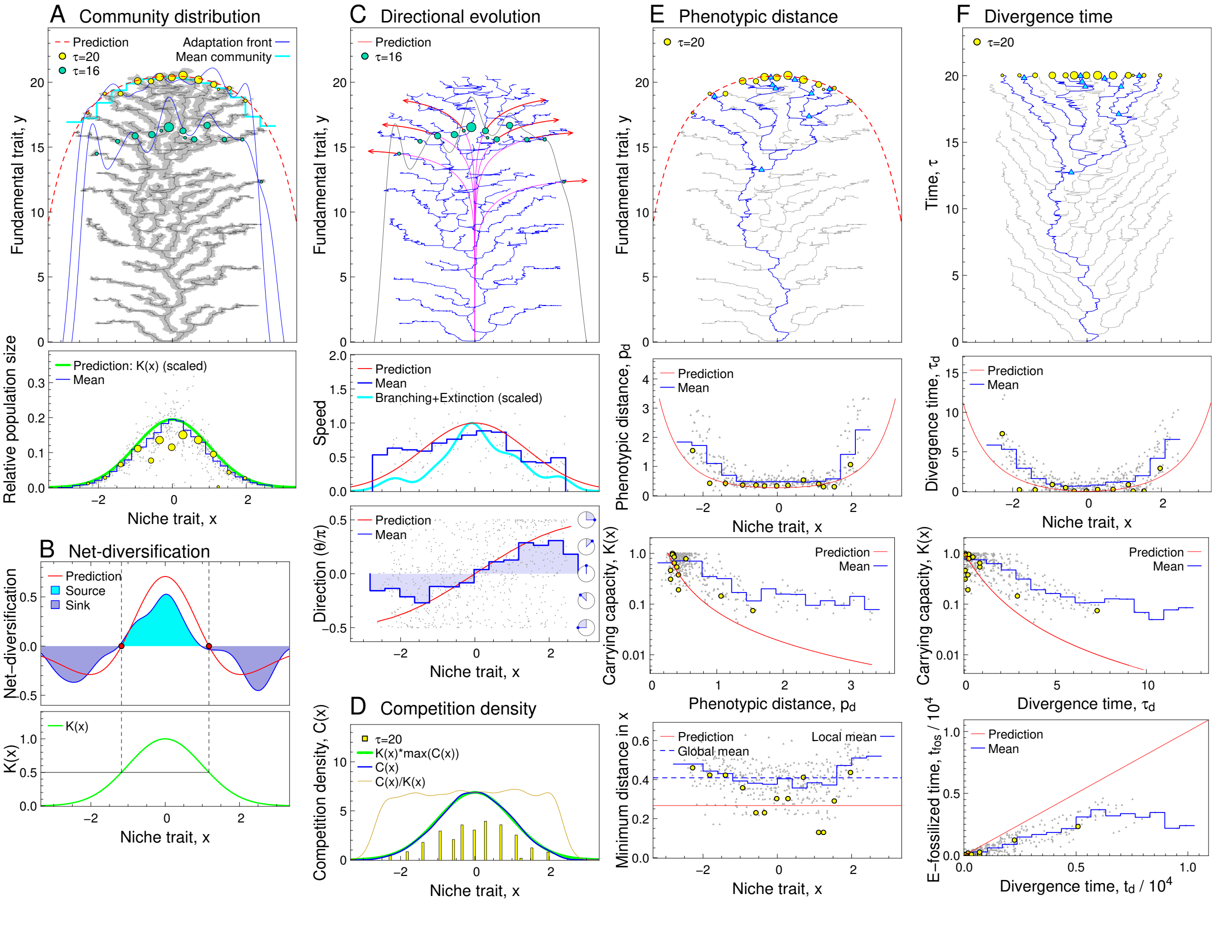

Fig. S3. Simulated evolution under sexual reproduction, plotted in the same manner as in Fig. S2. See SI Appendix, section 2B for the simulation algorithm and model parameters.

### 4 Robustness: Different carrying capacity distributions

#### 4.A Sharp-peaked and flat-topped distributions

We consider the carrying capacity distribution defined by

$$K(x) = \frac{1}{\nu^2 x^{2\phi} + 1} \qquad \text{[S4.1]}$$

with an integer parameter $\phi$ larger than 1 and a constant real number $\nu$. Note that $\phi = 1$ gives the Cauchy distribution. Substituting Eq. S4.1 into Eqs. 6-12 in the main text yields

$$\begin{aligned}
H(x) &= -\frac{\nu}{\phi+1}\frac{x}{|x|}|x^{\phi+1}|, \\
v_x(x) &= \frac{x}{|x|}\frac{\nu}{\nu^2 x^{2\phi}+1}|x^{\phi}|, \\
v_y(x) &= \frac{1}{\nu^2 x^{2\phi}+1}, \\
D(x) &= \left|\frac{2\nu\phi x^{\phi-1}}{\nu^2 x^{2\phi}+1}\right|\left[\frac{1}{\nu^2 x^{2\phi}+1} - \frac{1}{2}\right], \\
p_d(x) &= \sigma_\alpha\sqrt{\nu^2 x^{2\phi}+1}, \\
\tau_d(x) &= \frac{\nu}{\phi+1}|x^{\phi+1}|. \qquad \text{[S4.2]}
\end{aligned}$$

Figs. S4 an S5 show the predictions (Eq. S4.2) and simulated evolution under $\phi = 1$ (i.e., sharp-peaked distribution) and $\phi = 3$ (i.e., flat-topped distribution), respectively. Note that the net diversification rate for $\phi = 3$ in the simulated evolution shows a bimodal distribution (Fig. S5B), as predicted by $D(x)$ in Eq. S4.2 for $\phi = 3$.

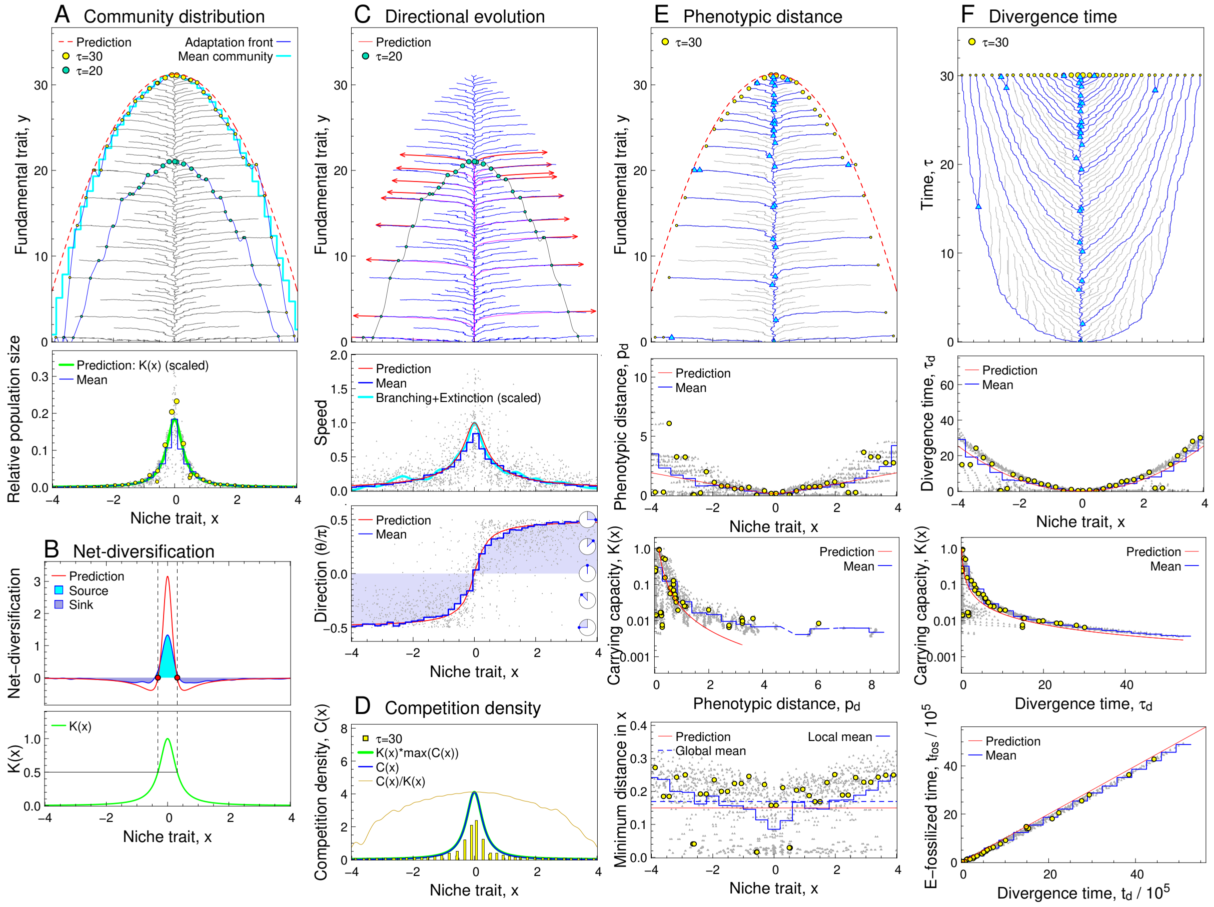

Fig. S4. Simulated evolution under sharp-peaked carrying capacity distribution defined by Eq. S4.1 with $\phi = 1$ and $\nu = 10$, plotted in the same manner as in Fig. S2. The other settings and parameters were the same with those for the simulated evolution shown in Fig. S2.

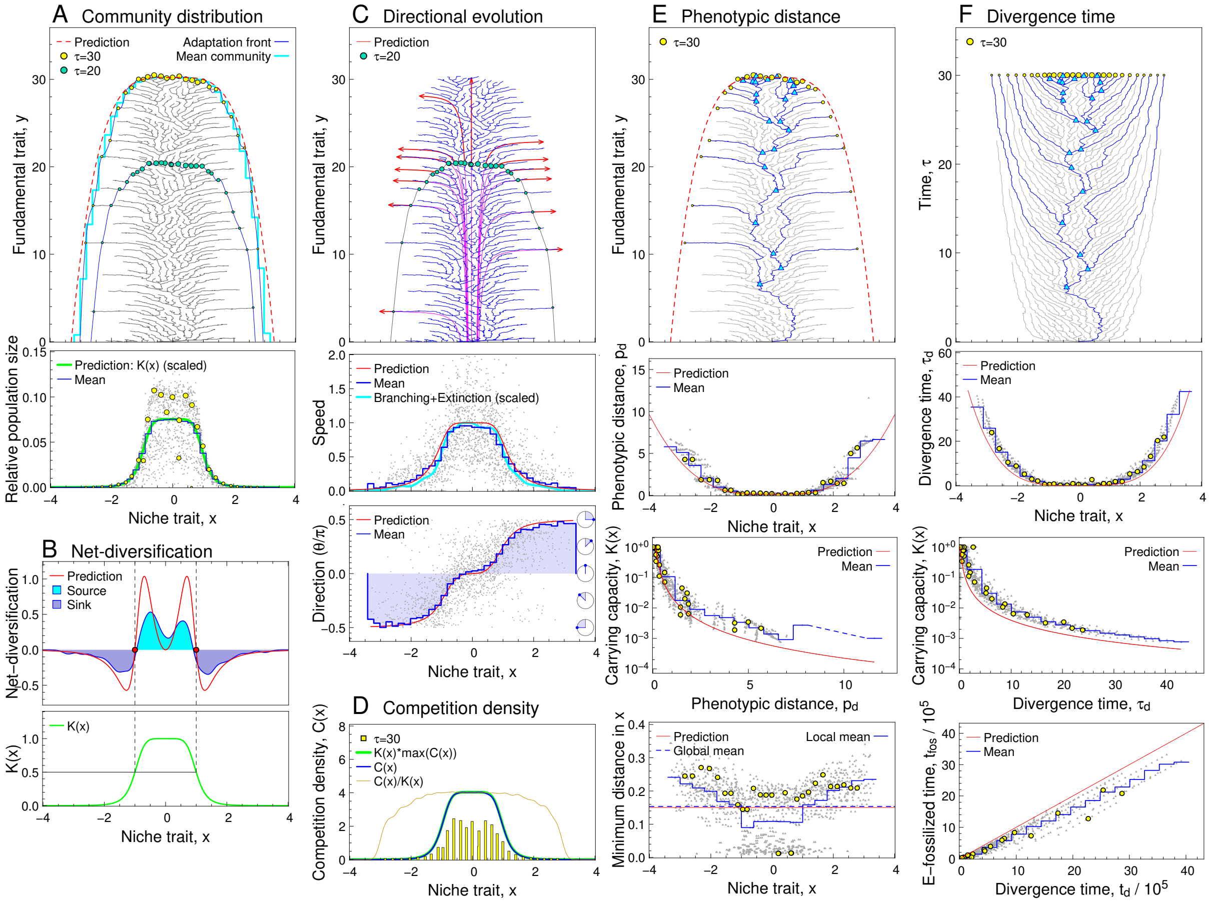

Fig. S5. Simulated evolution under flat-topped carrying capacity distribution defined by Eq. S4.1 with $\phi = 3$ and $\nu = 1$, plotted in the same manner as in Fig. S2. The other settings and parameters were the same with those for the simulated evolution shown in Fig. S2.

#### 4.B Bimodal distribution

We consider a bimodal carrying capacity distribution, defined by

$$K(x) = \nu_K \left[ \exp\left( -\frac{x^2}{2\sigma_K^2} \right) + w_K \exp\left( -\frac{[x - x_K]^2}{2\sigma_K^2} \right) \right] \quad \text{[S4.3]}$$

with $x_K = 2.0$, $w_K = 0.8$, and $\sigma_K = 0.6$, whereas $\nu_K$ is numerically chosen so that the maximum value of $K(x)$ is equal to 1. Note that the niche position that maximizes $K(x)$, denoted by $x = x^*$ is not equal to zero. We expect that $x = x^*$ attains the fastest progress of the adaptation front, resulting in the adaptation horizon given by

$$H(x) = -\frac{x - x^*}{|x - x^*|} \int_{x^*}^{x} \sqrt{\frac{1}{K(\tilde{x})} - 1} d\tilde{x}. \quad \text{[S4.4]}$$

Then by using this $H(x)$ (instead of Eq. 6 in the main text), we obtain $v_x(x)$, $v_y(x)$, $D(x)$, $p_d(x)$, and $\tau_d(x)$ from Eq. 7-12. Fig. S6 shows these predictions and the simulated evolution under $K(x)$ defined by Eq. S4.3.

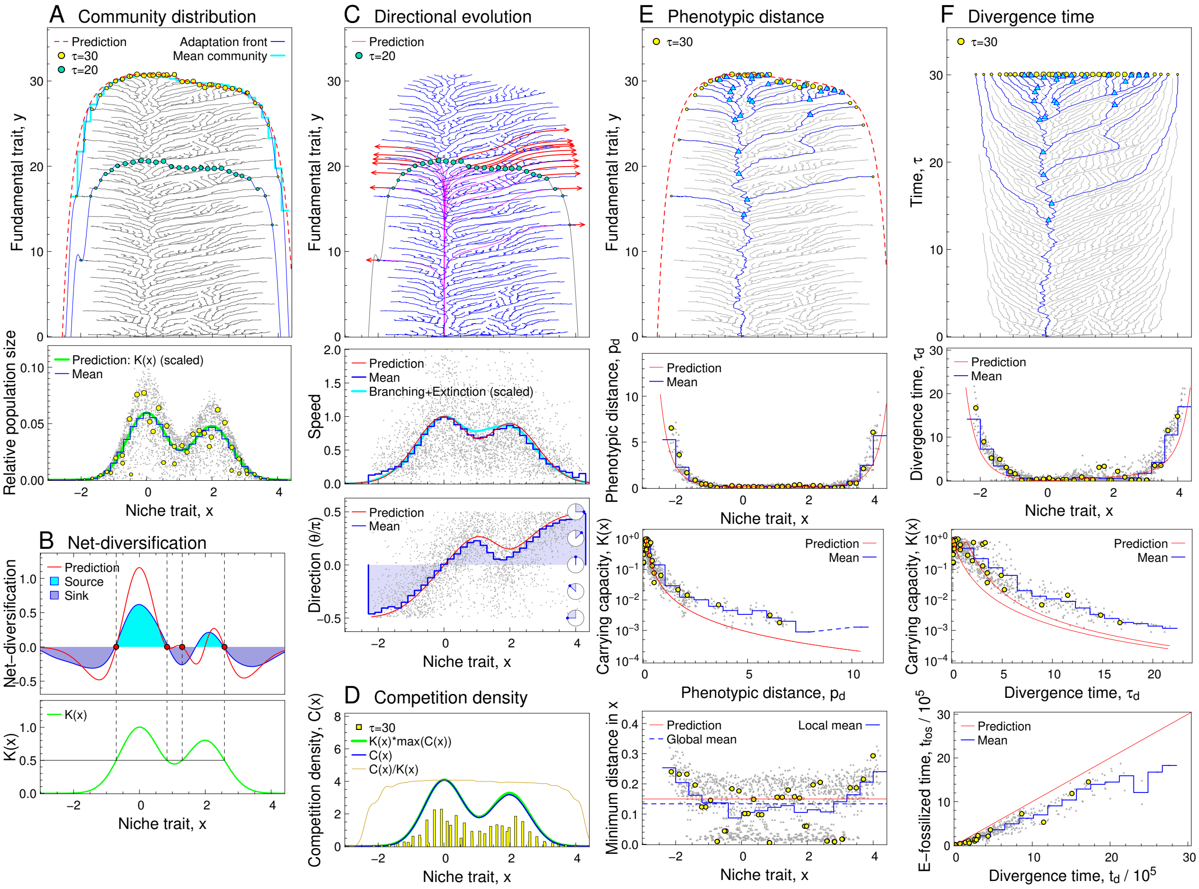

Fig. S6. Simulated evolution under bimodal carrying capacity distribution defined by Eq. S4.3, plotted in the same manner as in Fig. S2. The other settings and parameters were the same with those for the simulated evolution shown in Fig. S2.

### 5 Robustness: Dependency of fitness gradient for fundamental trait on niche trait

We assume that the fitness gradient in the $y$-direction is not a constant but a function of $x$, denoted by $\beta(x)$. For the case considered here, the general form of the rescaled adaptation front equation for two-dimensional trait spaces, Eq. S8.19, is simplified into

$$\begin{aligned} \frac{\partial h(x,\tau)}{\partial \tau} &= v_{hy}(x)\left[\left|\frac{\partial h(x,\tau)}{\partial x}\right| + 1\right], \\ v_{hy}(x) &= \lim_{t\to\infty} \frac{\beta(x)[\beta(x)h(x,t)+1]K(x)}{\beta(x^*)[\beta(x^*)h(x^*,t)+1]K(x^*)} = \frac{\beta(x)^2 K(x)}{\beta(x^*)^2 K(x^*)} \end{aligned} \quad \text{[S5.1]}$$

(by substitution of $\mu(x,t) = \mu$, $V_{\mu xx}(x,t) = V_{\mu yy}(x,t) = \sigma_\mu$, $K_E(x,t) = [\beta(x)h(x,t)+1]K(x)$, $g_y(x,t) = \beta(x)$, $\bar{\rho}(x,t) = \bar{\rho}$), provided that the front top is located at $x = x^*$ that maximizes $v_{hy}(x)$ for sufficiently large $t$. By using Eqs. S8.20-S8.22, we obtain

$$\begin{aligned} H(x) &= -\frac{x-x^*}{|x-x^*|}\int_{x^*}^{x}\sqrt{\frac{1}{v_{hy}(\tilde{x})}-1}d\tilde{x}, \\ v_x(x) &= \frac{x-x^*}{|x-x^*|}\sqrt{v_{hy}(x)[1-v_{hy}(x)]}, \\ v_y(x) &= v_{hy}(x), \\ D(x) &= \frac{dv_x(x)}{dx}, \\ p_d(x) &= \frac{\sigma_a}{\sqrt{v_{hy}(x)}}, \\ \tau_d(x) &= |H(x)|. \end{aligned} \qquad [S5.2]$$

Fig. S7 shows these predictions and the simulated evolution under $\beta(x) = \beta_0[1 + \exp(-[x - x_\beta]^2/[2\sigma_\beta^2])]$ with $\beta_0 = 0.1$, $x_\beta = 1.5$, and $\sigma_\beta = 0.55$.

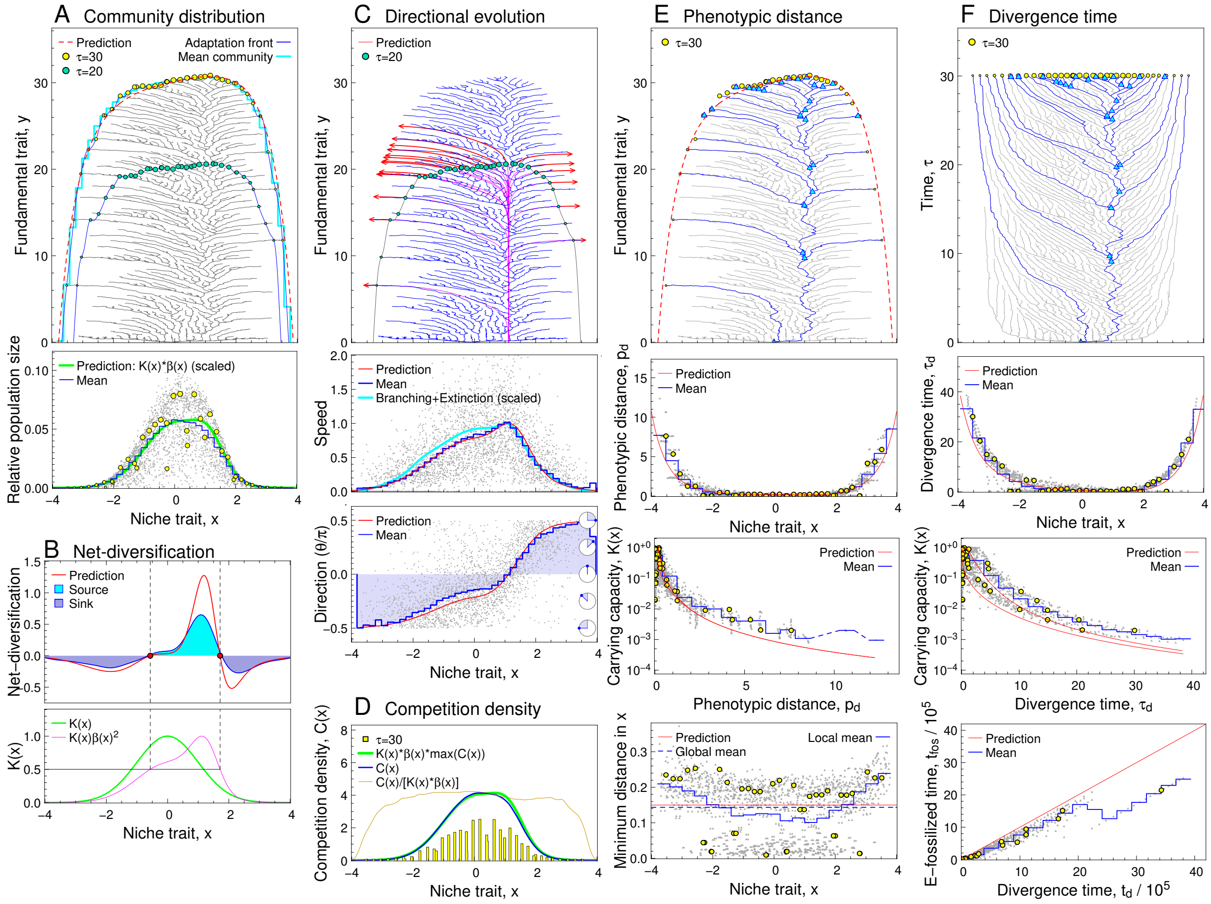

Fig. S7. Simulated evolution under fitness gradient along the $y$-direction depending on $x$, plotted in the same manner as in Fig. S2. The fitness gradient along the $y$-direction was defined by $\beta(x) = \beta_0[1 + \exp(-[x - x_\beta]^2/[2\sigma_\beta^2])]$ with $\beta_0 = 0.1$, $x_\beta = 1.5$, and $\sigma_\beta = 0.55$. The other settings and parameters were the same with those for the simulated evolution shown in Fig. S2.

### 6 Robustness: Different competition kernels

#### 6.A Platykurtic and leptokurtic kernels

The original competition kernel is defined as the Gaussian distribution, in Eq. 3 in the main text. We here consider a modified kernel defined by

$$\begin{aligned} \alpha(x) &= \tilde{\alpha}(x;\omega,\tilde{\sigma}_\alpha), \\ \tilde{\alpha}(x;\omega,\tilde{\sigma}_\alpha) &= \exp\left(-\frac{x^2}{2\tilde{\sigma}_\alpha^2}\right)\exp\left(\frac{1+|x/\tilde{\sigma}_\alpha|\omega^{-2}}{1+|x/\tilde{\sigma}_\alpha|}\right), \end{aligned} \qquad \text{[S6.1]}$$

where $\omega$ is a modification parameter such that $\omega > 1$ and $\omega < 1$ correspond to a leptokurtic kernel (curtosis $> 0$) and a platykurtic kernel (curtosis $< 0$), respectively (Fig. S8). The value for $\tilde{\sigma}_\alpha$ is numerically specified so that the standard deviation for $\tilde{\alpha}(x;\omega,\tilde{\sigma}_\alpha)$ with a given $\omega$ is kept equal to that for the original kernel, $\sigma_\alpha$.

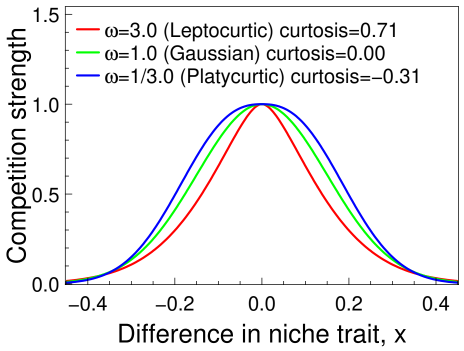

Fig. S8. Platykurtic and leptokurtic kernels.

Figs. S9 and S10 show the predictions (Eq. 6-12 in the main text) and the simulated evolution under the competition kernel defined by Eq. S6.1 with $\omega = 1.5$ (leptokurtic kernel) and $\omega = 1/1.5$ (platykurtic kernel), respectively.

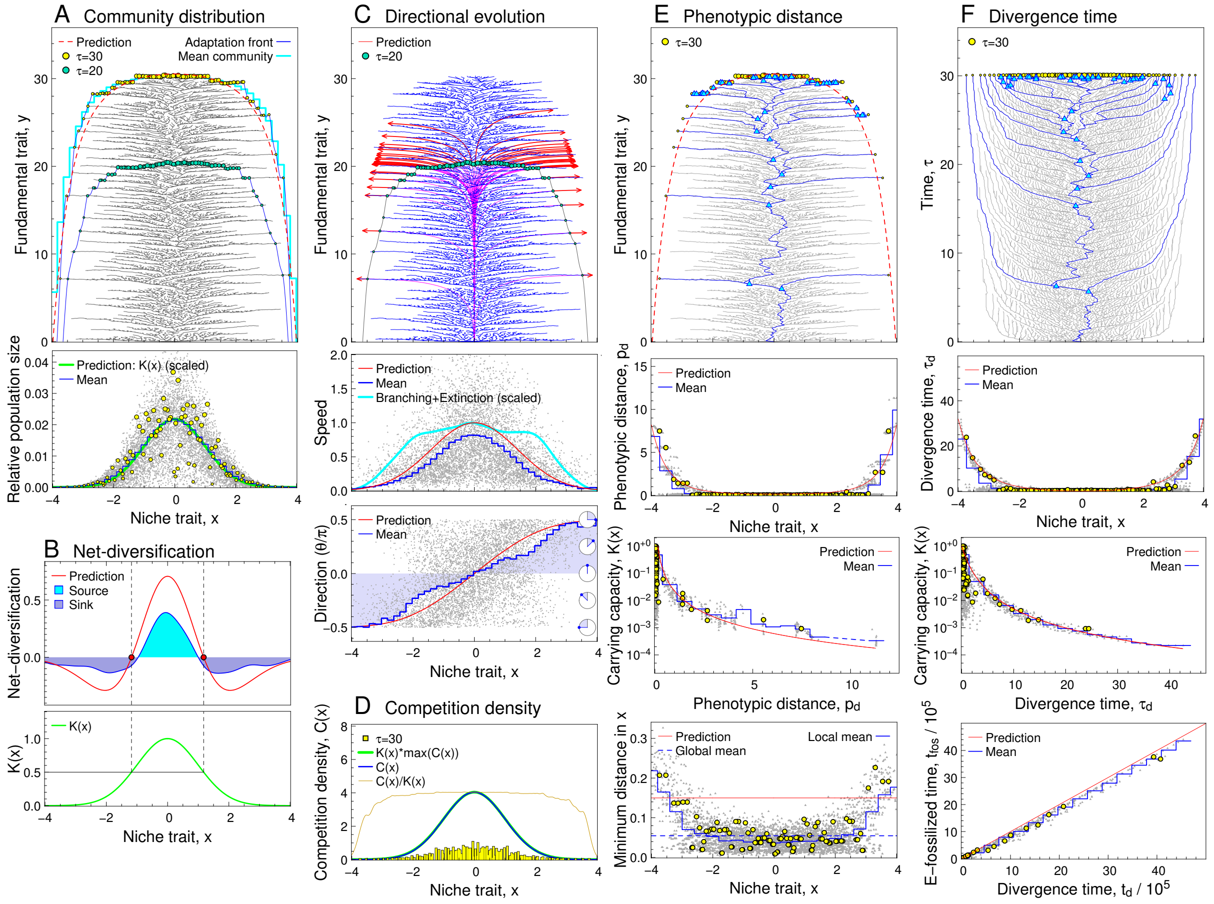

Fig. S9. Simulated evolution under leptokurtic competition kernel defined by Eq. S6.1 with $\omega = 1.5$, plotted in the same manner as in Fig. S2. The other parameters were the same with those for the simulated evolution shown in Fig. S2.

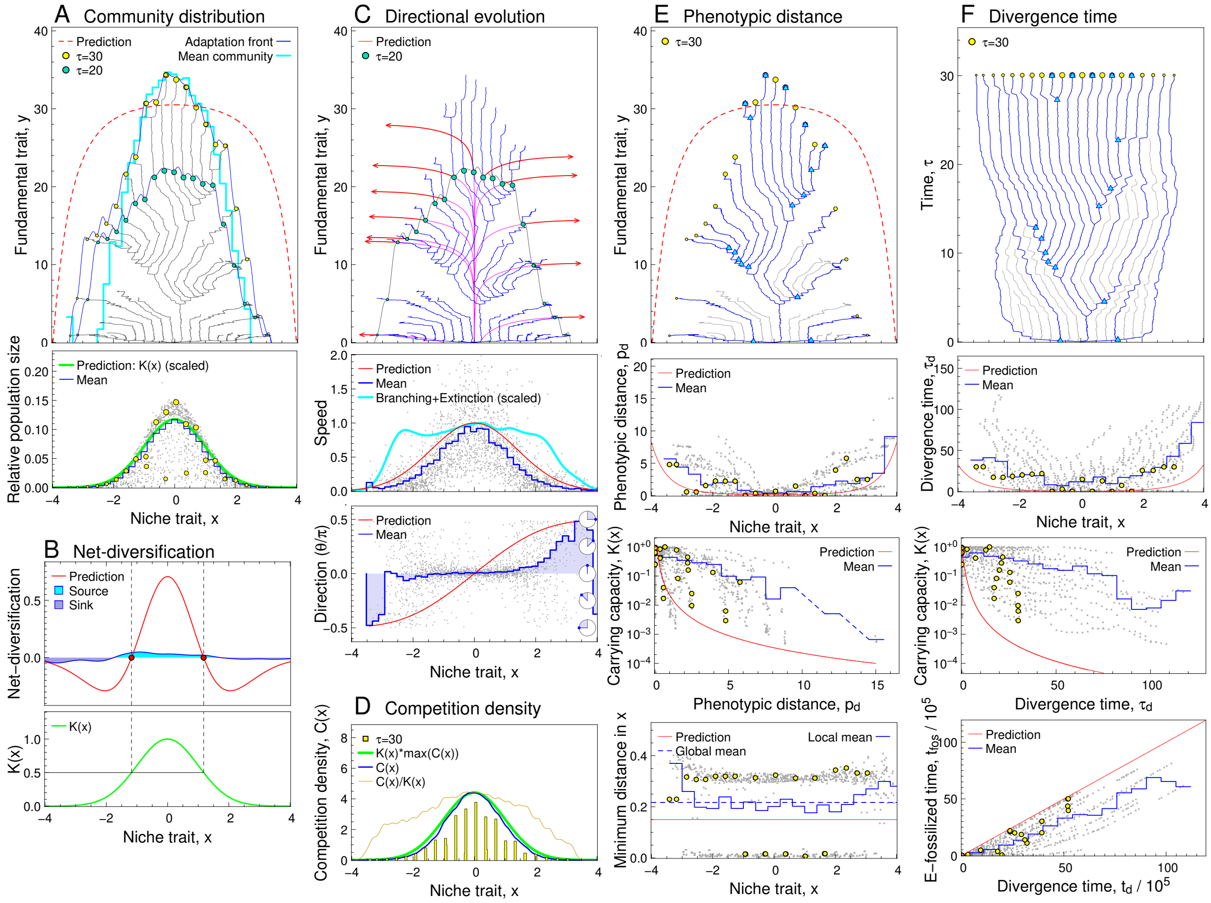

Fig. S10. Simulated evolution under platykurtic competition kernel defined by Eq. S6.1 with $\omega = 1/1.5$, plotted in the same manner as in Fig. S2. The other parameters were the same with those for the simulated evolution shown in Fig. S2.

#### 6.B Asymmetric kernel

We consider an asymmetric competition kernel, defined by

$$\alpha(x) \;=\; \exp\left(-\frac{m_\alpha^2}{2\sigma_\alpha^2}\right)\exp\left(-\frac{[x-m_\alpha]^2}{2\sigma_\alpha^2}\right) \qquad \text{[S6.2]}$$

with $m_\alpha = \sigma_\alpha/2$, following (7). Fig. S11 shows the predictions (Eqs. 6-12 in the main text) and the simulated evolution under this asymmetric competition kernel.

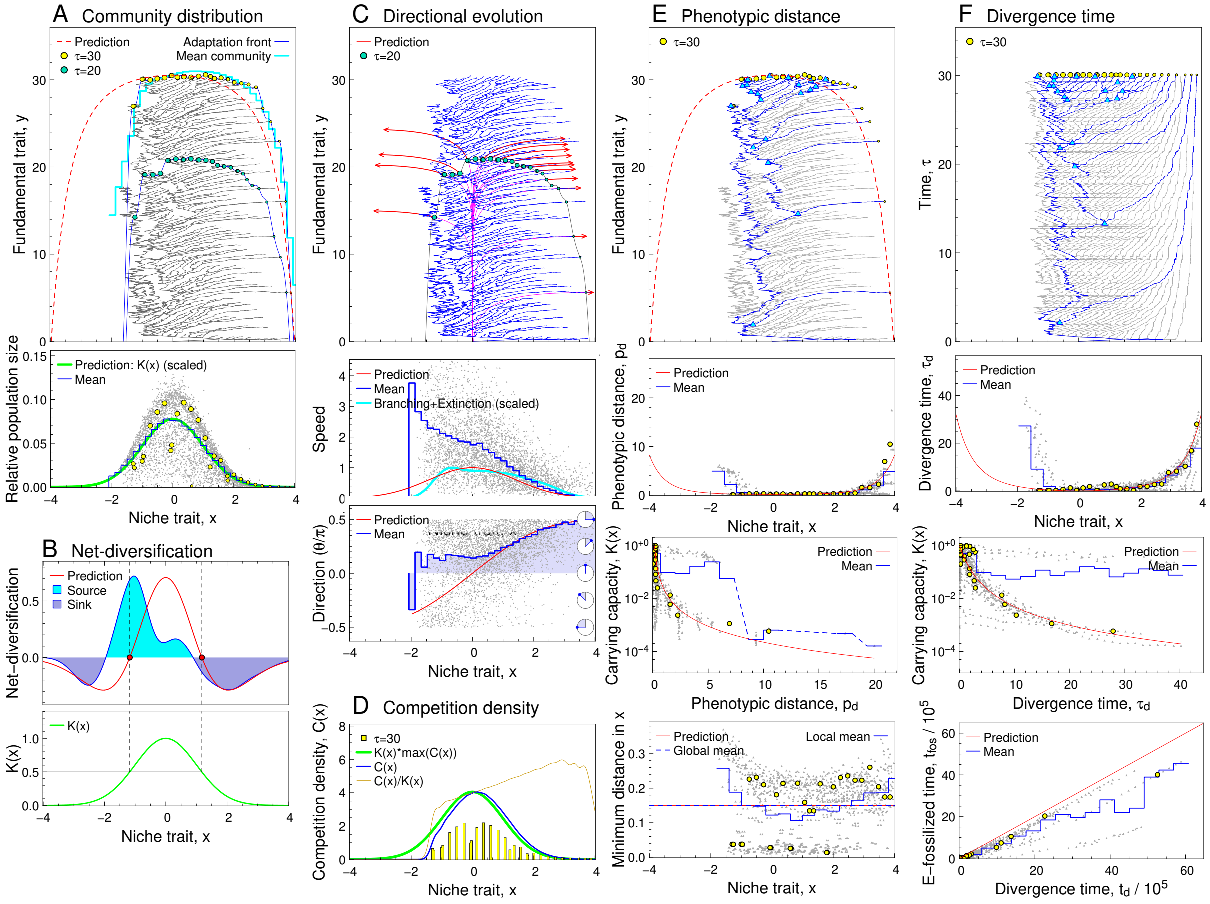

Fig. S11. Simulated evolution under asymmetric competition kernel defined by Eq. S6.2 with $m_\alpha = \sigma_\alpha/2$, plotted in the same manner as in Fig. S2. The other settings and parameters were the same with those for the simulated evolution shown in Fig. S2.

### 7 Robustness: Rare mutation for fundamental trait

#### 7.A Prediction by adaptation front equation

When mutation occurs separately for traits $x$ and $y$, the mutational covariance matrix is calculated from Eq. S2.2 as

$$\begin{aligned}
\mathbf{V}_\mu &= \begin{pmatrix} V_{\mu xx} & V_{\mu xy} \\ V_{\mu xy} & V_{\mu yy} \end{pmatrix} \\
&= \int\int \xi(x', y'; x, y) \begin{pmatrix} [x'-x]^2 & [x'-x][y'-y] \\ [x'-x][y'-y] & [y'-y]^2 \end{pmatrix} dxdy \\
&= \begin{pmatrix} \mu_x \sigma_{\mu x}^2/\mu & 0 \\ 0 & \mu_y \sigma_{\mu y}^2/\mu \end{pmatrix}, \qquad \text{[S7.1]}
\end{aligned}$$

where $\mu_x$ and $\mu_y$ describe mutation rates for $x$ and $y$, respectively, and $\mu = \mu_x + \mu_y$ is the total mutation rate. $\sigma_{\mu x}$ describes the average mutation size for $x$ (i.e., standard deviation among mutations in $x$), while $\sigma_{\mu y}$ describes the average mutation size for $y$ (i.e., standard deviation among mutations in $y$). Note that $\sigma_{\mu x}$ and $\sigma_{\mu y}$ differ from the average mutation sizes $\sigma_{\mu xx}$ and $\sigma_{\mu yy}$ for the case that Eq. S2.2 is treated as a non-separated mutation distribution (i.e., $\sigma_{\mu xx} = \sqrt{V_{\mu \mathrm{xx}}} = \sqrt{\mu_x \sigma_{\mu x}^2/\mu}$ and $\sigma_{\mu yy} = \sqrt{V_{\mu yy}} = \sqrt{\mu_y \sigma_{\mu y}^2/\mu}$).

When $\mu_y/\mu_x$ is very small, as explained in the next subsection, species-level selection has a large contribution to the progress of adaptation front in the simulation, measured by the apparent amplification $\eta$ of the mutation rate for trait $y$. This effect can be incorporated in the adaptation front equation by replacing $\mu_y$ with $\eta\mu_y$ in the equation. Then, for the case considered here, the general form of the rescaled adaptation front equation for two-dimensional trait spaces, Eq. S8.19, is simplified into

$$\begin{aligned} \frac{\partial h(x,\tau)}{\partial \tau} &= v_{hy}(x)\left[V_{\mu,x/y}\left|\frac{\partial h(x,\tau)}{\partial x}\right|^2 + 1\right], \\ v_{hy}(x) &= \frac{K(x)}{K(x^*)}, \\ V_{\mu,x/y} &= \frac{V_{\mu xx}}{V_{\mu yy}} = \frac{1}{\eta}\left[\frac{\mu_x\sigma_{\mu x}^2}{\mu_y\sigma_{\mu y}^2}\right] \end{aligned} \quad \text{[S7.2]}$$

(by substitution of $\mu(x,t) = \mu = \mu_x + \eta\mu_y$, $V_{\mu xx}(x,t) = \mu_x\sigma_{\mu x}^2/\mu$, $V_{\mu yy}(x,t) = \eta\mu_y\sigma_{\mu y}^2/\mu$, $K_E(x,t) = [\beta(x)h(x,t)+1]K(x)$, $g_y(x,t) = \beta$, $\bar{\rho}(x,t) = \bar{\rho}$), provided that the front top is located at $x = x^*$ that maximizes $v_{hy}(x)$ for sufficiently large $t$. Clearly, we see that $x^* = 0$, which gives $K(x^*) = 1$. By using Eqs. S8.20-S8.22, we obtain

$$\begin{aligned} H(x) &= -\frac{x}{|x|}\int_0^x \frac{1}{\sqrt{V_{\mu,x/y}(\tilde{x})}}\sqrt{\frac{1}{v_{hy}(\tilde{x})} - 1}d\tilde{x} = -\sqrt{\eta}\sqrt{\frac{\mu_y\sigma_{\mu y}^2}{\mu_x\sigma_{\mu x}^2}}\frac{x}{|x|}\int_0^x \sqrt{\frac{1}{K(\tilde{x})} - 1}, \\ v_x(x) &= -v_{hy}(x)V_{\mu,x/y}\frac{dH(x)}{dx} = \frac{x}{|x|}\frac{1}{\sqrt{\eta}}\sqrt{\frac{\mu_x\sigma_{\mu x}^2}{\mu_y\sigma_{\mu y}^2}}\sqrt{K(x)[1-K(x)]}, \\ v_y(x) &= v_{hy}(x) = K(x), \\ D(x) &= \frac{\partial v_x(x)}{\partial x} = \frac{1}{\sqrt{\eta}}\sqrt{\frac{\mu_x\sigma_{\mu x}^2}{\mu_y\sigma_{\mu y}^2}}\sqrt{\frac{K(x)}{1-K(x)}}\left|\frac{d\log K(x)}{dx}\right|\left[K(x) - \frac{1}{2}\right], \\ p_d(x) &= \sigma_a\sqrt{\left[\frac{dH(x)}{dx}\right]^2 + 1} = \sigma_\alpha\sqrt{\eta\frac{\mu_y\sigma_{\mu y}^2}{\mu_x\sigma_{\mu x}^2}\left[\frac{1}{K(x)} - 1\right] + 1}, \\ \tau_d(x) &= |H(x)| = \sqrt{\eta}\sqrt{\frac{\mu_y\sigma_{\mu y}^2}{\mu_x\sigma_{\mu x}^2}}\int_0^x \sqrt{\frac{1}{K(\tilde{x})} - 1}. \end{aligned} \quad \text{[S7.3]}$$

Fig. S12 shows these predictions (Eq. S7.3) and the simulated evolution (same with Fig. 4 in the main text) for $\mu_y/\mu_x = 2.5 \times 10^{-5}$, $\sigma_{\mu y}/\sigma_{\mu x} = 2.0 \times 10^2$, satisfying $\mu_x\sigma_{\mu x}^2/[\mu_y\sigma_{\mu y}^2] = 1$, and $\eta = 9.92$ estimated by using Eq. S7.13.

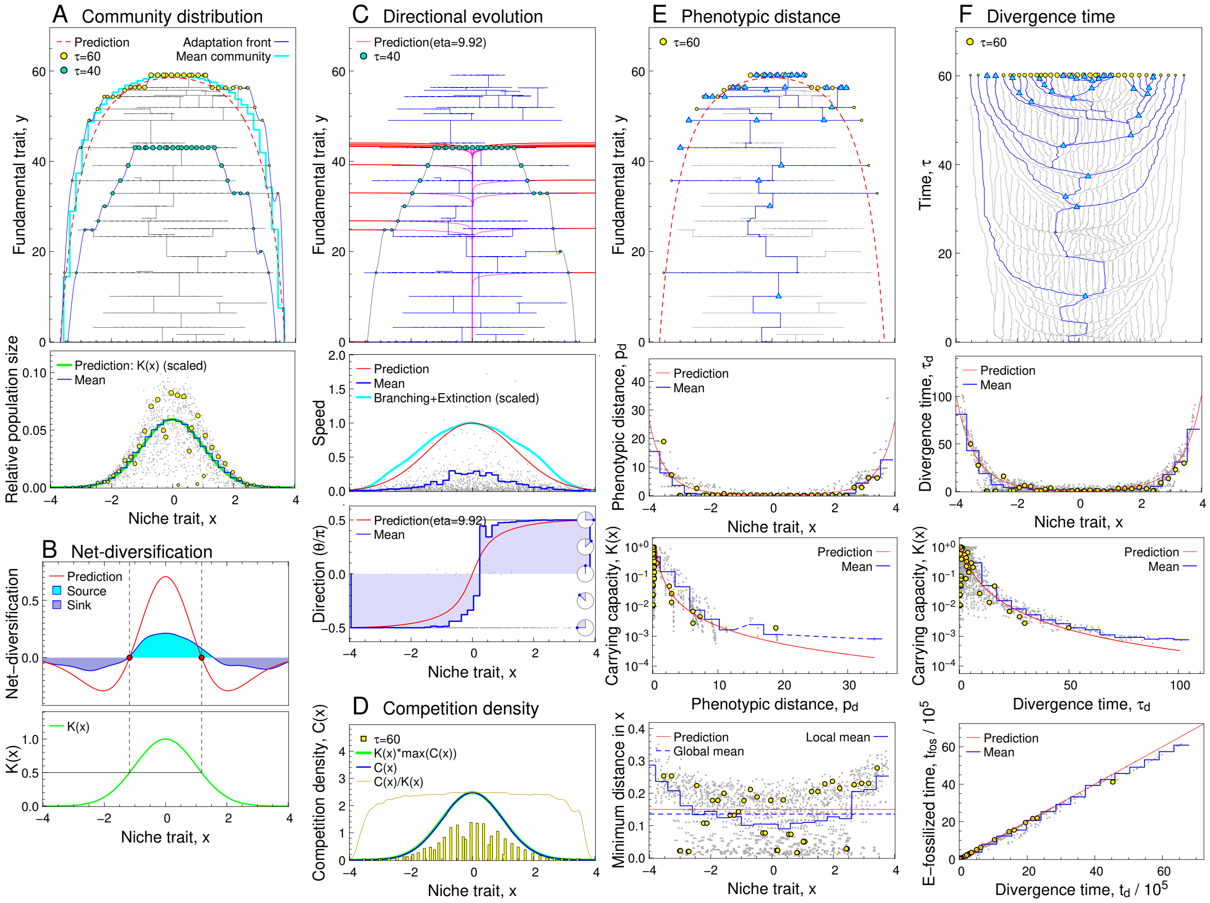

Fig. S12. Simulated evolution under rare mutation for trait $y$, plotted in the same manner as in Fig. S2. The mutation distribution was given by Eq. S2.2 with $\mu_x = 1$, $\mu_y = 2.5 \times 10^{-5}$, $\sigma_{\mu x} = 0.01$, $\sigma_{\mu y} = 2.0$, satisfying $\mu_x \sigma_{\mu x}^2 / [\mu_y \sigma_{\mu y}^2] = 1$. The other settings and parameters were the same with those for the simulated evolution shown in Fig. S2. Since the growth of $\bar{y}$ was not monotonic along the non-rescaled time $t$ in this case, we calculated the rescaled time, $\tau$, as $\bar{y}$ with diffusion smoothing over $t$, denoted by $\tau = \bar{y}_{\rm sm}(t; 1000)$, where $\bar{y}_{\rm sm}(t; k) = \bar{y}_{\rm sm}(t; k-1) + 0.1 \cdot [y(t+\Delta t; k-1) - 2y(t; k-1) + y(t-\Delta t; k-1)]$ for $k = 1, \ldots, 1000$ with $\bar{y}_{\rm sm}(t_{\rm min}; k)$ and $\bar{y}_{\rm sm}(t_{\rm max}; k)$ being fixed, and $\bar{y}_{\rm sm}(t, 0)$ was given by $\bar{y}(t)$ for $t = 0, \Delta t, 2\Delta t, \ldots$, obtained by equally-spaced 500 sampling from $t_{\rm min} = 0$ to $t_{\rm max} = 9.76 \times 10^6$ (corresponding to $\tau = 150$), i.e., $\Delta t = [t_{\rm max} - t_{\rm min}]/500$.

#### 7.B Effect of species-level selection

We derive an estimation formula for $\eta$ (i.e., the apparent amplification of mutation rate for trait $y$), which measures the effect of species-level selection. The adaptation horizon without the effect of species-level selection is given by Eq. S7.3 with $\eta = 1$,

$$H(x) = -\sqrt{\frac{\mu_y \sigma_{\mu y}^2}{\mu_x \sigma_{\mu x}^2}} \frac{x}{|x|} \int_0^x \sqrt{\frac{1}{K(\tilde{x})} - 1} d\tilde{x}. \quad [S7.4]$$

We start with assuming that $\mu_y/\mu_x$ is extremely small while $\sigma_{\mu y}/\sigma_{\mu x}$ is extremely large, so that $\mu_y \sigma_{\mu y}^2$ has the same order of magnitude with $\mu_x \sigma_{\mu x}^2$. In this case, the $k$th invasion by a mutant with improved $y = y_k$, referred to as the $k$th innovated mutant, produces the $k$th clade that excludes all species having less innovated $y$, followed by an extremely long waiting time for emergence of the $(k+1)$th innovated mutant. Then, the emergence rate of the next innovated mutant is expected to be proportional to the total number of individuals in the clade monopolizing the system, denoted by $n_{\mathrm{clade},k}$. As in the main text, we assume that the expected between-species niche distance is given by $\sigma_\alpha$, and that the expected population size for a species occupying niche $x$ is given by $n_{\max,k}K(x)$, where $n_{\max,k}$ describes the expected population size for a species occupying the optimum niche $x = 0$ in the $k$-th clade monopolizing the system. Then, we can express $n_{\mathrm{clade},k}$ as

$$n_{\mathrm{clade},k} = \sum_{i}^{N} \hat{n}_i \simeq \sum_{j=-\infty}^{\infty} n_{\max,k} K(j\sigma_\alpha) \simeq \frac{n_{\max,k} \int_{-\infty}^{\infty} K(x)\mathrm{d}x}{\sigma_\alpha}. \qquad \text{[S7.5]}$$

Hence, the expected mutant production from the clade is greater than that from the species occupying $x = 0$, by factor

$$\eta = \frac{n_{\mathrm{clade},k}}{n_{\max,k}} \simeq \frac{\int_{-\infty}^{\infty} K(x)\mathrm{d}x}{\sigma_\alpha} =: \eta_{\max}. \qquad \text{[S7.6]}$$

Therefore, in the non-rescaled time $t$ (where the adaptation front equation is given by Eq. S7.14 with $\eta = 1$), we expect that the progress speed of adaptation front in the simulated evolution is faster than the prediction made from the adaptation front equation (i.e., the expected evolutionary speed of the species occupying $x = 0$), by factor $\eta_{\max}$. When $K(x)$ is a Gaussian distribution with standard deviation $\sigma_K$, we see from Eq. S7.6 that $\eta_{\max} = \sqrt{2\pi}\sigma_K/\sigma_\alpha$ (e.g., $\eta_{\max} = 16.7$ for the simulated evolution in Fig. 4 in the main text, i.e., $\sigma_\alpha = 0.15$ and $\sigma_K = 1.0$).

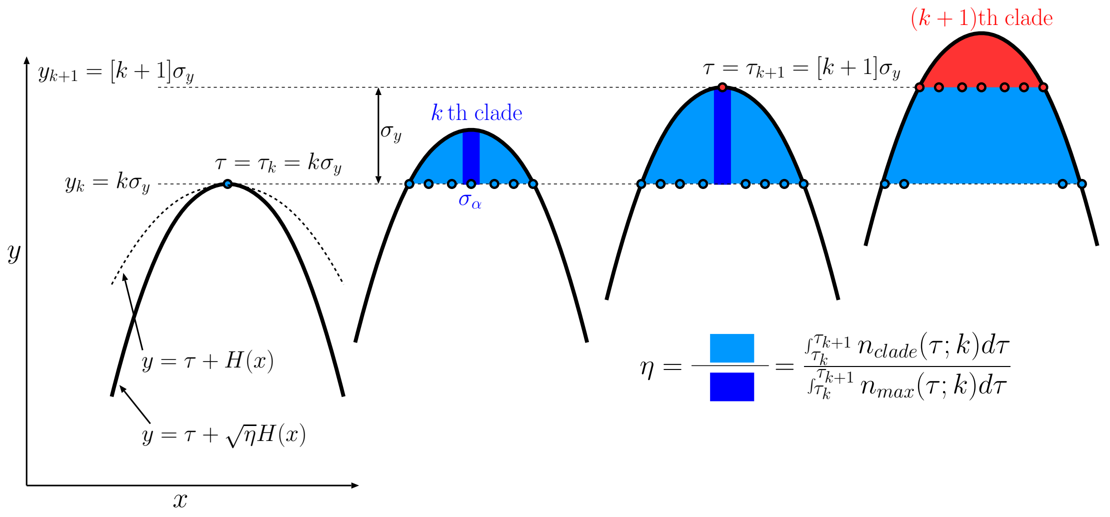

Fig. S13. Illustration of community progress in trait $y$.

Next, we consider the case that $\mu_y/\mu_x$ is still significantly small but not extremely small, so that the $(k+1)$th innovated mutant may emerge before the $k$th clade monopolizes the system. In order to derive the estimation formula for $\eta$ in such a case, we make the following four assumptions. First, the $(k+1)$th innovated mutant always emerges from the $k$th clade. Second, the dynamics of adaptation front is given by

$$y = h(x,\tau) = \tau + \sqrt{\eta}H(x), \qquad \text{[S7.7]}$$

and the $k$th innovated mutant has phenotype $(x, y) = (0, k\sigma_{\mu y})$ and emerges at time $\tau_k = k\sigma_{\mu y}$. Third, the niche region covered by the $k$th clade at time $\tau$ (before the emergence of the $(k+1)$th clade) is given by

$$\mathcal{X}(\tau - \tau_k) = \{x | \tau - \tau_k + \sqrt{\eta}H(x) \geqslant 0\} \qquad \text{[S7.8]}$$

(Fig. S13), so that the expected number of individuals in the $k$th clade at time $\tau$ is given by

$$n_{\text{clade}}(\tau, k) \;\simeq\; \frac{\int_{x \in \mathcal{X}(\tau-\tau_k)} n_{\max,k} K(x) dx}{\sigma_\alpha}. \qquad \text{[S7.9]}$$

Fourth, during the time period from $\tau_k$ to $\tau_{k+1}$, the relationship between the rescaled time $\tau$ and the non-rescaled time $t$ are approximately linear, so that the expected number of mutants that emerge from the $k$th clade within the time period from $\tau_k$ to $\tau_{k+1}$ can be expressed as

$$\begin{aligned} c(\tau)\int_{\tau_k}^{\tau_{k+1}} n_{\text{clade}}(\tau, k) d\tau &\simeq \frac{c(\tau)}{\sigma_\alpha}\int_{\tau_k}^{\tau_{k+1}}\int_{x\in\mathcal{X}(\tau-\tau_k)} n_{\max,k}K(x)dxd\tau \\ &= \frac{c(\tau)}{\sigma_\alpha}\int_{x\in\mathcal{X}(\tau-\tau_k)} n_{\max,k}K(x)\left[\tau_{k+1}-\tau_k+\sqrt{\eta}H(x)\right]dx \\ &= \frac{c(\tau)n_{\max,k}}{\sigma_\alpha}\int K(x)\left[\sigma_{\mu y}+\sqrt{\eta}H(x)\right]_+ dx, \end{aligned} \qquad \text{[S7.10]}$$

with $c(\tau) = \mu_y[dt/d\tau]$, where transformation from the first row to the second is the interchange of integration order between $x$ and $\tau$ (see Fig. S13), and $[\bullet]_+$ converts all negative values for $\bullet$ into zero. On the other hand, the expected number of mutants that emerge from the species occupying $x = 0$ in the $k$th clade within the time period from $\tau_k$ to $\tau_{k+1}$ can be expressed as

$$c(\tau)\int_{\tau_k}^{\tau_{k+1}} n_{\max,k} d\tau \;=\; c(\tau)\sigma_{\mu y} n_{\max,k}. \qquad \text{[S7.11]}$$

Hence, $\eta$ is expressed as

$$\eta \;\simeq\; \frac{c(\tau)\int_{\tau_k}^{\tau_{k+1}} n_{\text{clade},k}(\tau,k)d\tau}{c(\tau)\int_{\tau_k}^{\tau_{k+1}} n_{\max,k}d\tau} = \frac{1}{\sigma_\alpha}\int K(x)\left[1+\frac{\sqrt{\eta}}{\sigma_{\mu y}}H(x)\right]_+ \text{dx}. \qquad \text{[S7.12]}$$

Finally, substitution of Eq. S7.4 into Eq. S7.12 yields that

$$\eta_{\text{est}} \;=\; \frac{1}{\sigma_\alpha}\int_{-\infty}^{\infty} K(x)\left[1-\sqrt{\eta_{\text{est}}}\sqrt{\frac{\mu_y}{\mu_x\sigma_x^2}}\frac{x}{|x|}\int_0^x\sqrt{\frac{1}{K(\tilde{x})}-1d\tilde{x}}\right]_+ \text{dx} \qquad \text{[S7.13]}$$

approximately holds. Note that this equation has $\eta_{\text{est}}$ in both sides. In the present study, $\eta_{\text{est}}$ is obtained by solving this equation numerically. For example, $\eta_{\text{est}} = 9.92$ is obtained from the parameter set used for Fig. 4 and Fig. S12 ($\mu_y/\mu_x = 2\times10^{-5}$, $\sigma_x = 0.01$, $\sigma_\alpha = 0.15$, and $\sigma_K = 1$ for the Gaussian $K(x)$). Note that an infinitesimally small $\mu_y$ such that the $\eta_{\text{est}}$ in the right-hand side is negligible gives Eq. S7.6, i.e., $\eta_{\text{est}} = \int_{-\infty}^{\infty} K(x)dx/\sigma_\alpha = \eta_{\max}$.

From the simulated evolution, we can calculate $\eta$ by using the non-rescaled adaptation front equation:

$$\frac{\partial h(x,t)}{\partial t} \;=\; \eta\frac{\mu_y\sigma_{\mu y}^2\beta}{2}\frac{[\beta h(x,t)+1]K(x)}{\bar{\rho}}\left[\left|\frac{\partial h(x,t)}{\partial x}\right|^2+1\right] \qquad \text{[S7.14]}$$

(Eq. S8.18 with $\mu(x,t) = \mu$, $V_{\mu xx}(x,t) = \mu_x\sigma_{\mu x}^2/\mu$, $V_{\mu yy}(x,t) = \eta\mu_y\sigma_{\mu y}^2/\mu$, $K_E(x,t) = [\beta(x)h(x,t) + 1]K(x)$, $g_y(x,t) = \beta$, $\bar{\rho}(x,t) = \bar{\rho}$). By solving this equation for $\eta$, we get

$$\eta_{\text{sim}}(t) \;=\; \frac{2\sqrt{2\pi}\bar{\rho}}{\mu_y\sigma_{\mu y}^2\beta[\beta h(0,t)+1]}\frac{\partial h(0,t)}{\partial t}, \qquad \text{[S7.15]}$$

where $h(0,t)$ and $\frac{\partial h(0,t)}{\partial t}$ are calculated from the simulated evolution.

As shown in Fig. S14, the $\eta_{\text{est}}$ obtained by Eq. S7.13 seems to work for predicting the average value for the $\eta_{\text{sim}}(t)$ calculated from the simulated evolution by using Eq. S7.15, not only when $\mu_y/\mu_x$ is significantly small ($\mu_y/\mu_x = 2 \times 10^{-5}$) but also when it is equal to 1.

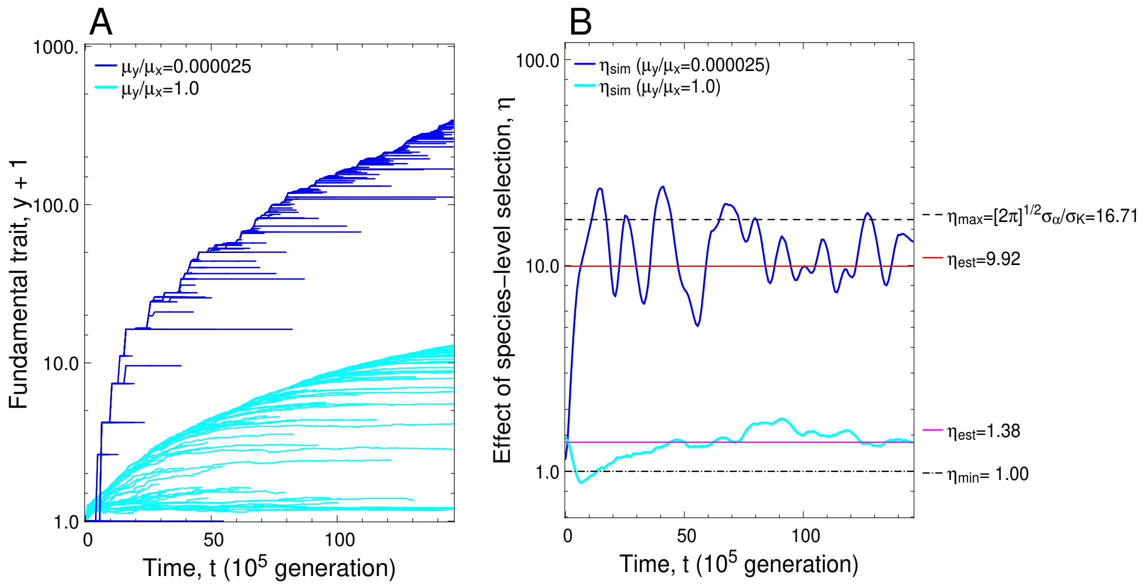

Fig. S14. Effect of species-level selection in community evolution in trait $y$. In panel (A), blue curves indicate species' trajectories on the $t$-$y$ plane for the simulated evolution shown in Fig. S12. Cyan curves indicate species' trajectories for the simulated evolution under the same setting, except that the mutation was isotropic, where its rate $\mu$ and average size $\sigma_\mu$ were chosen so that $\mu\sigma_\mu^2 = \mu_x\sigma_{\mu x}^2 = \mu_y\sigma_{\mu y}^2$. In panel (B), $\eta_{\text{sim}}(t)$ for the blue ($\mu_y/\mu_x = 2 \times 10^{-5}$) and cyan ( $\mu_y/\mu_x = 1$) trajectories in panel (A) are plotted with the corresponding colors, which are compared with the red ($\eta_{\text{est}} = 9.92$) and purple ($\eta_{\text{est}} = 1.38$) lines obtained from Eq. S7.13 for $\mu_y/\mu_x = 2 \times 10^{-5}$ and $\mu_y/\mu_x = 1$, respectively (with $\sigma_x = 0.01$, $\sigma_\alpha = 0.15$, and $\sigma_K = 1$). For the calculation of $\eta_{\text{sim}}(t)$, community states were sampled (with equally-spaced time sampling with $\Delta t = 80000$ from $\tau = 0$ to $\tau = 150$). $h(0,t)$ and $\frac{\partial h(0,t)}{\partial t}$ in Eq. S7.15 were calculated as $h(0,t) = \bar{y}$ (with the same smoothing for $\tau$ in Fig. S12) and $\frac{\partial h(0,t)}{\partial t} = \Delta\bar{y}/\Delta t$, respectively, at each sampling time point. $\bar{\rho}$ was chosen at $\bar{\rho} = \sqrt{2\pi}$, as derived in Eq. S10.28 for the Gaussian competition kernel.

#### 7.C Prediction curves in Fig. 4C and 4D

Regarding Fig. 4C, we see from Fig. S13 that the niche region occupied by the $k$th clade, after emergence of the $(k+1)$th clade, can be expressed as

$$\mathcal{X}(\tau - \tau_k) \quad = \quad \{x | \tau_k \leqslant \tau + \sqrt{\eta} H(x) \leqslant \tau_k + \sigma_{\mu y}\}\,. \tag{S7.16}$$

Under the assumption of the constant between-species niche distance, $\sigma_\alpha$, we can express the number of species in the $k$th clade as

$$N_{\text{sp}}(\tau - \tau_k) \quad = \quad \frac{\int_{x \in \mathcal{X}(\tau - \tau_k)} 1 dx}{\sigma_\alpha} \tag{S7.17}$$

In addition, the "within-clade minimum $|x|$" is given by

$$|x|_{\min}(\tau - \tau_k) \quad = \quad \min(\{|x| | x \in \mathcal{X}(\tau - \tau_k)\}). \tag{S7.18}$$

The prediction curve in Fig. 4C is defined by Eqs. S7.17 and S7.18 with $\tau - \tau_k$ as a parameter.

Regarding the prediction formula for Fig. 4D, we see from Fig. S13 that the age of the $k$th clade, $\tau - \tau_k$, can be expressed as

$$\begin{aligned} \tau - \tau_k &= \sqrt{\eta}|H(|x|_{\max}(\tau - \tau_k))|, \\ |x|_{\max}(\tau - \tau_k) &= \max(\{|x||x \in \mathcal{X}(\tau - \tau_k)\}). \end{aligned} \qquad \text{[S7.19]}$$

Since $|x|_{\max}(\tau - \tau_k) \simeq |x|_{\min}(\tau - \tau_k)$ holds for sufficiently large $\tau - \tau_k$, the prediction curve in Fig. 4D is defined by $\tau - \tau_k = \sqrt{\eta}|H(|x|_{\min}(\tau - \tau_k))|$ and Eq. S7.18 with $\tau - \tau_k$ as a parameter.

##### 7.D Advancement speed and species diversity

From Eq. S7.5, the relative population size $p_i = \hat{n}_i / \sum_{j=1}^{N} \hat{n}_j$ for the $i$th species is approximately expressed as

$$p_i = n_{\max,k} K(x_i) \left[ \frac{n_{\max} \int_{-\infty}^{\infty} K(x) \mathrm{dx}}{\sigma_\alpha} \right]^{-1} = \frac{\sigma_\alpha K(x_i)}{\int_{-\infty}^{\infty} K(x) \mathrm{dx}}. \qquad \text{[S7.20]}$$

On this basis, we can calculate the effective number of species in the system, denoted by $N_{\mathrm{sh}}$, as the exponential of the Shannon species diversity:

$$\begin{aligned} N_{\mathrm{sh}} &= \exp\left( -\sum_{i=1}^{N} p_i \log p_i \right) \simeq \frac{\int_{-\infty}^{\infty} K(x) \mathrm{dx}}{\sigma_\alpha} Q = \eta_{\max} Q, \\ Q &= \exp\left( \frac{\int_{-\infty}^{\infty} K(x)[-\log K(x)] \mathrm{dx}}{\int_{-\infty}^{\infty} K(x) \mathrm{dx}} \right). \end{aligned} \qquad \text{[S7.21]}$$

If $K(x)$ is a Gaussian distribution, $K(x) = \exp\left(\frac{-x^2}{2\sigma_K^2}\right)$ (Eq. 2 in the main text), then $Q = \exp\left(\frac{1}{2}\right)$ holds for any $\sigma_K$, resulting in $\eta_{\max} = N_{\mathrm{sh}} / \exp(\frac{1}{2})$. In this case, larger effective species number gives faster potential speed for the community advancement.

#### 8 Extension for higher-dimensional trait space

##### 8.A Generalized adaptation front equation

We consider an arbitrary $M$-dimensional trait space $\mathbf{s} = (x_{.1}, \ldots, x_{.M})^T$, where ".$l$" for $l = 1, \ldots, M$ indicates the $l$-th component of $\mathbf{s}$. This subscript rule is introduced for distinguishing the subscripts for trait axes from the identification numbers for species. This rule allows us to denote the phenotype of the $i$th species by $\mathbf{s}_i = (x_{i.1}, \ldots, x_{i.M})^T$ without confusion. We assume $N$ coexisting resident phenotypes $\mathbf{s}_1, \ldots, \mathbf{s}_N$, each of which is treated as a species. Phenotypes of the coexisting species are combined into a matrix $\mathbf{S} = (\mathbf{s}_1, \ldots, \mathbf{s}_N)$. Their population dynamics are written as

$$\frac{dn_i}{dt} = n_i F(\mathbf{s}_i; \mathbf{S}, \mathbf{n}) \qquad \text{[S8.1]}$$

for $i = 1, \ldots, N$, where $\mathbf{n} = (n_1, \ldots, n_N)^T$ describes population sizes of the coexisting species, and $F(\mathbf{s}_i; \mathbf{S}, \mathbf{n})$ describes an arbitrary fitness function, i.e., per capita growth rate for $\mathbf{s}_i$ under residents $\mathbf{S}$ with their population sizes $\mathbf{n}$. We assume that the invasion fitness of a mutant phenotype $\mathbf{s}' = (x'_{.1}, \ldots, x'_{.M})^T$ is expressed as $f(\mathbf{s}'; \mathbf{S}) = F(\mathbf{s}'; \mathbf{S}, \hat{\mathbf{n}})$ with equilibrium population sizes $\hat{\mathbf{n}} = (\hat{n}_1, \ldots, \hat{n}_N)^T$ for the coexisting species, which is a function of $\mathbf{S}$.

We describe the directional evolution of each species with the canonical equation (4):

$$\frac{d\mathbf{s}_i}{dt} = \frac{\mu(\mathbf{s}_i)}{2}\hat{n}_i \mathbf{V}_\mu(\mathbf{s}_i)\mathbf{g}(\mathbf{s}_i;\mathbf{S}) \qquad \text{[S8.2]}$$

for $i = 1, \ldots, N$, where an $M \times M$ matrix $\mathbf{V}_\mu(\mathbf{s}_i)$ describes the mutational covariance matrix. $\mathbf{V}_\mu(\mathbf{s})$ and $\mu(\mathbf{s})$ may vary depending on parental phenotypes $\mathbf{s}$ of mutants. $\mathbf{g}(\mathbf{s};\mathbf{S})$ describes the fitness gradient for $\mathbf{s}$, given by

$$\mathbf{g}(\mathbf{s};\mathbf{S}) = \left[\frac{\partial f(\mathbf{s}';\mathbf{S})}{\partial \mathbf{s}'}\right]_{\mathbf{s}'=\mathbf{s}} = \begin{pmatrix} \frac{\partial f(\mathbf{s}';\mathbf{S})}{\partial x'_{.1}} \\ \vdots \\ \frac{\partial f(\mathbf{s}';\mathbf{S})}{\partial x'_{.L}} \end{pmatrix}_{\mathbf{s}'=\mathbf{s}} . \qquad \text{[S8.3]}$$

Here, for the notational clarity, the dependency of $\mathbf{g}(\mathbf{s};\mathbf{S})$ on $\mathbf{S}$ is explicitly shown as one of the arguments, differently from the case of two-dimensional trait space. We assume that the trait space can be decomposed into an $L$-dimensional subspace for niche traits $\mathbf{x} = (x_{.1}, \ldots, x_{.L})^T$ (undergoing negative frequency-dependent selection through competition with narrow interaction ranges) and into an $(M-L)$-dimensional subspace for fundamental traits $\mathbf{y} = (y_{.1}, \ldots, y_{.M-L})^T = (x_{.L+1}, \ldots, x_{.M})^T$ (undergoing weak and monotonic directional selection). In this case, the zero-isocline of the invasion fitness can be expressed as

$$\mathbf{y} = \begin{pmatrix} h_{.1}(\mathbf{x},t) \\ \vdots \\ h_{.M-L}(\mathbf{x},t) \end{pmatrix} = \mathbf{h}(\mathbf{x},t). \qquad \text{[S8.4]}$$

Then, as derived in Eq. S11.6, the fitness gradient for $\mathbf{s} = \begin{pmatrix} \mathbf{x} \\ \mathbf{h}(\mathbf{x},t) \end{pmatrix}$ on the adaptation front is expressed as

$$\mathbf{g}(\mathbf{x},t) = \begin{pmatrix} \mathbf{g}_\mathbf{x}(\mathbf{x},t) \\ \mathbf{g}_\mathbf{y}(\mathbf{x},t) \end{pmatrix} = \begin{pmatrix} \frac{\partial f(\mathbf{s}';\mathbf{S})}{\partial \mathbf{x}'} \\ \frac{\partial f(\mathbf{s}';\mathbf{S})}{\partial \mathbf{y}'} \end{pmatrix}_{\mathbf{s}'=(\mathbf{x},\mathbf{h}(\mathbf{x},t))^T} = \begin{pmatrix} -\frac{\partial \mathbf{h}(\mathbf{x},t)^T}{\partial \mathbf{x}} \\ \mathbf{I}_\mathbf{y} \end{pmatrix} \mathbf{g}_\mathbf{y}(\mathbf{x},t), \qquad \text{[S8.5]}$$

where $\mathbf{I}_\mathbf{y}$ is the $(M-L)$-by-$(M-L)$ identity matrix.

On this basis, in a manner analogous to the case of the two-dimensional trait space $\mathbf{s} = (x,y)^T$, we can derive the expected dynamics of the adaptation front (see SI Appendix, section 11A and 11B for the derivation) as

$$\frac{\partial \mathbf{h}(\mathbf{x},t)}{\partial t} = \frac{\mu(\mathbf{x},t)\sigma_h(\mathbf{x},t)^2}{2}\bar{n}(\mathbf{x},t)\left[\frac{\partial \mathbf{h}(\mathbf{x},t)}{\partial \mathbf{x}^T}\frac{\partial \mathbf{h}(\mathbf{x},t)^T}{\partial \mathbf{x}} + \mathbf{I}_\mathbf{y}\right]\mathbf{g}_\mathbf{y}(\mathbf{x},t). \qquad \text{[S8.6]}$$

Here, $\bar{n}(\mathbf{x},t)$ describes the expected population size for a species of phenotype $\mathbf{s} = \begin{pmatrix} \mathbf{x} \\ \mathbf{h}(\mathbf{x},t) \end{pmatrix}$, given by

$$\begin{aligned} \bar{n}(\mathbf{x},t) &= \frac{K_E(\mathbf{x},t)}{\bar{\rho}(\mathbf{x},t)}, \\ K_E(\mathbf{x},t) &= \frac{\sum_{j=1}^N b(\mathbf{s},\mathbf{s}_j;\mathbf{S})\hat{n}_j}{b(\mathbf{s},\mathbf{s};\mathbf{S})}, \\ b(\mathbf{s}_i,\mathbf{s}_j;\mathbf{S}) &= \left[\frac{\partial F(\mathbf{s}_i;\mathbf{S},\mathbf{n})}{\partial n_j}\right]_{\mathbf{n}=\hat{\mathbf{n}}} . \end{aligned} \qquad \text{[S8.7]}$$

$\mu(\mathbf{x},t)$ and $\sigma_h(\mathbf{x},t)$ respectively describe the mutation rate and the average mutation size for phenotype $\mathbf{s} = \begin{pmatrix} \mathbf{x} \\ \mathbf{h}(\mathbf{x},t) \end{pmatrix}$ along the orthogonal direction to the adaptation front, given by

$$\begin{aligned} \mu(\mathbf{x},t) &= \mu(\mathbf{s}), \\ \sigma_h(\mathbf{x},t)^2 &= \frac{\mathbf{g}(\mathbf{x},t)^T \mathbf{V}_\mu(\mathbf{x},t)\mathbf{g}(\mathbf{x},t)}{|\mathbf{g}(\mathbf{x},t)|^2}, \\ \mathbf{V}_\mu(\mathbf{x},t) &= \mathbf{V}_\mu(\mathbf{s}). \end{aligned} \quad \text{[S8.8]}$$

$\bar{\rho}(\mathbf{x},t)$ describes the locally-averaged (i.e., ”expected” in the main text) competing species density around $\mathbf{x}$ at time $t$, defined by Eq. S11.25. The $\frac{\partial \mathbf{h}(\mathbf{x},t)^T}{\partial \mathbf{x}}$ and $\frac{\partial \mathbf{h}(\mathbf{x},t)}{\partial \mathbf{x}^T}$ are defined by

$$\begin{aligned} \frac{\partial \mathbf{h}(\mathbf{x},t)^T}{\partial \mathbf{x}} &= \begin{pmatrix} \frac{\partial \mathbf{h}(\mathbf{x},t)^T}{\partial x_{.1}} \\ \vdots \\ \frac{\partial \mathbf{h}(\mathbf{x},t)^T}{\partial x_{.L}} \end{pmatrix} = \begin{pmatrix} \frac{\partial h_{.1}(\mathbf{x},t)}{\partial x_{.1}} & \cdots & \frac{\partial h_{.M-L}(\mathbf{x},t)}{\partial x_{.1}} \\ \vdots & \ddots & \vdots \\ \frac{\partial h_{.1}(\mathbf{x},t)}{\partial x_{.L}} & \cdots & \frac{\partial h_{.M-L}(\mathbf{x},t)}{\partial x_{.L}} \end{pmatrix}, \\ \frac{\partial \mathbf{h}(\mathbf{x},t)}{\partial \mathbf{x}^T} &= \left[ \frac{\partial \mathbf{h}(\mathbf{x},t)^T}{\partial \mathbf{x}} \right]^T. \end{aligned} \quad \text{[S8.9]}$$

#### 8.B Directional evolution

In the right-hand side of Eq. S8.2, replacements of $\mathbf{s}_i$ with $\mathbf{s} = (\mathbf{x}, \mathbf{h}(\mathbf{x},t))^T$ and of $\hat{n}_i$ with $\bar{n}(\mathbf{x},t) = \frac{K_E(\mathbf{x},t)}{\bar{\rho}(\mathbf{x},t)}$ give the expected directional evolution for a species of phenotype $\mathbf{s} = (\mathbf{x}, \mathbf{h}(\mathbf{x},t))^T$:

$$\begin{aligned} \begin{pmatrix} \mathbf{u}_\mathbf{x}(\mathbf{x},t) \\ \mathbf{u}_\mathbf{y}(\mathbf{x},t) \end{pmatrix} &= \mathbf{u}(\mathbf{x},t) = \frac{\mu(\mathbf{x},t)}{2}\bar{n}(\mathbf{x},t)\mathbf{V}_\mu(\mathbf{x},t)\mathbf{g}(\mathbf{x},t) \\ &= \frac{\mu(\mathbf{x},t)}{2}\bar{n}(\mathbf{x},t) \begin{pmatrix} \mathbf{V}_{\mu\mathbf{xx}}(\mathbf{x},t) & \mathbf{V}_{\mu\mathbf{xy}}(\mathbf{x},t) \\ \mathbf{V}_{\mu\mathbf{yx}}(\mathbf{x},t) & \mathbf{V}_{\mu\mathbf{yy}}(\mathbf{x},t) \end{pmatrix} \begin{pmatrix} -\frac{\partial \mathbf{h}(\mathbf{x},t)^T}{\partial \mathbf{x}} \mathbf{g}_\mathbf{y}(\mathbf{x},t) \\ \mathbf{g}_\mathbf{y}(\mathbf{x},t) \end{pmatrix} \\ &= \begin{pmatrix} \frac{\mu(\mathbf{x},t)}{2} \frac{K_E(\mathbf{x},t)}{\bar{\rho}(\mathbf{x},t)} \left[ -\mathbf{V}_{\mu\mathbf{xx}}(\mathbf{x},t) \frac{\partial \mathbf{h}(\mathbf{x},t)^T}{\partial \mathbf{x}} + \mathbf{V}_{\mu\mathbf{xy}}(\mathbf{x},t) \right] \mathbf{g}_\mathbf{y}(\mathbf{x},t) \\ \frac{\mu(\mathbf{x},t)}{2} \frac{K_E(\mathbf{x},t)}{\bar{\rho}(\mathbf{x},t)} \left[ -\mathbf{V}_{\mu\mathbf{yx}}(\mathbf{x},t) \frac{\partial \mathbf{h}(\mathbf{x},t)^T}{\partial \mathbf{x}} + \mathbf{V}_{\mu\mathbf{yy}}(\mathbf{x},t) \right] \mathbf{g}_\mathbf{y}(\mathbf{x},t) \end{pmatrix}. \end{aligned} \quad \text{[S8.10]}$$

#### 8.C Net-diversification

Provided that both $\frac{\partial \ln \bar{\rho}(\mathbf{x},t)}{\partial t} \simeq 0$ and $\frac{\partial \ln \bar{\rho}(\mathbf{x},t)}{\partial \mathbf{x}} \simeq 0$ are kept through the evolutionary dynamics, we derive the net-diversification rate for $\mathbf{x}$ at time $t$ as

$$\begin{aligned} D(\mathbf{x},t) &\simeq \operatorname{div}(\mathbf{u}_\mathbf{x}(\mathbf{x},t)) - \frac{\partial \mathrm{lnA}(\mathbf{x},t)}{\partial t} - [\mathbf{u}_\mathbf{x}(\mathbf{x},t)]^T \frac{\partial \ln A(\mathbf{x},t)}{\partial \mathbf{x}}, \\ \operatorname{div}(\mathbf{u}_\mathbf{x}(\mathbf{x},t)) &= \sum_{l=1}^{L} \frac{\partial u_{\mathbf{x}.l}(\mathbf{x},t)}{\partial x_{.l}} \end{aligned} \quad \text{[S8.11]}$$

(see SI Appendix, section 11C for the derivation), where $A(\mathbf{x},t)$ gives the volume of competition kernel for a species occupying niche $\mathbf{x}$ at time $\tau$ ( defined by Eq. S11.16).

#### 8.D Time rescaling and adaptation horizon

For analytical tractability, we assume that $\mathbf{x}$ and $\mathbf{y}$ have no mutational correlation with each other, i.e., $\mathbf{V}_{\mu\mathbf{xy}}(\mathbf{x},t) = \mathbf{0}$ and $\mathbf{V}_{\mu\mathbf{yx}}(\mathbf{x},t) = \mathbf{0}$, and that the direction of $\mathbf{g}_\mathbf{y}(\mathbf{x},t)$ is solely determined by $\mathbf{x}$, i.e.,

$\mathbf{g_y}(\mathbf{x},t) = g_y(\mathbf{x},t)\mathbf{e_{gy}}(\mathbf{x})$ with $|\mathbf{e_{gy}}(\mathbf{x})| = 1$ and $g_y(\mathbf{x},t) = \mathbf{e_{gy}}(\mathbf{x})^T\mathbf{g_y}(\mathbf{x},t)$. Then, from Eqs. S8.6-S8.9, we derive the dynamics of $h(x,t) = \mathbf{e_{gy}}(\mathbf{x})^T\mathbf{h}(\mathbf{x},t)$ as

$$\begin{aligned}
\frac{\partial h(\mathbf{x},t)}{\partial t} &= u_{hy}(\mathbf{x},t)\left[\frac{\partial h(\mathbf{x},t)}{\partial \mathbf{x}^T}\frac{\mathbf{V}_{\mu\mathbf{xx}}(\mathbf{x},t)}{V_{\mu yy}(\mathbf{x},t)}\frac{\partial h(\mathbf{x},t)^T}{\partial \mathbf{x}} + 1\right], \\
u_{hy}(\mathbf{x},t) &= \frac{\mu(\mathbf{x},t)V_{\mu yy}(\mathbf{x},t)K_E(\mathbf{x},t)g_y(\mathbf{x},t)}{2\bar{\rho}(\mathbf{x},t)}, \\
V_{\mu yy}(\mathbf{x},t) &= \mathbf{e_{gy}}(\mathbf{x})^T\mathbf{V}_{\mu\mathbf{yy}}(\mathbf{x},t)\mathbf{e_{gy}}(\mathbf{x}) \qquad \text{[S8.12]}
\end{aligned}$$

(see Eqs. S11.9-S11.12 for the derivation).

If $v_{hy}(\mathbf{x},t)$ has its maximum at $\mathbf{x} = \mathbf{x}^*$ for is sufficiently large $t$, then, in a manner analogous to Eq. 23 in the main text, we can approximately transform Eq. S8.12 by rescaling the time $t$ into $\tau = h(\mathbf{x}^*,t)$ (i.e., $\frac{\partial h(\mathbf{x},t)}{\partial \tau} = \frac{\partial h(\mathbf{x},t)}{\partial t}\frac{dt}{d\tau} = \frac{\partial h(\mathbf{x},t)}{\partial t}\left[\frac{\partial h(\mathbf{x}^*,t)}{\partial t}\right]^{-1}$ and replacement of $t$ in $h(x,t)$ with $\tau$ ) into

$$\frac{\partial h(\mathbf{x},\tau)}{\partial \tau} = v_{hy}(\mathbf{x})\left[\frac{\partial h(\mathbf{x},\tau)}{\partial \mathbf{x}^T}\mathbf{V}_{\mu,\mathbf{x}/y}(\mathbf{x})\frac{\partial h(\mathbf{x},\tau)^T}{\partial \mathbf{x}} + 1\right] \qquad \text{[S8.13]}$$

with

$$\begin{aligned}
v_{hy}(\mathbf{x}) &= \lim_{t\to\infty}\frac{u_{hy}(\mathbf{x},t)}{u_{hy}(\mathbf{x}^*,t)} = \lim_{t\to\infty}\left[\frac{\mu(\mathbf{x},t)V_{\mu yy}(\mathbf{x},t)K_E(\mathbf{x},t)g_y(\mathbf{x},t)\bar{\rho}(\mathbf{x}^*,t)}{\mu(\mathbf{x}^*,t)V_{\mu yy}(\mathbf{x}^*,t)K_E(\mathbf{x}^*,t)g_y(\mathbf{x}^*,t)\bar{\rho}(\mathbf{x},t)}\right], \\
\mathbf{V}_{\mu,\mathbf{x}/y}(\mathbf{x}) &= \lim_{t\to\infty}\frac{\mathbf{V}_{\mu\mathbf{xx}}(\mathbf{x},t)}{V_{\mu\mathrm{yy}}(\mathbf{x},t)}. \qquad \text{[S8.14]}
\end{aligned}$$

If there exists $H(\mathbf{x})$ such that $\lim_{\tau\to\infty}[h(\mathbf{x},\tau) - h(\mathbf{x}^*,\tau)] = H(\mathbf{x})$, then $H(\mathbf{x})$ must satisfy

$$\lim_{t\to\infty}\frac{\partial h(\mathbf{x},\tau)}{\partial \tau} = v_{hy}(\mathbf{x})\left[\frac{\partial H(\mathbf{x})}{\partial \mathbf{x}^T}\mathbf{V}_{\mu,\mathbf{x}/y}(\mathbf{x})\frac{\partial H(\mathbf{x})^T}{\partial \mathbf{x}} + 1\right] = 1, \qquad \text{[S8.15]}$$

from which $H(\mathbf{x})$ can be obtained. Then, the time rescaling from $t$ into $\tau$ for Eqs. S8.10 and S8.11 with $t \to \infty$ yields

$$\begin{aligned}
\mathbf{v_x}(\mathbf{x}) &= \lim_{t\to\infty}\frac{\mathbf{u_x}(\mathbf{x},t)}{u_{hy}(\mathbf{x}^*,t)} \\
&= \lim_{t\to\infty}\frac{1}{u_{hy}(\mathbf{x}^*,t)}\left[-\frac{\mu(\mathbf{x},t)}{2}\frac{K_E(\mathbf{x},t)}{\bar{\rho}(\mathbf{x},t)}\mathbf{V}_{\mu\mathbf{xx}}(\mathbf{x},t)\frac{\partial \mathbf{h}(\mathbf{x},t)^T}{\partial \mathbf{x}}\mathbf{e_{gy}}(\mathbf{x})g_y(\mathbf{x},t)\right] \\
&= \lim_{t\to\infty}\frac{\mu(\mathbf{x},t)V_{\mu yy}(\mathbf{x},t)K_E(\mathbf{x},t)g_y(\mathbf{x},t)\bar{\rho}(\mathbf{x}^*,t)}{\mu(\mathbf{x}^*,t)V_{\mu yy}(\mathbf{x}^*,t)K_E(\mathbf{x}^*,t)g_y(\mathbf{x}^*,t)\bar{\rho}(\mathbf{x},t)}\left[-\frac{\mathbf{V}_{\mu\mathbf{xx}}(\mathbf{x},t)}{V_{\mu yy}(\mathbf{x},t)}\frac{\partial \mathbf{h}(\mathbf{x},t)^T}{\partial \mathbf{x}}\mathbf{e_{gy}}(\mathbf{x})\right] \\
&= -v_{hy}(\mathbf{x})\mathbf{V}_{\mu,\mathbf{x}/y}(\mathbf{x})\frac{\partial H(\mathbf{x})\mathbf{e_{gy}}(\mathbf{x})^T}{\partial \mathbf{x}}\mathbf{e_{gy}}(\mathbf{x}) \\
&= -v_{hy}(\mathbf{x})\mathbf{V}_{\mu,\mathbf{x}/y}(\mathbf{x})\left[\frac{\partial H(\mathbf{x})}{\partial \mathbf{x}}\mathbf{e_{gy}}(\mathbf{x})^T\mathbf{e_{gy}}(\mathbf{x}) + H(\mathbf{x})\frac{\partial \mathbf{e_{gy}}(\mathbf{x})^T}{\partial \mathbf{x}}\mathbf{e_{gy}}(\mathbf{x})\right] \\
&= -v_{hy}(\mathbf{x})\mathbf{V}_{\mu,\mathbf{x}/y}(\mathbf{x})\frac{\partial H(\mathbf{x})}{\partial \mathbf{x}}, \\
\mathbf{v_y}(\mathbf{x}) &= \lim_{t\to\infty}\frac{\mathbf{u_y}(\mathbf{x},t)}{u_{hy}(\mathbf{x}^*,t)} = v_{hy}(\mathbf{x})\mathbf{V}_{\mu,\mathbf{y}/y}(\mathbf{x})\mathbf{e_{gy}}(\mathbf{x}), \\
\mathbf{V}_{\mu,\mathbf{y}/y}(\mathbf{x}) &= \lim_{t\to\infty}\frac{\mathbf{V}_{\mu\mathbf{yy}}(\mathbf{x},t)}{V_{\mu yy}(\mathbf{x},t)}, \\
D(\mathbf{x}) &= \mathrm{div}(\mathbf{v_x}(\mathbf{x})) - [\mathbf{v_x}(\mathbf{x})]^T\frac{\partial \ln A(\mathbf{x})}{\partial \mathbf{x}}. \qquad \text{[S8.16]}
\end{aligned}$$

In addition, we can derive the divergence time $\tau_d(\mathbf{x})$ for a species occupying niche $\mathbf{x}$ from its closest relative as

$$\tau_d(\mathbf{x}) \simeq |H(\mathbf{x})| \qquad \text{[S8.17]}$$

in a manner analogous to "Divergence time" in Methods in the main text, provided that $\int_{\tau_0}^{\infty} v_{hy}(\mathbf{x}(\tau))d\tau$ is finite for $d\mathbf{x}(\tau)/d\tau = \mathbf{v_x}(\mathbf{x})$ and $\mathbf{x}(\tau_0) \neq \mathbf{x}^*$. On the other hand, the general form of $p_d(\mathbf{x})$ (for predicting the phenotypic distance for a species occupying niche $\mathbf{x}$ from its closest relative) has not been derived in our present study.

#### 8.E General form of adaptation front equation for two-dimensional trait spaces

For a two-dimensional trait space $\mathbf{s} = (x, y)^T$, Eq. S8.12 is reduced to

$$\begin{aligned}
\frac{\partial h(x,t)}{\partial t} &= u_{hy}(x,t)\left[\frac{V_{\mu xx}(x,t)}{V_{\mu yy}(x,t)}\left|\frac{\partial h(x,t)}{\partial x}\right|^2 + 1\right], \\
u_{hy}(x,t) &= \frac{\mu(x,t)V_{\mu yy}(x,t)K_E(x,t)g_y(x,t)}{2\bar{\rho}(x,t)}.
\end{aligned} \qquad \text{[S8.18]}$$

The rescaled adaptation front equation, Eq. S8.13, is reduced to

$$\begin{aligned}
\frac{\partial h(x,\tau)}{\partial \tau} &= v_{hy}(x)\left[V_{\mu,x/y}(x)\left|\frac{\partial h(x,\tau)}{\partial x}\right|^2 + 1\right], \\
v_{hy}(x) &= \lim_{t\to\infty}\frac{u_{hy}(x,t)}{u_{hy}(x^*,t)} = \lim_{t\to\infty}\left[\frac{\mu(x,t)V_{\mu yy}(x,t)K_E(x,t)g_y(x,t)\bar{\rho}(x^*,t)}{\mu(x^*,t)V_{\mu yy}(x^*,t)K_E(x^*,t)g_y(x^*,t)\bar{\rho}(x,t)}\right], \\
V_{\mu,x/y}(x) &= \lim_{t\to\infty}\frac{V_{\mu xx}(x,t)}{V_{\mu yy}(x,t)},
\end{aligned} \qquad \text{[S8.19]}$$

provided that the front top is located at $x = x^*$ that maximizes $u_{hy}(x)$ for sufficiently large $t$. By solving $\frac{\partial h(x,\tau)}{\partial \tau} = \frac{\partial h(x^*,\tau)}{\partial \tau} = 1$ with $h(x,\tau) = h(x^*,\tau) + H(x)$, we obtain the adaptation horizon as

$$H(x) = -\frac{x - x^*}{|x - x^*|}\int_{x^*}^{x}\frac{1}{\sqrt{V_{\mu,x/y}(\tilde{x})}}\sqrt{\frac{1}{v_{hy}(\tilde{x})} - 1}d\tilde{x}. \qquad \text{[S8.20]}$$

Eqs. S8.16 and S8.17 are reduced to

$$\begin{aligned}
v_x(x) &= -v_{hy}(x)V_{\mu,x/y}(x)\frac{dH(x)}{dx}, \\
v_y(x) &= v_{hy}(x), \\
D(x) &= \frac{dv_x(x)}{dx} - v_x(x)\frac{d\ln A(x)}{dx}, \\
\tau_d(x) &= |H(x)|.
\end{aligned} \qquad \text{[S8.21]}$$

Regarding the phenotypic distance $p_d(x)$ from a species occupying niche $x$ to its closest relative, we can apply the same formula with Eq. 11 in the main text:

$$p_d(x) = \sigma_a\sqrt{\left[\frac{dH(x)}{dx}\right]^2 + 1}. \qquad \text{[S8.22]}$$

#### 8.F Application example: Isotropic higher-dimensional niche space

We extend the one-dimensional niche space $x$ for the resource competition model defined by Eqs. 1-3 (in the main text) into that for the $L$-dimensional isotropic niche space $\mathbf{x} = (x_{.1}, \ldots, x_{.L})^T$, so that the population dynamics of arbitrary $N$-coexisting species are given by

$$\begin{aligned} F(\mathbf{s}_i; \mathbf{S}, \mathbf{n}) = \frac{1}{n_i}\frac{dn_i}{dt} &= \beta y_i + 1 - \frac{\sum_{j=1}^{N} \alpha(\mathbf{x}_i, \mathbf{x}_j) n_j}{K(r_i)}, \\ \alpha(\mathbf{x}_i, \mathbf{x}_j) &= \exp\left(-\frac{|\mathbf{x}_i - \mathbf{x}_j|^2}{2\sigma_\alpha^2}\right), \\ K(r_i) &= K_0 \exp\left(-\frac{r_i^2}{2\sigma_K^2}\right), \end{aligned} \qquad [\text{S8.23}]$$

where $r_i = |\mathbf{x}_i| = \sqrt{\sum_{l=1}^{L} x_{i.l}^2}$. Clearly, we can express the adaptation front as $y = h(\mathbf{x}, t)$. From Eq. S8.7, we obtain $K_E(\mathbf{x}_i, t) = \sum_{j=1}^{N} \alpha(\mathbf{x}_i, \mathbf{x}_j)\hat{n}_j = [\beta h(\mathbf{x}_i, t) + 1]K(r_i)$. Hence, the general form for the rescaled adaptation front equation, Eq. S8.13, is reduced to

$$\begin{aligned} \frac{\partial h(\mathbf{x}, \tau)}{\partial \tau} &= v_{hy}(r) \left[\sum_{l=1}^{L} \left[\frac{\partial h(\mathbf{x}, t)}{\partial x_{.l}}\right]^2 + 1\right], \\ v_{hy}(r) &= \lim_{t\to\infty} \left[\frac{[\beta h(\mathbf{x}, t) + 1]K(r)}{[\beta h(\mathbf{0}, t) + 1]}\right] = K(r) \end{aligned} \qquad [\text{S8.24}]$$

(by substitution of $\mu(\mathbf{x}, t) = \mu$, $V_{\mu yy}(\mathbf{x}, t) = \sigma_\mu^2$, $V_{\mu,\mathbf{x}/y}(\mathbf{x}, t) = \mathbf{I}_\mathbf{x}$, $K_E(\mathbf{x}, t) = [\beta h(\mathbf{x}, t) + 1]K(r)$, $g_y(\mathbf{x}, t) = \beta$, $\bar{\rho}(\mathbf{x}, t) = \bar{\rho}$, where $\mathbf{I}_\mathbf{x}$ is the $L$-by-$L$ identity matrix). Then, by assuming $\lim_{\tau\to\infty}[h(\mathbf{x}, \tau) - h(\mathbf{0}, \tau)] = H(r)$, we see that

$$\begin{aligned} \lim_{\tau\to\infty} \frac{\partial h(\mathbf{x}, \tau)}{\partial \tau} &= v_{hy}(r) \left[\sum_{l=1}^{L} \left[\frac{\partial H(r)}{\partial x_{.l}}\right]^2 + 1\right] = K(r) \left[\sum_{l=1}^{L} \left[\frac{dH(r)}{dr}\frac{\partial r}{\partial x_{.l}}\right]^2 + 1\right] \qquad [\text{S8.25}] \\ &= K(r) \left[\sum_{l=1}^{L} \left[\frac{dH(r)}{dr}\frac{x_{.l}}{r}\right]^2 + 1\right] = K(r) \left[\left[\frac{dH(r)}{dr}\right]^2 + 1\right] = 1, \end{aligned}$$

from which we get

$$H(r) = -\int_0^r \sqrt{\frac{1}{K(\tilde{r})} - 1} d\tilde{r}. \qquad [\text{S8.26}]$$

Substitution of Eq. S8.26 into Eq. S8.16 yields

$$\begin{aligned} v_y(r) &= v_{hy}(r) = K(r), \\ v_{x.l}(\mathbf{x}) &= -v_{hy}(r)\frac{\partial H(r)}{\partial x_{.l}} = -K(r)\frac{dH(r)}{dr}\frac{\partial r}{\partial x_{.l}} = -K(r)\frac{dH(r)}{dr}\frac{x_{.l}}{r} \\ v_r(r) &= \mathbf{v}_\mathbf{x}(\mathbf{x})^T \mathbf{e}_r(\mathbf{x}) = \begin{pmatrix} v_{x.1}(\mathbf{x}) \\ \vdots \\ v_{x.L}(\mathbf{x}) \end{pmatrix}^T \begin{pmatrix} \frac{\partial r}{\partial x_{.1}} \\ \vdots \\ \frac{\partial r}{\partial x_{.L}} \end{pmatrix} = \sum_{l=1}^{L} \left[-K(r)\frac{dH(r)}{dr}\frac{x_{.l}^2}{r^2}\right] = -K(r)\frac{dH(r)}{dr}, \\ D(r) &= \sum_{l=1}^{L} \frac{\partial v_{x.l}(\mathbf{x})}{\partial x_{.l}} \\ &= \sqrt{\frac{K(r)}{1 - K(r)}} \left|\frac{\partial \log K(r)}{\partial r}\right| \left[K(r) - \frac{1}{2}\right] + \frac{L-1}{r}\sqrt{[1 - K(r)]K(r)}, \end{aligned} \qquad [\text{S8.27}]$$

and $D(0) = L/\left[\sqrt{2}\sigma_K\right]$.

### 9 Extension for multiple geographic regions

#### 9.A Model

We extend the original resource competition model (i.e., Eqs. 1-3 in the main text) for multiple geographic regions by defining population dynamics as

$$\begin{aligned}
\frac{1}{n_i}\frac{dn_i}{dt} &= \beta y_i + 1 - \frac{\sum_{j=1}^{N} \alpha(x_i, z_i; x_j, z_j) n_j}{K(x_i, z_i)}, \\
\alpha(x_i, z_i; x_j, z_j) &= \begin{cases} \exp\left(-\frac{[x_i - x_j]^2}{2\sigma_\alpha^2}\right) & \text{for} \quad |z_i - z_j| = 0 \\ 0 & \text{for} \quad |z_i - z_j| \neq 0 \end{cases}, \\
K(x_i, z_i) &= \tilde{K}_0(z) \exp\left(-\frac{x_i^2}{2\sigma_K^2}\right), \\
\tilde{K}_0(z) &= \exp\left(-\frac{z_i^2}{2\sigma_{Kz}^2}\right), \qquad \text{[S9.1]}
\end{aligned}$$

where $z_i$ have only discrete values, denoted by $z_i = k\Delta z$ with an integer $k$ and a positive constant $\Delta z = 1$.

We assume that successful migration occurs only between the neighboring regions with a rate $\mu_m$ per unit $t$ and per unit population size, and that the effect of individual deaths due to unsuccessful migration is already included in $K(x, z)$ (like as individual deaths due to deleterious mutation is included in $K(x, z)$). Additionally, we assume that the invasion fitness function does not distinguish whether a newly introduced individual in a region is a mutant emerged within that region or an immigrant from the other regions, so that its establishment probability (i.e., invasion probability) is given by

$$f(x', z', y'; \mathbf{x}, \mathbf{z}, \mathbf{y}) = \beta y' + 1 - \frac{\sum_{j=1}^{N} \alpha(x', z'; x_j, z_j) \hat{n}_j}{K(x', z')}. \qquad \text{[S9.2]}$$

Under these assumptions, we can treat the geographic location $z$ as if it is a kind of evolutionary trait, like as $x$ and $y$, under a sufficiently weak slope for $\tilde{K}_0(z)$ (for continuous approximation of $z$). For numerical tractability, phenotypes sharing the same $(x, y)$ but existing in different geographic regions were treated as different species.

#### 9.B Adaptation front equation

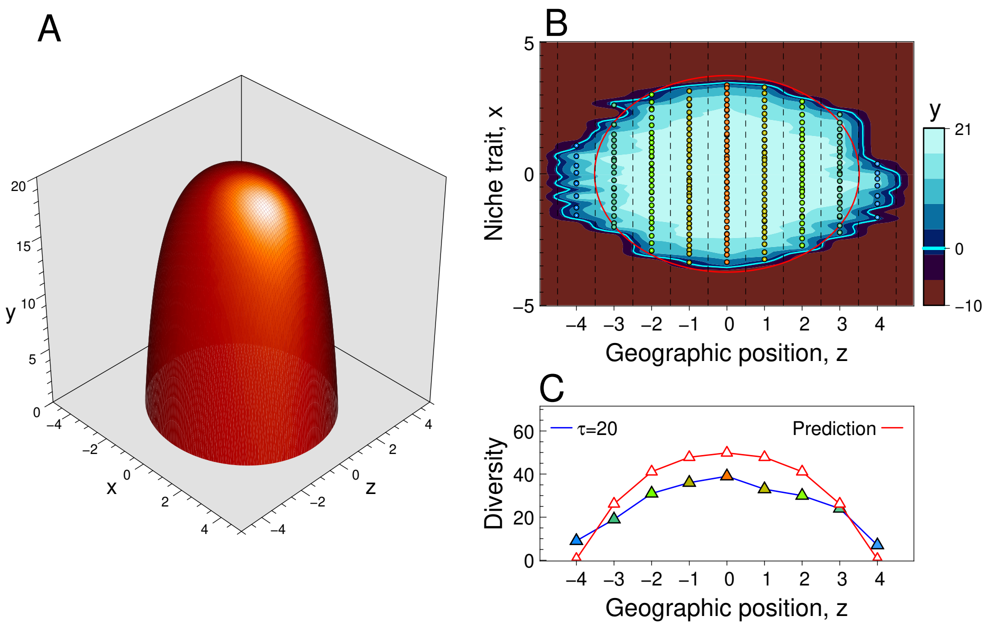

Fig. S15. Adaptation horizon for geographically-extended resource competition model (Fig. 5 in the main text). Panel (A) shows the numerically obtained adaptation horizon, $H(x,z)$, from Eq. S9.3. The horizon is plotted as a surface $y = \max_{x,z}(h(x,z,\tau)) + H(x,z)$ with $\tau = 20$. Panel (B) shows positions of coexisting species on the $(x,z)$-plane at $\tau = 20$. The color gradation indicates the height of adaptation front, $y = h(x,z,\tau)$, on which all coexisting species must be located. The intersection of the adaptation front with the $(x,z)$-plane, i.e., $h(x,z,\tau) = 0$, gives the area of species existence (cyan closed curve), and which is well characterized by $\max_{x,z}(h(x,z,\tau)) + H(x,z) = 0$ (red closed curve). Panel (C) shows the species diversity (i.e., species number) in each geographic region with colored triangles connected with blue line segments. The prediction (red) for each region was given by the width of the red closed curve in panel (B) along the $x$-direction for that region $z$, $\int_{\tau+H(x,z)\geqslant 0} 1dx$, divided by the expected niche width for a species, $\sigma_\alpha$.

Under the constant fitness gradient, $\beta$, along the $y$-direction, the adaptation front is expressed as $y = h(x,z,t)$. For the eco-evolutionary setting considered here, the general form of rescaled adaptation front equation, Eq. S8.13, is reduced to

$$\begin{aligned}\frac{\partial h(x,z,\tau)}{\partial \tau} &= v_{hy}(x,z)\left[\left[\frac{\partial h(x,z,\tau)}{\partial x}\right]^2 + \sigma_{z/y}^2\left[\frac{\partial h(x,z,\tau)}{\partial z}\right]^2 + 1\right],\\ v_{hy}(x,z) &= K(x,z),\\ \sigma_{z/y} &= \sqrt{\frac{\mu_m \Delta z^2}{\mu\sigma_\mu^2}}\end{aligned} \quad \text{[S9.3]}$$

(by substitution of $\mathbf{x} = (x,z)^T$, $\mu(\mathbf{x},t) = \mu + \mu_m$, $V_{\mu yy}(\mathbf{x},t) = \frac{1}{\mu+\mu_m}\mu\sigma_\mu^2$, $V_{\mu\mathbf{xx}}(\mathbf{x},t) = \frac{1}{\mu+\mu_m}\begin{pmatrix}\mu\sigma_\mu^2 & 0\\ 0 & \mu_m\Delta z^2\end{pmatrix}$, $K_E(\mathbf{x},t) = [\beta h(x,z,t)+1]K(x,z)$, $g_y(\mathbf{x},t) = \beta$, $\bar{\rho}(\mathbf{x},t) = \bar{\rho}$).

Then, from $\lim_{\tau\to\infty}\frac{\partial h(\mathbf{x},\tau)}{\partial\tau} = 1$ and $\lim_{\tau\to\infty}[h(\mathbf{x},\tau) - h(\mathbf{0},\tau)] = H(\mathbf{x})$, we see that

$$\left[\frac{\partial H(x,z)}{\partial x}\right]^2 + \sigma_{z/y}^2\left[\frac{\partial H(x,z)}{\partial z}\right]^2 = \frac{1}{K(x,z)} - 1. \quad \text{[S9.4]}$$

Along the $x$- and $z$-axes, respectively, we can easily integrate this equation to obtain

$$\begin{aligned} H(x,0) &= -\frac{x}{|x|}\int_0^x \sqrt{\frac{1}{K(\tilde{x},0)} - 1 d\tilde{x}}, \\ H(0,z) &= -\frac{z}{|z|}\frac{1}{\sigma_{z/y}}\int_0^z \sqrt{\frac{1}{K(0,\tilde{z})} - 1 d\tilde{z}} \end{aligned} \quad \text{[S9.5]}$$

(red dashed curves in Fig 5C and 5D in the main text). Since analytical derivation of the whole shape of $H(x,z)$ is not easy, we numerically obtained $H(x,z)$ by direct time integration of Eq. S9.3, where a slight artificial diffusion was added for the numerical stability. Based on the numerically obtained $H(x,z)$, shown in Fig. S15A, we calculated the expected directional evolution, net-diversification rate, and divergence time, by using Eq. S8.16, expressed as

$$\begin{aligned} v_y(x,z) &= v_{hy}(x,z) = K(x,z), \\ v_x(x,z) &= -v_{hy}(x,z)\frac{\partial H(x,z)}{\partial x} = -K(x,z)\frac{\partial H(x,z)}{\partial x}, \\ v_z(x,z) &= -v_{hy}(x,z)\sigma_{z/y}^2\frac{\partial H(x,z)}{\partial x} = -\sigma_{z/y}^2 K(x,z)\frac{\partial H(x,z)}{\partial z}, \\ D(x,z) &= \frac{\partial v_x(x,z)}{\partial x} + \frac{\partial v_z(x,z)}{\partial z}, \\ \tau_d(x,z) &= |H(x,z)|. \end{aligned} \quad \text{[S9.6]}$$

Regarding $p_d(x,z)$, we expected that

$$p_d(x,z) \leqslant p_{d,\max}(x,z) = \sigma_\alpha \sqrt{1 + \left[\frac{\partial H(x,z)}{\partial x}\right]^2 + \left[\frac{\partial H(x,z)}{\partial z}\right]^2}. \quad \text{[S9.7]}$$

The prediction by Eqs. S9.5-9.7 and the simulated evolution are shown in Figs. 5-6 in the main text and Figs. S15-S16. Note that the adaptation horizon may also be used for predicting the number of species for each geographic region (Fig. S15B and S15C).

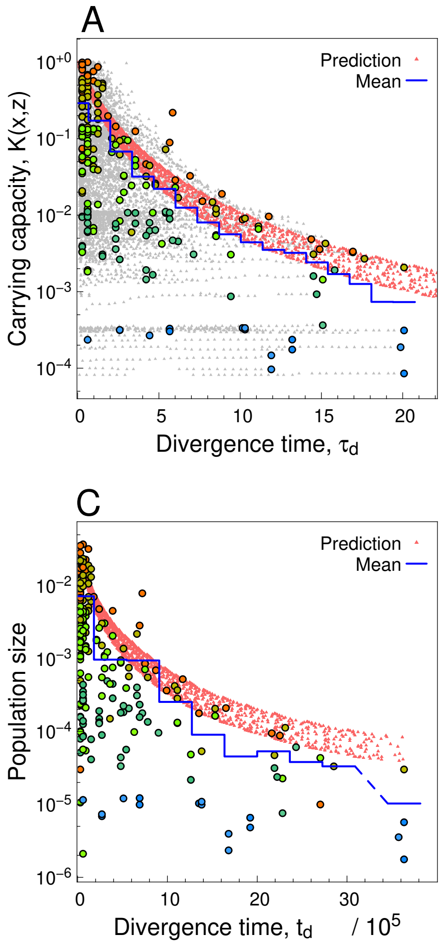

Fig. S16. Prediction by the adaptation front equation for phenotypic distances and divergence times among coexisting species (for the simulated evolution shown in Fig. 5 in the main text). Panel (A) shows the relationship for each species between its divergence time from its closest relative and the carrying capacity for it at $\tau = 20$, in a manner analogous to the third panel of Fig. S2E. The color for each circle indicates the geographic position of each species. The prediction (red dots) shows a set of points $(\tau_d, K(x,z))$ generated from a set of points $(\tau_d, x, z)$ satisfying $\tau_d(x,z) = |H(x,z)|$ in Eq. S9.6. Panel (B) shows the relationship for each species between its phenotypic distance from its closest relative and the carrying capacity for it at $\tau = 20$, in a manner analogous to panel (A). The prediction (red dots) shows a set of points $(p_d, K(x,z))$ generated from a set of points $(p_d, x, z)$ satisfying Eq. S9.7 (inequality sign replaced with equality). Panel (C) shows the relationship between the non-rescaled divergence time, $t_d$, and population size for each coexisting species in a manner analogous to panel (A) (but only for $\tau = 20$). The prediction (red dots) shows a set of points $(t(\tau_d), K(x,z)n_{\max})$ generated from a set of points $(\tau_d, x, z)$ satisfying $\tau_d(x,z) = |H(x,z)|$ in Eq. S9.6, where $n_{\max}$ is the maximum population size among coexisting species at $\tau = 20$, and $t(\tau_d)$ gives the non-rescaled time at $\tau = \tau_d$. Panel (D) shows the relationship between the non-rescaled divergence time, $t_d$, and the effectively-fossilized time, $t_{\rm fos}$, calculated in the same manner as in the bottom panel of Fig. S2F. The prediction (the red line) is $t_d = t_{\rm fos}$.

### 10 SI for Methods

#### 10.A Eqs. 15 and 16 in the main text

We express the adaptation front as a set of points having zero invasion fitness, i.e., $\{(x_0, y_0)|f(x_0, y_0; \mathbf{x}, \mathbf{y}) = 0\}$. Dynamics of the adaptation front can be described with movements of these points, provided that each point moves in parallel with the normal vector of adaptation front at its position. The velocity of such a point $(x_0, y_0)$ can be expressed as

$$\mathbf{u}(x_0, y_0) = \begin{pmatrix} u_x(x_0, y_0) \\ u_y(x_0, y_0) \end{pmatrix} = u(x_0, y_0)\frac{\mathbf{g}(x_0, y_0)}{|\mathbf{g}(x_0, y_0)|}, \quad \text{[S10.1]}$$

where $u(x_0, y_0)$ describes its speed, and $\mathbf{g}(x_0, y_0) = (g_x(x_0, y_0), g_y(x_0, y_0))^T$ (with $g_x(x_0, y_0) = \partial f(x_0, y_0; \mathbf{x}, \mathbf{y})/\partial x_0$ and $g_y(x_0, y_0) = \partial f(x_0, y_0; \mathbf{x}, \mathbf{y})/\partial y_0$) is the fitness gradient vector at $(x_0, y_0)$. Since $f(x_0, y_0; \mathbf{x}, \mathbf{y}) = 0$ always holds, we see by the chain rule that

$$\begin{aligned} 0 = \frac{df(x_0, y_0; \mathbf{x}, \mathbf{y})}{dt} &= \frac{dx_0}{dt}\frac{\partial f(x_0, y_0; \mathbf{x}, \mathbf{y})}{\partial x_0} + \frac{dy_0}{dt}\frac{\partial f(x_0, y_0; \mathbf{x}, \mathbf{y})}{\partial y_0} \\ &\quad + \sum_{j=1}^{N} \frac{dx_j}{dt}\frac{\partial f(x_0, y_0; \mathbf{x}, \mathbf{y})}{\partial x_j} + \sum_{i=1}^{N} \frac{dy_j}{dt}\frac{\partial f(x_0, y_0; \mathbf{x}, \mathbf{y})}{\partial y_j} \\ &= \mathbf{u}(x_0, y_0)^T \mathbf{g}(x_0, y_0) + \sum_{i=1}^{N} \left[\frac{d}{dt}\begin{pmatrix} x_j \\ y_j \end{pmatrix}\right]^T \mathbf{g}^j(x_0, y_0), \end{aligned} \quad \text{[S10.2]}$$

where

$$\mathbf{u}(x_0, y_0) = \begin{pmatrix} \frac{dx_0}{dt} \\ \frac{dy_0}{dt} \end{pmatrix}, \quad \mathbf{g}(x_0, y_0) = \begin{pmatrix} g_x(x_0, y_0) \\ g_y(x_0, y_0) \end{pmatrix}, \quad \mathbf{g}^j(x_0, y_0) = \begin{pmatrix} \frac{\partial f(x_0, y_0; \mathbf{x}, \mathbf{y})}{\partial x_j} \\ \frac{\partial f(x_0, y_0; \mathbf{x}, \mathbf{y})}{\partial y_j} \end{pmatrix}. \quad \text{[S10.3]}$$

$\mathbf{g}^j(x_0, y_0)$ describes the impact of trait change of the $j$th species on the fitness of $(x_0, y_0)$. Since $f(x', y'; \mathbf{x}, \mathbf{y})$ can be treated as a fitness landscape for $(x', y')$, we can intuitively understand Eq. S10.2 as follows. In Eq. S10.2, the first term at the last row describes the fitness change at $(x_0, y_0)$ by the movement of $(x_0, y_0)$ when the fitness landscape is fixed (i.e., when $\mathbf{x} = (x_1, \ldots, x_N)$ and $\mathbf{y} = (y_1, \ldots, y_N)$ are fixed). The second term describes the fitness change for fixed $(x_0, y_0)$, caused by the change of fitness landscape due to the changes of $\mathbf{x}$ and $\mathbf{y}$.

Substituting Eq. S10.1 into Eq. S10.2 gives

$$u(x_0, y_0) = -\frac{1}{|\mathbf{g}(x_0, y_0)|}\sum_{j=1}^{N} \left[\frac{d}{dt}\begin{pmatrix} x_j \\ y_j \end{pmatrix}\right]^T \mathbf{g}^j(x_0, y_0). \quad \text{[S10.4]}$$

By expressing Eq. 14 in the main text in the vector form as

$$\frac{d}{dt}\begin{pmatrix} x_j \\ y_j \end{pmatrix} = \frac{\mu\sigma_\mu^2}{2}\hat{n}_j \mathbf{g}(x_j, y_j), \quad \text{[S10.5]}$$

and by substituting Eq. S10.4 and S10.5 into Eq. S10.1, we get

$$\mathbf{u}(x_0, y_0) = \frac{\mu\sigma_\mu^2}{2}\breve{n}(x_0, y_0)\mathbf{g}(x_0, y_0) \quad \text{[S10.6]}$$

with the "apparent population size," $\breve{n}(x_0, y_0)$, given by

$$\begin{aligned} \breve{n}(x_0, y_0) &= \sum_{j=1}^{N} w_j(x_0, y_0)\hat{n}_j, \\ w_j(x_0, y_0) &= \frac{\mathbf{g}(x_j, y_j)^T[-\mathbf{g}^j(x_0, y_0)]}{|\mathbf{g}(x_0, y_0)|^2}. \end{aligned} \quad \text{[S10.7]}$$

Eqs. S10.6 with S10.7 are identical to Eqs. 15 with 16 in the main text.

As derived in the next subsection,

$$\mathbf{g}^j(x_i, y_i) = \begin{cases} -\mathbf{g}(x_i, y_i) & \text{for} \quad i = j \\ 0 & \text{for} \quad i \neq j \end{cases} \qquad \text{[S10.8]}$$

holds for all $i = 1, \ldots, N$, resulting in that $w_j(x_i, y_i) = 1$ for $i = j$ and zero for otherwise. Hence, $\breve{n}(x_i, y_i) = \hat{n}_i$ holds for all $i = 1, \ldots, N$.

#### 10.B Derivation of Eq. S10.8

For the clarity of explanation, we start with a one-dimensional trait space $x$ that has only a single species of phenotype $x_1$, where the invasion fitness of a mutant phenotype $x'$ is denoted by $f(x'; x_1)$. We expand $f(x'; x_1)$ at an arbitrary fixed point $\hat{x}_1$ close to $x_1$ as

$$\begin{aligned} f(x'; x_1) &= f(\hat{x}_1; \hat{x}_1) + \left[\frac{\partial f(x'; x_1)}{\partial x'}\right]_{x'=x_1=\hat{x}_1} [x' - \hat{x}_1] + \left[\frac{\partial f(x'; x_1)}{\partial x_1}\right]_{x'=x_1=\hat{x}_1} [x_1 - \hat{x}_1] + h.o.t. \\ &= g(\hat{x}_1; \hat{x}_1)[x' - \hat{x}_1] + g^1(\hat{x}_1; \hat{x}_1)[x_1 - \hat{x}_1] + h.o.t.. \end{aligned} \qquad \text{[S10.9]}$$

Note that the change of $x_1$ can affect its equilibrium population size $\hat{n}_1$ as well. Hence, $g^1(\hat{x}_1; \hat{x}_1)$ is the sum of the direct impact of the change of $x_1$ (on the invasion fitness for $x' = \hat{x}_1$) and its indirect impact through the change of $\hat{n}_1$. Since the invasion fitness function assumes population-dynamical equilibrium, we see by definition that $f(x_1; x_1) = 0$ must hold for any $x_1$. Hence,

$$f(x_1; x_1) = [x_1 - \hat{x}_1]^T [g(\hat{x}_1; \hat{x}_1) + g^1(\hat{x}_1; \hat{x}_1)] + h.o.t. = 0 \qquad \text{[S10.10]}$$

must hold for any $x_1$ in the neighborhood of $\hat{x}_1$, from which we see that

$$g^1(\hat{x}_1; \hat{x}_1) = -g(\hat{x}_1; \hat{x}_1), \qquad \text{[S10.11]}$$

as shown in (8).

We extend Eq. S10.11 for arbitrary $N$ coexisting species in a two-dimensional trait space $(x, y)$, as follows. The invasion fitness function is expressed as $f(x', y'; \mathbf{x}, \mathbf{y})$ with $\mathbf{x} = (x_1, \ldots, x_N)$ and $\mathbf{y} = (y_1, \ldots, y_N)$. We expand the invasion fitness function with respect to $(x', y')$, $\mathbf{x}$, and $\mathbf{y}$, respectively at $(\hat{x}_i, \hat{y}_i)$, $\hat{\mathbf{x}} = (\hat{x}_1, \ldots, \hat{x}_N)$, and $\hat{\mathbf{y}} = (\hat{y}_1, \ldots, \hat{y}_N)$, where $i$ is arbitrary chosen from $i = 1, \ldots, N$:

$$\begin{aligned} f(x', y'; \mathbf{x}, \mathbf{y}) &= f(\hat{x}_i, \hat{y}_i; \hat{\mathbf{x}}, \hat{\mathbf{y}}) \\ &\quad + g_x(\hat{x}_i, \hat{y}_i)[x' - \hat{x}_i] + g_y(\hat{x}_i, \hat{y}_i)[y' - \hat{y}_i] \\ &\quad + \sum_{j=1}^{N} g_x^j(\hat{x}_i, \hat{y}_i)[x_j - \hat{x}_j] + \sum_{j=1}^{N} g_y^j(\hat{x}_i, \hat{y}_i)[y_j - \hat{y}_j] + h.o.t. \\ &= \begin{pmatrix} x' - \hat{x}_i \\ y' - \hat{y}_i \end{pmatrix}^T \mathbf{g}(\hat{x}_i, \hat{y}_i) + \sum_{j=1}^{N} \begin{pmatrix} x_j - \hat{x}_j \\ y_j - \hat{y}_j \end{pmatrix}^T \mathbf{g}^j(\hat{x}_i, \hat{y}_i) + h.o.t.., \end{aligned} \qquad \text{[S10.12]}$$

where

$$\begin{aligned} \mathbf{g}(\hat{x}_i, \hat{y}_i) &= \begin{pmatrix} g_x(\hat{x}_i, \hat{y}_i) \\ g_y(\hat{x}_i, \hat{y}_i) \end{pmatrix} = \begin{pmatrix} \frac{\partial f(x', y'; \mathbf{x}, \mathbf{y})}{\partial x'} \\ \frac{\partial f(x', y'; \mathbf{x}, \mathbf{y})}{\partial y'} \end{pmatrix}_{x'=\hat{x}_i, y'=\hat{y}_i, \mathbf{x}=\hat{\mathbf{x}}, \mathbf{y}=\hat{\mathbf{y}}}, \\ \mathbf{g}^j(\hat{x}_i, \hat{y}_i) &= \begin{pmatrix} g_x^j(\hat{x}_i, \hat{y}_i) \\ g_y^j(\hat{x}_i, \hat{y}_i) \end{pmatrix} = \begin{pmatrix} \frac{\partial f(x', y'; \mathbf{x}, \mathbf{y})}{\partial x_j} \\ \frac{\partial f(x', y'; \mathbf{x}, \mathbf{y})}{\partial y_j} \end{pmatrix}_{x'=\hat{x}_i, y'=\hat{y}_i, \mathbf{x}=\hat{\mathbf{x}}, \mathbf{y}=\hat{\mathbf{y}}}. \end{aligned} \qquad \text{[S10.13]}$$

Then for any set of $\mathbf{x} = (x_1, \ldots, x_N)$ and $\mathbf{y} = (y_1, \ldots, y_N)$ in the neighborhood of $\mathbf{x} = \hat{\mathbf{x}}$ and $\mathbf{y} = \hat{\mathbf{y}} =$ and for any $i = 1, \ldots, N$, we require

$$\begin{aligned} f(x_i, y_i; \mathbf{x}, \mathbf{y}) &= \begin{pmatrix} x_i - \hat{x}_i \\ y_i - \hat{y}_i \end{pmatrix}^T \mathbf{g}(\hat{x}_i, \hat{y}_i) + \sum_{j=1}^{N} \begin{pmatrix} x_j - \hat{x}_j \\ y_j - \hat{y}_j \end{pmatrix}^T \mathbf{g}^j(\hat{x}_i, \hat{y}_i) + h.o.t. \\ &= \begin{pmatrix} x_i - \hat{x}_i \\ y_i - \hat{y}_i \end{pmatrix}^T [\mathbf{g}(\hat{x}_i, \hat{y}_i) + \mathbf{g}^i(\hat{x}_i, \hat{y}_i)] + \sum_{i=1, i \neq j}^{N} \begin{pmatrix} x_j - \hat{x}_j \\ y_j - \hat{y}_j \end{pmatrix}^T \mathbf{g}^j(\hat{x}_i, \hat{y}_i) + h.o.t. \\ &= 0, \end{aligned} \quad \text{[S10.14]}$$

from which we see that

$$\mathbf{g}^j(\hat{x}_i, \hat{y}_i) = \begin{cases} -\mathbf{g}(\hat{x}_i, \hat{y}_i) & \text{for } \; i = j \\ 0 & \text{for } \; i \neq j \end{cases} . \quad \text{[S10.15]}$$

This equation is identical to Eq. S10.8. For an arbitrary higher dimensional trait space, the same relationship is readily derived in the same manner.

#### 10.C Eq. 20 in the main text

From time $t$ to time $t + \Delta t$ with $\Delta t$ being infinitesimally small, the point $(x_0, y_0) = (x, h(x,t))$ on the adaptation front moves to $(x + \Delta x, h(x,t) + \Delta y)$ with $\Delta x = u_x(x, h(x,t))\Delta t$ and $\Delta y = u_y(x, h(x,t))\Delta t$. Because $h(x,t) + \Delta y = h(x + \Delta x, t + \Delta t)$ must hold at the leading order, we see that

$$\begin{aligned} h(x,t) + \Delta y &= h(x + \Delta x, t + \Delta t) \\ &= h(x,t) + \frac{\partial h(x,t)}{\partial x} \Delta x + \frac{\partial h(x,t)}{\partial t} \Delta t + h.o.t.. \end{aligned} \quad \text{[S10.16]}$$

By substituting $\Delta x = u_x(x, h(x,t))\Delta t$ and $\Delta y = u_y(x, h(x,t))\Delta t$, and taking limit $\Delta t \to 0$, we get Eq. 20 in the main text.

#### 10.D Eq. 22 in the main text

##### Assumption

We assume that the competition kernel is sufficiently narrow, i.e., $\sigma_\alpha$ is sufficiently small, so that we can introduce a constant parameter $\varepsilon$ such that $\sigma_\alpha \ll \varepsilon \ll \beta \ll 1 \leqslant \sigma_K$, satisfying $\sigma_\alpha = O(\varepsilon^2)$ and $\varepsilon = O(\beta^2)$. In this case, many species can coexist within the niche range of $[x - \varepsilon/2, x + \varepsilon/2]$ around $x$. On this basis, we assume that evolutionary dynamics in $x$ is kept close to evolutionary equilibrium, in a sense that even under $\sigma_\alpha \to 0$ the fitness landscape (i.e., $f(x, y; \mathbf{x}, \mathbf{y})$ as a function of $x$ and $y$) keeps its weak local slopes and curvatures as well as its slow change, satisfying for any set of non-negative integers $\theta$, $\phi$, and $\psi$ (with $\theta + \phi + \psi \leqslant 2$)

$$\frac{\partial^{(\theta+\phi+\psi)} f(x, y; \mathbf{x}, \mathbf{y})}{\partial x^\theta \partial y^\phi \partial t^\psi} = \frac{\partial^{(\theta+\phi+\psi)}}{\partial x^\theta \partial y^\phi \partial t^\psi} \left[ \beta y + 1 - \frac{C(x)}{K(x)} \right] = O(\beta) \quad \text{[S10.17]}$$

with $C(x) = \sum_{j=1}^{N} \alpha(x, x_j) \hat{n}_j$, for any $x$ as long as many species exist within $[x - \varepsilon/2, x + \varepsilon/2]$ through the dynamics. This situation can be attained when the shape of competition density distribution, $C(x)$, is kept similar to that of the carrying capacity distribution, $K(x)$, so that $C(x)/K(x)$ has weak and monotonic slopes over niches where species exist. Such a relationship between $C(x)$ and $K(x)$ is typically observed in the numerically simulated evolution under Gaussian competition kernels (defined by Eq. 2 in the main text), as shown in Panel (D) in Figs. S2-S7 and in Fig. S17A (identical to Figs. S2D) and S17B. Note that Eq. S10.17 seems not hold when the competition kernel is platykurtic (Fig. S17D, identical to Fig. S10D) or asymmetric (Fig. S17C, identical to Fig. S11D), which might explain the low prediction performance of the adaptation front equation for these cases (Figs. S10 and S11) at least in part.

Since the invasion fitness function can be expressed as $f(x, y; \mathbf{x}, \mathbf{y}) = \beta[y - h(x, t)]$, we see from Eq. S10.17 that the adaptation front $y = h(x, t)$ satisfies

$$\frac{\partial^{(\theta+\psi)} h(x,t)}{\partial x^{\theta} \partial t^{\psi}} = \frac{\partial^{(\theta+\psi)} \left[y - f(x, y; \mathbf{x}, \mathbf{y})/\beta\right]}{\partial x^{\theta} \partial t^{\psi}} = O(\beta^0) \qquad \text{[S10.18]}$$

for non-negative integers $\theta$ and $\psi$ with $\theta + \psi \leqslant 2$through the dynamics.

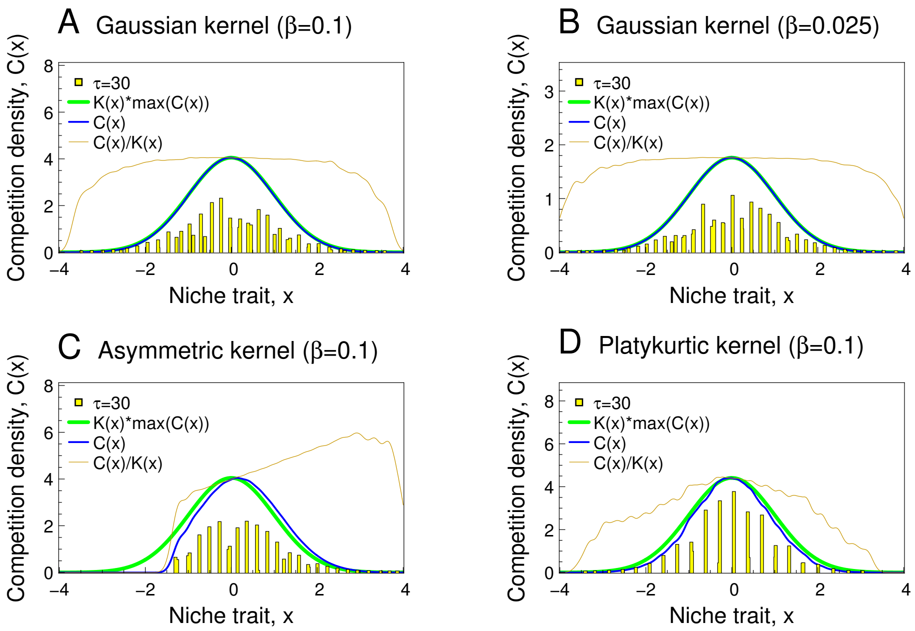

Fig. S17. $C(x)/K(x)$ under different competition kernels in simulated evolution, plotted in the same manner as in Fig. S2D. (A): Gaussian kernel defined by Eq. 2 in the main text with $\beta = 0.1$ (identical to Fig. S2D). (B): Gaussian kernel defined by Eq. 2 in the main text with $\beta = 0.025$. (C): Asymmetric kernel with $\beta = 0.1$ (identical to Fig. S10D). (D): Platykurtic kernel with $\beta = 0.1$ (identical to Fig. S11D).

#### Derivation

First, from a set of $N$ coexisting species, denoted by their ID numbers, $i = 1, \ldots, N$, we collect species with their trait $x$ being included in the range $[x - \varepsilon/2, x + \varepsilon/2]$ and denote the set of their IDs by $\mathcal{I}(x, \varepsilon)$.

By using Eq. S10.18, we transform $\frac{\partial h(x,t)}{\partial t}$ as

$$\begin{aligned}\frac{\partial h(x,t)}{\partial t} &= \frac{\sum_{j\in\mathcal{I}(x,\varepsilon)} \frac{\partial h(x,t)}{\partial t}}{\sum_{j\in\mathcal{I}(x,\varepsilon)} 1} \\ &= \frac{\sum_{j\in\mathcal{I}(x,\varepsilon)} \left[\left[\frac{\partial h(x,t)}{\partial t}\right]_{x=x_j} + \frac{\partial^2 h(x,t)}{\partial x \partial t}[x-x_j] + O(\beta^0)O(|x-x_j|^2)\right]}{\sum_{j\in\mathcal{I}(x,\varepsilon)} 1} \\ &= \frac{\sum_{j\in\mathcal{I}(x,\varepsilon)} \left[\left[\frac{\partial h(x,t)}{\partial t}\right]_{x=x_j} + O(\beta^0)O(\varepsilon)\right]}{\sum_{j\in\mathcal{I}(x,\varepsilon)} 1} \\ &= \frac{\sum_{j\in\mathcal{I}(x,\varepsilon)} \left[\frac{\partial h(x,t)}{\partial t}\right]_{x=x_j}}{\sum_{j\in\mathcal{I}(x,\varepsilon)} 1} + O(\beta^0)O(\varepsilon). \end{aligned} \quad \text{[S10.19]}$$

By using Eqs. 21 and 22 in the main text with $\check{n}(x_j, h(x_j,t)) = \hat{n}_j$, we express $\left[\frac{\partial h(x,t)}{\partial t}\right]_{x=x_j}$ for $j \in \mathcal{I}(x,\varepsilon)$as

$$\begin{aligned}\left[\frac{\partial h(x,t)}{\partial t}\right]_{x=x_j} &= \frac{\mu\sigma_\mu^2\beta}{2}\hat{n}_j\left[1+\left|\frac{\partial h(x,t)}{\partial x}\right|^2_{x=x_j}\right] \\ &= \frac{\mu\sigma_\mu^2\beta}{2}\left[1+\left|\frac{\partial h(x,t)}{\partial x}\right|^2 + O(\beta^0)O(\varepsilon)\right]\hat{n}_j \end{aligned} \quad \text{[S10.20]}$$

which upon substitution into Eq. S10.19 gives

$$\frac{\partial h(x,t)}{\partial t} = \frac{\mu\sigma_\mu^2\beta}{2}\left[1+\left|\frac{\partial h(x,t)}{\partial x}\right|^2\right]\bar{n}(x,t) + O(\beta^0)O(\varepsilon), \quad \text{[S10.21]}$$

$$\bar{n}(x,t) = \frac{\sum_{j\in\mathcal{I}(x,\varepsilon)} \hat{n}_j}{\sum_{j\in\mathcal{I}(x,\varepsilon)} 1} = \frac{\int_{x-\varepsilon/2}^{x+\varepsilon/2}\sum_{i=1}^N \delta(x'-x_j)\hat{n}_j dx'}{\int_{x-\varepsilon/2}^{x+\varepsilon/2}\sum_{i=1}^N \delta(x'-x_j) dx'}, \quad \text{[S10.22]}$$

where $\delta(x)$ is the Dirac's delta function, satisfying $\delta(x) = 0$ for $x \neq 0$, $\delta(x) = \infty$ for $x = 0$, and $\int_{-\infty}^{\infty}\delta(x)dx = 1$. Because $\sigma_\alpha$ is assumed to be much smaller than $\varepsilon$ (i.e., $\sigma_\alpha = O(\varepsilon^2)$), we can approximately substitute $\delta(x)$ by $\alpha(x)/\int_{-\infty}^{\infty}\alpha(x')dx'$ in Eq. S10.22, which results in

$$\bar{n}(x,t) = \frac{\int_{x-\varepsilon/2}^{x+\varepsilon/2}\sum_{i=1}^N \alpha(x'-x_j)\hat{n}_j dx'}{\varepsilon\bar{\rho}(x,t)} + O(\sigma_\alpha/\varepsilon), \quad \text{[S10.23]}$$

$$\bar{\rho}(x,t) = \frac{1}{\varepsilon}\int_{x-\varepsilon/2}^{x+\varepsilon/2}\sum_{j=1}^N \alpha(x'-x_j)dx', \quad \text{[S10.24]}$$

Since $f(x', h(x',t); \mathbf{x}, \mathbf{y}) = 0$ in Eq. 4 in the main text gives $\sum_{i=1}^N \alpha(x'-x_j)\hat{n}_j = [\beta h(x',t)+1]K(x')$, we can further transform Eq. S10.23 into

$$\begin{aligned}\bar{n}(x,t) &= \frac{\int_{x-\varepsilon/2}^{x+\varepsilon/2}[\beta h(x',t)+1]K(x')dx'}{\varepsilon\bar{\rho}(x,t)} + O(\sigma_\alpha/\varepsilon) \\ &= \frac{[\beta h(x,t)+1]K(x)}{\bar{\rho}(x,t)} + O(\beta^0)O(\varepsilon) + O(\sigma_\alpha/\varepsilon). \end{aligned} \quad \text{[S10.25]}$$

By substituting Eq. S10.25 with $O(\varepsilon) = O(\beta^2)$ and $O(\sigma_\alpha/\varepsilon) = O(O(\varepsilon^2)/\varepsilon) = O(\varepsilon) = O(\beta^2)$ into Eq. S10.21, we obtain

$$\frac{\partial h(x,t)}{\partial t} = \frac{\mu\sigma_\mu^2\beta}{2}\frac{[\beta h(x,t)+1]K(x)}{\bar{\rho}(x,t)}\left[1+\left|\frac{\partial h(x,t)}{\partial x}\right|^2\right] + O(\beta^2). \quad \text{[S10.26]}$$

Note that $\bar{\rho}(x,t)/A$ with $A = \int_{-\infty}^{\infty}\alpha(x')\mathrm{dx}' = \sqrt{2\pi}\sigma_\alpha$ approximately gives local species density, because we see that

$$\begin{aligned}\bar{q}(x,t) &= \frac{\int_{x-\varepsilon/2}^{x+\varepsilon/2}\sum_{j=1}^{N}\delta(x'-x)dx'}{\varepsilon} = \frac{\int_{x-\varepsilon/2}^{x+\varepsilon/2}\sum_{j=1}^{N}\frac{\alpha(x'-x_j)}{A}dx'}{\varepsilon}[1+O(\sigma_\alpha/\varepsilon)] \\ &= \frac{\bar{\rho}(x,t)}{A}[1+O(\beta^2)]. \quad \text{[S10.27]}\end{aligned}$$

Provided that the expected between-species niche distance is kept at $\sigma_\alpha$, we expect that $\bar{q}(x,t) = \sigma_\alpha^{-1}$, from which we obtain

$$\bar{\rho} = \lim_{\beta\to 0}\bar{\rho}(x,t) = \lim_{\beta\to 0}\left[\frac{A[1+O(\beta^2)]}{\sigma_\alpha}\right] = \sqrt{2\pi}. \quad \text{[S10.28]}$$

#### 10.E Eq. 27 in the main text

By introducing a positive constant $\Delta x$ such that $\varepsilon \ll \Delta x \ll 1$, we define the local number of species existing within $[x-\Delta x/2, x+\Delta x/2]$ by

$$N^{\Delta x}(x,\tau) = \int_{x-\Delta x/2}^{x+\Delta x/2}\bar{q}(x',\tau)\mathrm{dx}'. \quad \text{[S10.29]}$$

For the simplicity of explanation, we assume that $v_x(x,\tau)$ (i.e., the expected velocity of directional evolution along the $x$-direction for a species occupying niche $x$) is positive within $[x-\Delta x/2, x+\Delta x/2]$, without loss of generality. Then, the change of the local species number from $N^{\Delta x}(x,\tau)$ at time $\tau$ to $N^{\Delta x}(x,\tau+\Delta\tau)$ at time $\tau+\Delta\tau$ can be expressed as

$$\begin{aligned}N^{\Delta x}(x,\tau+\Delta\tau) - N^{\Delta x}(x,\tau) &= \Delta N_{\mathrm{In}}(x-\Delta x/2;\tau,\Delta\tau) - \Delta N_{\mathrm{Out}}(x+\Delta x/2;\tau,\Delta\tau) \\ &\quad +\Delta N_{\mathrm{Bran}}(x,\Delta x;\tau,\Delta\tau) - \Delta N_{\mathrm{Ext}}(x,\Delta x;\tau,\Delta\tau), \quad \text{[S10.30]}\end{aligned}$$

where $\Delta N_{\mathrm{In}}(x-\Delta x/2;\tau,\Delta\tau)$ describes the increase of the local species number by directionally evolving species passing through the left boundary, $x-\Delta x/2$, and thus coming into $[x-\Delta x/2, x+\Delta x/2]$, while $\Delta N_{\mathrm{Out}}(x+\Delta x/2;\tau,\Delta\tau)$ describes the decrease of the local species number by species passing through the right boundary, $x+\Delta x/2$, and thus leaving from $[x-\Delta x/2, x+\Delta x/2]$. The $\Delta N_{\mathrm{In}}(x-\Delta x/2;\tau,\Delta\tau)$ and $\Delta N_{\mathrm{Out}}(x+\Delta x/2;\tau,\Delta\tau)$ are approximately given by

$$\begin{aligned}\Delta N_{\mathrm{In}}(x-\Delta x/2;\tau,\Delta\tau) &\simeq \bar{q}(x-\Delta x/2,\tau)v_x(x-\Delta x/2)\Delta\tau, \\ \Delta N_{\mathrm{Out}}(x+\Delta x/2;\tau,\Delta\tau) &\simeq \bar{q}(x+\Delta x/2,\tau)v_x(x+\Delta x/2)\Delta\tau. \quad \text{[S10.31]}\end{aligned}$$

The remaining two terms at the right-hand side of Eq. S10.30, $\Delta N_{\mathrm{Bran}}(x,\Delta x;\tau,\Delta\tau)$ and $\Delta N_{\mathrm{Ext}}(x,\Delta x;\tau,\Delta\tau)$, respectively describe the numbers of evolutionary branching and extinction that occur within $[x-\Delta x/2, x+\Delta x/2]$ and within $[\tau,\tau+\Delta\tau]$. By using Eq. S10.31, we transform Eq. S10.30 into

$$\begin{aligned}\frac{N^{\Delta x}(x,\tau+\Delta\tau) - N^{\Delta x}(x,\tau)}{\Delta\tau\Delta x} &\simeq \frac{\bar{q}(x-\Delta x/2,\tau)v_x(x-\Delta x/2)}{\Delta x} - \frac{\bar{q}(x+\Delta x/2,\tau)u_x(x+\Delta x/2)}{\Delta x} \\ &\quad +\frac{\Delta N_{\mathrm{Bran}}(x,\Delta x;\tau,\Delta\tau) - \Delta N_{\mathrm{Ext}}(x,\Delta x;\tau,\Delta\tau)}{\Delta\tau\Delta x}. \quad \text{[S10.32]}\end{aligned}$$

Assuming $\Delta\tau \to 0$ and $\Delta x \to 0$ for Eq. S10.32 yields the continuity equation for the locally-averaged species density, $\bar{q}(x,\tau)$:

$$\frac{\partial \bar{q}(x,\tau)}{\partial \tau} = -\frac{\partial[\bar{q}(x,\tau)v_x(x,\tau)]}{\partial x} + \bar{q}(x,\tau)D(x,\tau) \qquad \text{[S10.33]}$$

with

$$D(x,\tau) = \frac{1}{\bar{q}(x,\tau)} \lim_{\Delta\tau\to 0,\Delta x\to 0} \frac{\Delta N_{\mathrm{Bran}}(x,\Delta x;\tau,\Delta\tau) - \Delta N_{\mathrm{Ext}}(x,\Delta x;\tau,\Delta\tau)}{\Delta\tau\Delta x}, \qquad \text{[S10.34]}$$

where $D(x,\tau)$ describes the net diversification rate around $x$, i.e., [the number of branching subtracted by the number of extinction] expected around $x$ per unit time per unit species density.

### 11 Derivation of Eq. S8.6: Higher-dimensional adaptation front equation

As shown below, the derivation is analogous to the case of two-dimensional trait space described in "Methods" in the main text and SI Appendix, section 10A-D.

#### 11.A Derivation of exact adaptation front equation

We express the adaptation front as a set of points having zero invasion fitness, as $\{\mathbf{s}_0 = (x_{0.1},\ldots,x_{0.L})^T | f(\mathbf{s}_0;\mathbf{S}) = 0\}$. Dynamics of the adaptation front can be described with movements of those points, where the velocity of such a point $\mathbf{s}_0$ is described as

$$\frac{d\mathbf{s}_0}{dt} = \mathbf{u}_h(\mathbf{s}_0;\mathbf{S}) = u_h(\mathbf{s}_0)\frac{\mathbf{g}(\mathbf{s}_0;\mathbf{S})}{|\mathbf{g}(\mathbf{s}_0;\mathbf{S})|}, \qquad \text{[S11.1]}$$

with its speed $u_h(\mathbf{s}_0;\mathbf{S})$ and the fitness gradient $\mathbf{g}(\mathbf{s}_0;\mathbf{S})$ at $\mathbf{s}_0$. Since $f(\mathbf{s}_0;\mathbf{S}) = 0$ always holds, we see by the chain rule that

$$\begin{aligned} 0 = \frac{df(\mathbf{s}_0;\mathbf{S})}{dt} &= \left[\frac{d\mathbf{s}_0}{dt}\right]^T \frac{\partial f(\mathbf{s}_0;\mathbf{S})}{\partial \mathbf{s}_0} + \sum_{j=1}^{N}\left[\frac{d\mathbf{s}_j}{dt}\right]^T \frac{\partial f(\mathbf{s}_0;\mathbf{S})}{\partial \mathbf{s}_j} \\ &= \mathbf{u}_h(\mathbf{s}_0;\mathbf{S})^T\mathbf{g}(\mathbf{s}_0;\mathbf{S}) + \sum_{j=1}^{N}\left[\frac{d\mathbf{s}_j}{dt}\right]^T \frac{\partial f(\mathbf{s}_0;\mathbf{S})}{\partial \mathbf{s}_j}. \end{aligned} \qquad \text{[S11.2]}$$

Substitution of Eq. S11.1 and Eq. S8.2, i.e.,

$$\frac{d\mathbf{s}_j}{dt} = \frac{\mu(\mathbf{s}_j)}{2}\hat{n}_j\mathbf{V}_\mu(\mathbf{s}_j)\mathbf{g}(\mathbf{s}_j;\mathbf{S}),$$

into Eq. S11.2 and dropping the subscript "$_0$" from "$\mathbf{s}_0$" gives

$$\mathbf{u}_h(\mathbf{s};\mathbf{S}) = \frac{\mu(\mathbf{s})}{2}\sigma_h(\mathbf{s})^2\breve{n}(\mathbf{s};\mathbf{S})\mathbf{g}(\mathbf{s};\mathbf{S}) \qquad \text{[S11.3]}$$

with

$$\begin{aligned} \breve{n}(\mathbf{s};\mathbf{S}) &= \sum_{j=1}^{N}\frac{\mathbf{g}(\mathbf{s}_j;\mathbf{S})^T\mu(\mathbf{s}_j)\mathbf{V}_\mu(\mathbf{s}_j)[-\mathbf{g}^j(\mathbf{s};\mathbf{S})]}{\mathbf{g}(\mathbf{s};\mathbf{S})^T\mu(\mathbf{s})\mathbf{V}_\mu(\mathbf{s})\mathbf{g}(\mathbf{s};\mathbf{S})}\hat{n}_j \\ \mathbf{g}^j(\mathbf{s}) &= \frac{\partial f(\mathbf{s};\mathbf{S})}{\partial \mathbf{s}_j}, \\ \sigma_h(\mathbf{s})^2 &= \frac{\mathbf{g}(\mathbf{s};\mathbf{S})^T\mathbf{V}_\mu(\mathbf{s})\mathbf{g}(\mathbf{s};\mathbf{S})}{|\mathbf{g}(\mathbf{s};\mathbf{S})|^2}, \end{aligned} \qquad \text{[S11.4]}$$

where $\mathbf{g}^j(\mathbf{s}_i;\mathbf{S}) = 0$ for $i \neq j$ and $\mathbf{g}^i(\mathbf{s}_i;\mathbf{S}) = -\mathbf{g}(\mathbf{s}_i;\mathbf{S})$ hold (see SI Appendix, section 10B). Note that the velocity of the adaptation front, $\mathbf{u}_h(\mathbf{s};\mathbf{S})$, is by definition always parallel with $\mathbf{g}(\mathbf{s};\mathbf{S})$, whereas the velocity of each species, $d\mathbf{s}_i/dt$, may not be parallel with $\mathbf{g}(\mathbf{s}_i;\mathbf{S})$, depending on $\mathbf{V}_\mu(\mathbf{s}_i)$.

Since the adaptation front in the trait space $\mathbf{s} = \begin{pmatrix} \mathbf{x} \\ \mathbf{y} \end{pmatrix}$ is given by $\mathbf{y} = \mathbf{h}(\mathbf{x},t)$ (Eq. S8.4), we can expand the invasion fitness of a mutant $\mathbf{s}' = \begin{pmatrix} \mathbf{x}' \\ \mathbf{y}' \end{pmatrix}$ at $\begin{pmatrix} \mathbf{x}' \\ \mathbf{h}(\mathbf{x}',t) \end{pmatrix}$ as

$$\begin{aligned} f(\mathbf{s}';\mathbf{S}) &= [\mathbf{y}' - \mathbf{h}(\mathbf{x}',t)]^T \mathbf{g}_\mathbf{y}(\mathbf{x}',t) + h.o.t., \\ \mathbf{g}_\mathbf{y}(\mathbf{x}',t) &= \left[\frac{\partial f(\mathbf{s}';\mathbf{S})}{\partial \mathbf{y}'}\right]_{\mathbf{y}'=\mathbf{h}(\mathbf{x}',t)}. \end{aligned} \quad \text{[S11.5]}$$

Hence, we can express the fitness gradient at $\mathbf{s} = \begin{pmatrix} \mathbf{x} \\ \mathbf{h}(\mathbf{x},t) \end{pmatrix}$ on the adaptation front as

$$\mathbf{g}(\mathbf{x},t) = \begin{pmatrix} \mathbf{g}_\mathbf{x}(\mathbf{x},t) \\ \mathbf{g}_\mathbf{y}(\mathbf{x},t) \end{pmatrix} = \begin{pmatrix} \frac{\partial f(\mathbf{s}';\mathbf{S})}{\partial \mathbf{x}'} \\ \frac{\partial f(\mathbf{s}';\mathbf{S})}{\partial \mathbf{y}'} \end{pmatrix}_{\mathbf{x}'=\mathbf{x},\mathbf{y}'=\mathbf{y}=\mathbf{h}(\mathbf{x},t)} = \begin{pmatrix} -\frac{\partial \mathbf{h}(\mathbf{x},t)^T}{\partial \mathbf{x}} \\ \mathbf{I}_\mathbf{y} \end{pmatrix} \mathbf{g}_\mathbf{y}(\mathbf{x},t). \quad \text{[S11.6]}$$

By using Eq. S11.6, we transform Eq. S11.3 into

$$\begin{aligned} \begin{pmatrix} \mathbf{u}_{h\mathbf{x}}(\mathbf{x},t) \\ \mathbf{u}_{h\mathbf{y}}(\mathbf{x},t) \end{pmatrix} &= \mathbf{u}_h\left(\begin{pmatrix} \mathbf{x} \\ \mathbf{h}(\mathbf{x},t) \end{pmatrix};\mathbf{S}\right) \\ &= \frac{\mu(\mathbf{x},t)}{2}\sigma_h(\mathbf{x},t)^2 \breve{n}(\mathbf{x},t) \begin{pmatrix} \left[-\frac{\partial \mathbf{h}(\mathbf{x},t)^T}{\partial \mathbf{x}}\right] \mathbf{g}_\mathbf{y}(\mathbf{x},t) \\ \mathbf{g}_\mathbf{y}(\mathbf{x},t) \end{pmatrix}, \end{aligned} \quad \text{[S11.7]}$$

where $\sigma_h(\mathbf{x},t) = \sigma_h(\mathbf{s};\mathbf{S})$, $\mu(\mathbf{x},t) = \mu(\mathbf{s})$, and $\breve{n}(\mathbf{x},t) = \breve{n}(\mathbf{s};\mathbf{S})$ for $\mathbf{s} = \begin{pmatrix} \mathbf{x} \\ \mathbf{h}(\mathbf{x},t) \end{pmatrix}$. In addition, we easily derive

$$\frac{\partial \mathbf{h}(\mathbf{x},t)}{\partial t} = -\frac{\partial \mathbf{h}(\mathbf{x},t)}{\partial \mathbf{x}^T}\mathbf{u}_{h\mathbf{x}}(\mathbf{x},t) + \mathbf{u}_{h\mathbf{y}}(\mathbf{x},t) \quad \text{[S11.8]}$$

in a manner analogous to the two-dimensional case (SI Appendix, section 10C) from $\mathbf{h}(\mathbf{x},t) + \mathbf{u}_{h\mathbf{y}}(\mathbf{x},t)\Delta t = \mathbf{h}(\mathbf{x} + \mathbf{u}_{h\mathbf{x}}(\mathbf{x},t)\Delta t, t + \Delta t)$ for infinitesimal $\Delta t$ at the leading order.

By substituting Eq. S11.7 into Eq. S11.8, we get the exact adaptation front equation:

$$\frac{\partial \mathbf{h}(\mathbf{x},t)}{\partial t} = \frac{\mu(\mathbf{x},t)}{2}\sigma_h(\mathbf{x},t)^2 \breve{n}(\mathbf{x},t) \left[\frac{\partial \mathbf{h}(\mathbf{x},t)}{\partial \mathbf{x}^T}\frac{\partial \mathbf{h}(\mathbf{x},t)^T}{\partial \mathbf{x}} + \mathbf{I}_\mathbf{y}\right] \mathbf{g}_\mathbf{y}(\mathbf{x},t) \quad \text{[S11.9]}$$

with

$$\begin{aligned} \sigma_h(\mathbf{x},t)^2 &= \frac{\mathbf{g}(\mathbf{s};\mathbf{S})^T \mathbf{V}_\mu(\mathbf{s}) \mathbf{g}(\mathbf{s};\mathbf{S})}{|\mathbf{g}(\mathbf{s};\mathbf{S})|^2} = \frac{\begin{pmatrix} \mathbf{g}_\mathbf{x}(\mathbf{x},t) \\ \mathbf{g}_\mathbf{y}(\mathbf{x},t) \end{pmatrix}^T \begin{pmatrix} \mathbf{V}_{\mu\mathbf{x}\mathbf{x}}(\mathbf{x},t) & \mathbf{V}_{\mu\mathbf{x}\mathbf{y}}(\mathbf{x},t) \\ \mathbf{V}_{\mu\mathbf{y}\mathbf{x}}(\mathbf{x},t) & \mathbf{V}_{\mu\mathbf{y}\mathbf{y}}(\mathbf{x},t) \end{pmatrix} \begin{pmatrix} \mathbf{g}_\mathbf{x}(\mathbf{x},t) \\ \mathbf{g}_\mathbf{y}(\mathbf{x},t) \end{pmatrix}}{\begin{pmatrix} \mathbf{g}_\mathbf{x}(\mathbf{x},t) \\ \mathbf{g}_\mathbf{y}(\mathbf{x},t) \end{pmatrix}^T \begin{pmatrix} \mathbf{g}_\mathbf{x}(\mathbf{x},t) \\ \mathbf{g}_\mathbf{y}(\mathbf{x},t) \end{pmatrix}} \\ &= \frac{\mathbf{g}_\mathbf{y}(\mathbf{x},t)^T \mathbf{B}(\mathbf{x},t) \mathbf{g}_\mathbf{y}(\mathbf{x},t)}{\mathbf{g}_\mathbf{y}(\mathbf{x},t)^T \left[\frac{\partial \mathbf{h}(\mathbf{x},t)}{\partial \mathbf{x}^T}\frac{\partial \mathbf{h}(\mathbf{x},t)^T}{\partial \mathbf{x}} + \mathbf{I}_\mathbf{y}\right] \mathbf{g}_\mathbf{y}(\mathbf{x},t)}, \\ \mathbf{B}(\mathbf{x},t) &= \frac{\partial \mathbf{h}(\mathbf{x},t)}{\partial \mathbf{x}^T}\mathbf{V}_{\mu\mathbf{x}\mathbf{x}}(\mathbf{x},t)\frac{\partial \mathbf{h}(\mathbf{x},t)^T}{\partial \mathbf{x}} + \mathbf{V}_{\mu\mathbf{y}\mathbf{y}}(\mathbf{x},t) \\ &\quad -\frac{\partial \mathbf{h}(\mathbf{x},t)}{\partial \mathbf{x}^T}\mathbf{V}_{\mu\mathbf{x}\mathbf{y}}(\mathbf{x},t) - \left[\frac{\partial \mathbf{h}(\mathbf{x},t)}{\partial \mathbf{x}^T}\mathbf{V}_{\mu\mathbf{x}\mathbf{y}}(\mathbf{x},t)\right]^T. \end{aligned} \quad \text{[S11.10]}$$

If the direction of $\mathbf{g_y}(\mathbf{x}, t)$ is solely determined by $\mathbf{x}$, denoted by $\mathbf{e_{gy}}(\mathbf{x})$ with $|\mathbf{e_{gy}}(\mathbf{x})| = 1$, then by introducing

$$\begin{aligned} g_y(\mathbf{x}, t) &= \mathbf{e_{gy}}(\mathbf{x})^T \mathbf{g_y}(\mathbf{x}, t), \\ y &= \mathbf{e_{gy}}(\mathbf{x})^T \mathbf{y}, \\ h(\mathbf{x}, t) &= \mathbf{e_{gy}}(\mathbf{x})^T \mathbf{h}(\mathbf{x}, t), \end{aligned} \qquad \text{[S11.11]}$$

we can simplify the exact adaptation front equation (Eq. S11.9) as

$$\begin{aligned} \frac{\partial h(\mathbf{x}, t)}{\partial t} &= \mathbf{e_{gy}}(\mathbf{x})^T \frac{\partial \mathbf{h}(\mathbf{x}, t)}{\partial t} \qquad \text{[S11.12]} \\ &= \frac{\mu(\mathbf{x}, t)}{2} \sigma_h(\mathbf{x}, t)^2 \breve{n}(\mathbf{x}, t) \mathbf{e_{gy}}(\mathbf{x})^T \\ &\quad \cdot \left[ \frac{\partial [h(\mathbf{x}, t)\mathbf{e_{gy}}(\mathbf{x})]}{\partial \mathbf{x}^T} \frac{\partial [h(\mathbf{x}, t)\mathbf{e_{gy}}(\mathbf{x})]^T}{\partial \mathbf{x}} + \mathbf{I_y} \right] \mathbf{e_{gy}}(\mathbf{x}) g_y(\mathbf{x}, t) \\ &= \frac{\mu(\mathbf{x}, t)}{2} \sigma_h(\mathbf{x}, t)^2 \breve{n}(\mathbf{x}, t) \left[ \frac{\partial h(\mathbf{x}, t)}{\partial \mathbf{x}^T} \frac{\partial h(\mathbf{x}, t)^T}{\partial \mathbf{x}} + 1 \right] g_y(\mathbf{x}, t), \end{aligned}$$

where $\mathbf{e_{gy}}(\mathbf{x})^T \frac{\partial \mathbf{e_{gy}}(\mathbf{x})}{\partial \mathbf{x}^T} = \mathbf{0}$ is used (derived from $\mathbf{e_{gy}}(\mathbf{x})^T \mathbf{e_{gy}}(\mathbf{x}) = 1$).

#### 11.B Approximate adaptation front equation

In this subsection, we derive that $\breve{n}(\mathbf{x}, t)$ for Eq. S11.12 can be approximated with Eq. S8.7, i.e., $\breve{n}(\mathbf{x}, t) \simeq \bar{n}(\mathbf{x}, t) = K_E(\mathbf{x}, t)/\bar{\rho}(\mathbf{x}, t)$.

##### Preparation

According to (9), as long as $F(\mathbf{s}'; \mathbf{S}, \mathbf{n})$ is ecologically plausible, and mutation sizes are sufficiently small so that population dynamics is kept close to equilibrium through the whole evolutionary dynamics, the fitness function $F(\mathbf{s}_i; \mathbf{S}, \mathbf{n})$ defined by Eq. S8.1 can be approximated with a linear function of $\mathbf{n}$:

$$\begin{aligned} F(\mathbf{s}_i; \mathbf{S}, \mathbf{n}) &\simeq F(\mathbf{s}_i; \mathbf{S}, \hat{\mathbf{n}}) + \sum_{j=1}^{N} \left[ \frac{F(\mathbf{s}_i; \mathbf{S}, \mathbf{n})}{\partial n_j} \right]_{\mathbf{n}=\hat{\mathbf{n}}} [n_j - \hat{n}_j], \\ &= \sum_{j=1}^{N} b(\mathbf{s}_i, \mathbf{s}_j; \mathbf{S}) [n_j - \hat{n}_j] \\ &= r(\mathbf{s}_i; \mathbf{S}) \left[ 1 - \frac{\sum_{j=1}^{N} a(\mathbf{s}_i, \mathbf{s}_j; \mathbf{S}) n_j}{K_E(\mathbf{s}_i; \mathbf{S})} \right], \end{aligned} \qquad \text{[S11.13]}$$

with

$$\begin{aligned} b(\mathbf{s}_i, \mathbf{s}_j; \mathbf{S}) &= \left[ \frac{\partial F(\mathbf{s}_i; \mathbf{S}, \mathbf{n})}{\partial n_j} \right]_{\mathbf{n}=\hat{\mathbf{n}}}, \\ a(\mathbf{s}_i, \mathbf{s}_j; \mathbf{S}) &= \frac{b(\mathbf{s}_j, \mathbf{s}_j; \mathbf{S})}{b(\mathbf{s}_i, \mathbf{s}_i; \mathbf{S})}, \\ K_E(\mathbf{s}_i; \mathbf{S}) &= \sum_{j=1}^{N} a(\mathbf{s}_i, \mathbf{s}_j; \mathbf{S}) \hat{n}_j, \\ r(\mathbf{s}_i; \mathbf{S}) &= -b(\mathbf{s}_i, \mathbf{s}_i; \mathbf{S}) K_E(\mathbf{s}_i; \mathbf{S}). \end{aligned} \qquad \text{[S11.14]}$$

Note that $a(\mathbf{s}_i, \mathbf{s}_i) = 1$ always holds trivially.

Since $\mathbf{s}_i$ and $\mathbf{s}_j$ are located on the adaptation front $y = h(\mathbf{x}, t)$, satisfying $\mathbf{s}_i = \begin{pmatrix} \mathbf{x}_i \\ h(\mathbf{x}_i, t) \end{pmatrix}$ and $\mathbf{s}_j = \begin{pmatrix} \mathbf{x}_j \\ h(\mathbf{x}_j, t) \end{pmatrix}$, we introduce the following notations

$$
\begin{aligned}
a(\mathbf{x}_i, \mathbf{x}_j, t) &= a\left(\begin{pmatrix} \mathbf{x}_i \\ h(\mathbf{x}_i, t) \end{pmatrix}, \begin{pmatrix} \mathbf{x}_j \\ h(\mathbf{x}_j, t) \end{pmatrix}; \mathbf{S}\right), \\
K_E(\mathbf{x}_i, t) &= K_E\left(\begin{pmatrix} \mathbf{x}_i \\ h(\mathbf{x}_i, t) \end{pmatrix}; \mathbf{S}\right), \\
r(\mathbf{x}_i, t) &= r\left(\begin{pmatrix} \mathbf{x}_i \\ h(\mathbf{x}_i, t) \end{pmatrix}; \mathbf{S}\right), \\
\beta &= \min_{\mathbf{x}, t}(|g_y(\mathbf{x}, t)|).
\end{aligned} \tag{S11.15}
$$

On this basis, we specifically describe the two assumptions made in the main text: (i) $\mathbf{x}$ is a niche space undergoing negative frequency-dependent selection with narrow interaction range, and (ii) $y$ is a fundamental trait undergoing a weak and monotonic directional selection.

**Assumption (i):**

$a(\mathbf{x}_i, \mathbf{x}_j, t)$ is always positive (i.e., interaction is competitive), and it is unimodal about $\mathbf{x}_j$ with its peak at $\mathbf{x}_j = \mathbf{x}_i$ (i.e., competition is the most intense when species $i$ and $j$ occupy the same niche position). The width $\sigma_\alpha$ of the competition kernel $a(\mathbf{x}_i, \mathbf{x}_j, t)$ is sufficiently narrow, so that we can introduce $\varepsilon$ such that $\sigma_\alpha \ll \varepsilon \ll \beta$, satisfying $\sigma_\alpha = O(\varepsilon^2)$ and $\varepsilon = O(\beta^2)$, where $\sigma_\alpha$ is defined by

$$
\begin{aligned}
\sigma_a &= \max_{\mathbf{x}, t}\left(\frac{\sqrt{\lambda_{\max}(\mathbf{V}_a(\mathbf{x}, t))}}{A(\mathbf{x}, t)}\right), \\
\mathbf{V}_a(\mathbf{x}, t) &= \int a(\mathbf{x}', \mathbf{x}, t)[\mathbf{x}' - \mathbf{x}][\mathbf{x}' - \mathbf{x}]^T d\mathbf{x}', \\
A(\mathbf{x}, t) &= \int a(\mathbf{x}', \mathbf{x}, t) d\mathbf{x}',
\end{aligned} \tag{S11.16}
$$

and where $\lambda_{\max}(\bullet)$ gives the maximum eigenvalue of a symmetric matrix $\bullet$, and $\int d\mathbf{x}'$means integration over the whole niche space $\mathbf{x}$. Under $\sigma_\alpha \ll \varepsilon \ll \beta$ assumed here, many species can coexist within an $L$-dimensional hyper-cube with its center $\mathbf{x}$ and its side length $\varepsilon$, referred to as the $\varepsilon$-hyper-cube for $\mathbf{x}$, defined by

$$
\mathcal{C}(\mathbf{x}, \varepsilon) = \left\{\mathbf{x}' = (x'_{.1}, \ldots, x'_{.L})^T \middle| \; |x'_{.l} - x_{.l}| \leqslant \varepsilon/2 \mathrm{for} l = 1, \ldots, L\right\}. \tag{S11.17}
$$

**Assumption (ii):**

The fitness gradient along the $y$-direction is weak and monotonic, satisfying

$$
\beta \leqslant g_y(\mathbf{x}, t) \leqslant \max(g_y(\mathbf{x}, t)) = O(\beta) \tag{S11.18}
$$

for a small constant $\beta$, so that evolutionary dynamics in $x$ is kept close to evolutionary equilibrium, in a sense that even under $\sigma_\alpha \to 0$ the fitness landscape (i.e., $f(x, y; \mathbf{x}, \mathbf{y})$) keeps its weak local slopes and curvatures as well as its slow change, satisfying for any set of non-negative integers $\theta$, $\theta'$, $\phi$, and $\psi$ (with $0 < \theta + \theta' + \phi + \psi \leqslant 2$)

$$
\frac{\partial^{(\theta+\theta'+\phi+\psi)}}{\partial x_{.l}^{\theta} \partial x_{.l'}^{\theta'} \partial y^{\phi} \partial t^{\psi}} \left[\frac{f(\mathbf{s}; \mathbf{S})}{g_y(\mathbf{x}, t)}\right] = O(\beta^0), \tag{S11.19}
$$

for $l, l' = 1, \ldots, L$ and for any $\mathbf{x}$ (around which many species exist) and for any $t$ through the evolutionary dynamics. Since $f(\mathbf{s}; \mathbf{S})$ can be expressed as $f(\mathbf{s}; \mathbf{S}) = f\left(\left(\begin{array}{c} \mathbf{x} \\ y\mathbf{e}_{\mathrm{gy}}(\mathbf{x}) \end{array}\right); \mathbf{S}\right) = [y - h(\mathbf{x}, t)]g_y(\mathbf{x}, t) +$ $h.o.t.$, we see from Eq. S11.19 that

$$\frac{\partial^{(\theta+\theta'+\psi)} h(\mathbf{x}, t)}{\partial x_{.l}^{\theta} \partial x_{.l'}^{\theta'} \partial t^{\psi}} \;=\; O(\beta^0). \qquad \text{[S11.20]}$$

In addition, we assume that under Eq. S11.20 the following relationships hold,

$$\begin{aligned} \frac{\partial K_E(\mathbf{x}, t)}{\partial x_{.l}} &= O(\beta^0), \\ \frac{\partial V(\mathbf{x}, t)}{\partial x_{.l}} &= O(\beta^0), \\ V(\mathbf{x}, t) &= \frac{\mu(\mathbf{x}, t)\sigma_h(\mathbf{x}, t)^2 g_y(\mathbf{x}, t)}{2\beta}. \qquad \text{[S11.21]} \end{aligned}$$

for all $l = 1, ..., L$.

#### Derivation

The derivation is analogous to the case of two-dimensional trait spaces (SI Appendix, section 10D). First, from a set of $N$ coexisting species, denoted by their IDs, $i = 1, \ldots, N$, we collect species with their niche traits being included in $\mathcal{C}(\mathbf{x}, \varepsilon)$ and denote the set of their IDs by $\mathcal{I}(\mathbf{x}, \varepsilon)$. By using Eq. S11.20, we transform $\frac{\partial h(\mathbf{x},t)}{\partial t}$ as

$$\begin{aligned} \frac{\partial h(\mathbf{x}, t)}{\partial t} &= \frac{\sum_{j\in\mathcal{I}(x,\varepsilon)} \frac{\partial h(\mathbf{x},t)}{\partial t}}{\sum_{j\in\mathcal{I}(x,\varepsilon)} 1} \\ &= \frac{\sum_{j\in\mathcal{I}(\mathbf{x},\varepsilon)} \left[\frac{\partial h(\mathbf{x},t)}{\partial t}\right]_{\mathbf{x}=\mathbf{x}_j}}{\sum_{j\in\mathcal{I}(\mathbf{x},\varepsilon)} 1} + O(\beta^0)O(\varepsilon). \qquad \text{[S11.22]} \end{aligned}$$

By using Eq. S11.12 with $\breve{n}(x_j, h(x_j, t)) = \hat{n}_j$ and Eq. S11.21, we express $\left[\frac{\partial h(\mathbf{x},t)}{\partial t}\right]_{\mathbf{x}=\mathbf{x}_j}$ for $j \in \mathcal{I}(\mathbf{x}, \varepsilon)$ as

$$\begin{aligned} \left[\frac{\partial h(\mathbf{x}, t)}{\partial t}\right]_{\mathbf{x}=\mathbf{x}_j} &= \frac{\mu(\mathbf{x}_j, t)}{2}\sigma_h(\mathbf{x}_j, t)^2 \hat{n}_j \left[\frac{\partial h(\mathbf{x}, t)}{\partial \mathbf{x}^T}\frac{\partial h(\mathbf{x}, t)^T}{\partial \mathbf{x}} + 1\right]_{\mathbf{x}=\mathbf{x}_j} g_y(\mathbf{x}_j, t) \\ &= \beta V(\mathbf{x}_j, t)\left[\frac{\partial h(\mathbf{x}, t)}{\partial \mathbf{x}^T}\frac{\partial h(\mathbf{x}, t)^T}{\partial \mathbf{x}} + 1\right]_{\mathbf{x}=\mathbf{x}_j} \hat{n}_j \\ &= \beta V(\mathbf{x}, t)\left[\frac{\partial h(\mathbf{x}, t)}{\partial \mathbf{x}^T}\frac{\partial h(\mathbf{x}, t)^T}{\partial \mathbf{x}} + 1\right]\hat{n}_j + O(\beta)O(\varepsilon) \qquad \text{[S11.23]} \end{aligned}$$

By using Eq. S11.23 and $\sum_{i=1}^{N} a(\mathbf{x}', \mathbf{x}_j, t)\hat{n}_j = K_E(\mathbf{x}', t)$ (derived from $F(\mathbf{s}_i; \mathbf{S}, \mathbf{n}) = 0$ in Eq. S11.13), we transform Eq. S11.22 into

$$\begin{aligned}
\frac{\partial h(\mathbf{x},t)}{\partial t} &= \beta V(\mathbf{x},t) \left[ \frac{\partial h(\mathbf{x},t)}{\partial \mathbf{x}^T} \frac{\partial h(\mathbf{x},t)^T}{\partial \mathbf{x}} + 1 \right] \bar{n}(\mathbf{x},t) + O(\beta^0)O(\varepsilon), \\
\bar{n}(\mathbf{x},t) &= \frac{\sum_{j\in\mathcal{I}(\mathbf{x},\varepsilon)} \hat{n}_j}{\sum_{j\in\mathcal{I}(\mathbf{x},\varepsilon)} 1} = \frac{\int_{\mathbf{x}'\in\mathcal{C}(\mathbf{x},\varepsilon)} \sum_{i=1}^{N} \delta_L(\mathbf{x}'-\mathbf{x}_j)\hat{n}_j d\mathbf{x}'}{\int_{\mathbf{x}'\in\mathcal{C}(\mathbf{x},\varepsilon)} \sum_{i=1}^{N} \delta_L(\mathbf{x}'-\mathbf{x}_j) d\mathbf{x}'} \\
&= \frac{\int_{\mathbf{x}'\in\mathcal{C}(\mathbf{x},\varepsilon)} \sum_{i=1}^{N} a(\mathbf{x}',\mathbf{x}_j,t)\hat{n}_j \mathrm{dx}'}{\varepsilon^L \bar{\rho}(\mathbf{x},t)} + O(\sigma_\alpha/\varepsilon), \\
&= \frac{\int_{\mathbf{x}'\in\mathcal{C}(\mathbf{x},\varepsilon)} K_E(\mathbf{x}',t)\hat{n}_j \mathrm{dx}'}{\varepsilon^L \bar{\rho}(\mathbf{x},t)} + O(\sigma_\alpha/\varepsilon), \\
&= \frac{K_E(\mathbf{x}',t)}{\bar{\rho}(\mathbf{x},t)} + O(\beta^0)O(\varepsilon) + O(\sigma_\alpha/\varepsilon)
\end{aligned} \quad \text{[S11.24]}$$

where $\delta_L(\mathbf{x}' - \mathbf{x}_j)$ is the $L$-dimensional Dirac's delta function (satisfying $\delta(\mathbf{x}) = 0$ for $\mathbf{x} \neq \mathbf{0}$, $\delta(\mathbf{x}) = \infty$ for $\mathbf{x} = \mathbf{0}$, and $\int \delta_L(\mathbf{x})d\mathbf{x} = 1$), and

$$\bar{\rho}(\mathbf{x},t) = \frac{1}{\varepsilon^L} \int_{\mathbf{x}'\in\mathcal{C}(\mathbf{x},\varepsilon)} \sum_{j=1}^{N} a(\mathbf{x}',\mathbf{x}_j,t) d\mathbf{x}'. \quad \text{[S11.25]}$$

Finally, by substituting Eq. S11.21, $O(\varepsilon) = O(\beta^2)$, and $O(\sigma_\alpha/\varepsilon) = O(\varepsilon) = O(\beta^2)$ into Eq. S10.24 we obtain

$$\frac{\partial h(\mathbf{x},t)}{\partial t} = \frac{\mu(\mathbf{x},t)\sigma_h(\mathbf{x},t)^2}{2} \bar{n}(\mathbf{x},t) \left[ \frac{\partial h(\mathbf{x},t)}{\partial \mathbf{x}^T} \frac{\partial h(\mathbf{x},t)^T}{\partial \mathbf{x}} + 1 \right] g_y(\mathbf{x},t) + O(\beta^2), \quad \text{[S11.26]}$$

$$\bar{n}(\mathbf{x},t) = \frac{K_E(\mathbf{x},t)}{\bar{\rho}(\mathbf{x},t)} + O(\beta^2). \quad \text{[S11.27]}$$

By using Eqs. S11.11 and S11.12, we can transform Eq. S11.27 into

$$\frac{\partial \mathbf{h}(\mathbf{x},t)}{\partial t} = \frac{\mu(\mathbf{x},t)}{2} \sigma_h(\mathbf{x},t)^2 \bar{n}(\mathbf{x},t) \left[ \frac{\partial \mathbf{h}(\mathbf{x},t)}{\partial \mathbf{x}^T} \frac{\partial \mathbf{h}(\mathbf{x},t)^T}{\partial \mathbf{x}} + \mathbf{I_y} \right] \mathbf{g_y}(\mathbf{x},t) + O(\beta^2). \quad \text{[S11.28]}$$

Comparison of Eq. S11.28 with Eq. S11.9 implies that the approximate adaptation front equation is given by approximation of the apparent population size, $\breve{n}(\mathbf{x},t)$, with the locally-averaged population size, $\bar{n}(\mathbf{x},t)$, in the exact adaptation front equation, Eq. S11.9. Even when the direction of $\mathbf{g_y}(\mathbf{x},t)$ depends on $t$, if Eq. S11.9 can be expressed as

$$\begin{aligned}
\frac{\partial \mathbf{h}(\mathbf{x},t)}{\partial t} &= \mathbf{r}(\mathbf{x},t)\breve{n}(\mathbf{x},t), \\
\mathbf{r}(\mathbf{x},t) &= \frac{\mu(\mathbf{x},t)}{2} \sigma_h(\mathbf{x},t)^2 \left[ \frac{\partial \mathbf{h}(\mathbf{x},t)}{\partial \mathbf{x}^T} \frac{\partial \mathbf{h}(\mathbf{x},t)^T}{\partial \mathbf{x}} + \mathbf{I_y} \right] \mathbf{g_y}(\mathbf{x},t),
\end{aligned} \quad \text{[S11.29]}$$

where the derivatives of $\frac{\partial \mathbf{h}(\mathbf{x},t)}{\partial t}$ and $\mathbf{r}(\mathbf{x},t)$ with respect to $\mathbf{x}$ are both $O(\beta^0)$, then we can derive the same form with Eq. S11.28 in a manner analogous to Eqs. S11.22-S11.28 as

$$\begin{aligned}
\frac{\partial \mathbf{h}(\mathbf{x},t)}{\partial t} &\simeq \frac{\sum_{j\in\mathcal{I}(\mathbf{x},\varepsilon)} \left[\frac{\partial \mathbf{h}(\mathbf{x},t)}{\partial t}\right]_{\mathbf{x}=\mathbf{x}_j}}{\sum_{j\in\mathcal{I}(\mathbf{x},\varepsilon)} 1} = \frac{\sum_{j\in\mathcal{I}(\mathbf{x},\varepsilon)} \mathbf{r}(\mathbf{x}_j,t)\hat{n}_j}{\sum_{j\in\mathcal{I}(\mathbf{x},\varepsilon)} 1} \\
&\simeq \mathbf{r}(\mathbf{x},t)\frac{\sum_{j\in\mathcal{I}(\mathbf{x},\varepsilon)} \hat{n}_j}{\sum_{j\in\mathcal{I}(\mathbf{x},\varepsilon)} 1} \\
&\simeq \mathbf{r}(\mathbf{x},t)\frac{\int_{\mathbf{x}'\in\mathcal{C}(\mathbf{x},\varepsilon)} \sum_{i=1}^{N} a(\mathbf{x}',\mathbf{x}_j,t)\hat{n}_j d\mathbf{x}'}{\int_{\mathbf{x}'\in\mathcal{C}(\mathbf{x},\varepsilon)} \sum_{i=1}^{N} a(\mathbf{x}',\mathbf{x}_j,t) d\mathbf{x}'} \simeq \mathbf{r}(\mathbf{x},t)\frac{K_E(\mathbf{x},t)}{\bar{\rho}(\mathbf{x},t)} \\
&= \frac{\mu(\mathbf{x},t)}{2}\sigma_h(\mathbf{x},t)^2\bar{n}(\mathbf{x},t)\left[\frac{\partial \mathbf{h}(\mathbf{x},t)}{\partial \mathbf{x}^T}\frac{\partial \mathbf{h}(\mathbf{x},t)^T}{\partial \mathbf{x}} + \mathbf{I}_\mathbf{y}\right]\mathbf{g}_\mathbf{y}(\mathbf{x},t). \qquad \text{[S11.30]}
\end{aligned}$$

#### 11.C Derivation of Eq. S8.11

We define the local species density by

$$\begin{aligned}
\bar{q}(\mathbf{x},t) &= \frac{\int_{\mathbf{x}'\in\mathcal{C}(\mathbf{x},\varepsilon)} \sum_{i=1}^{N} \delta_L(\mathbf{x}'-\mathbf{x}_j)d\mathbf{x}'}{\int_{\mathbf{x}'\in\mathcal{C}(\mathbf{x},\varepsilon)} d\mathbf{x}'} = \frac{\int_{\mathbf{x}'\in\mathcal{C}(\mathbf{x},\varepsilon)} \sum_{i=1}^{N} \delta_L(\mathbf{x}'-\mathbf{x}_j)d\mathbf{x}'}{\varepsilon^L} \\
&= \frac{\int_{\mathbf{x}'\in\mathcal{C}(\mathbf{x},\varepsilon)} \sum_{i=1}^{N} a(\mathbf{x}',\mathbf{x}_j,t)d\mathbf{x}'}{\varepsilon^L \int a(\mathbf{x}',\mathbf{x},t)d\mathbf{x}'}[1+O(\sigma_\alpha/\varepsilon)] \\
&= \left[\int a(\mathbf{x}',\mathbf{x},t)d\mathbf{x}'\right]^{-1} \frac{\int_{\mathbf{x}'\in\mathcal{C}(\mathbf{x},\varepsilon)} \sum_{i=1}^{N} a(\mathbf{x}',\mathbf{x}_j,t)d\mathbf{x}'}{\varepsilon^L}[1+O(\beta^2)] \\
&= [A(\mathbf{x},t)]^{-1}\bar{\rho}(\mathbf{x},t)[1+O(\beta^2)]. \qquad \text{[S11.31]}
\end{aligned}$$

In the same manner with the standard fluid dynamics, the continuity equation for one-dimensional niche space (SI Appendix, section 10E) is readily extended as

$$\begin{aligned}
\frac{\partial \bar{q}(\mathbf{x},t)}{\partial t} &= -\mathrm{div}(\bar{q}(\mathbf{x},t)\mathbf{u}_\mathbf{x}(\mathbf{x},t)) + \bar{q}(\mathbf{x},t)D(\mathbf{x},t), \\
\mathrm{div}(\bar{q}(\mathbf{x},t)\mathbf{u}_\mathbf{x}(\mathbf{x},t)) &= \sum_{l=1}^{L} \frac{\partial[\bar{q}(\mathbf{x},t)u_{x.l}(\mathbf{x},t)]}{\partial x_{.l}}, \qquad \text{[S11.32]}
\end{aligned}$$

where $D(\mathbf{x},t)$ describes the net-diversification rate around $\mathbf{x}$, i.e., [the number of branching subtracted by the number of extinction] expected around $\mathbf{x}$ per unit time and per unit species density. From Eqs. S11.31

and S11.32 we derive $\frac{\partial \ln \bar{\rho}(\mathbf{x},t)}{\partial t}$ as

$$\begin{aligned}
\frac{\partial \ln \bar{\rho}(\mathbf{x},t)}{\partial t} &\simeq \frac{\partial \ln[\bar{q}(\mathbf{x},t)A(\mathbf{x},t)]}{\partial t} = \frac{1}{\bar{q}(\mathbf{x},t)}\frac{\partial \bar{q}(\mathbf{x},t)}{\partial t} + \frac{\partial \mathrm{lnA}(\mathbf{x},t)}{\partial t} \\
&= \frac{1}{\bar{q}(\mathbf{x},t)}\left[-\sum_{l=1}^{L}\frac{\partial[\bar{q}(\mathbf{x},t)u_{x.l}(\mathbf{x},t)]}{\partial x_{.l}} + \bar{q}(\mathbf{x},t)D(\mathbf{x},t)\right] + \frac{\partial \mathrm{lnA}(\mathbf{x},t)}{\partial t} \\
&= -\sum_{l=1}^{L}\left[\frac{\partial u_{x.l}(\mathbf{x},t)}{\partial x_{.l}} + \frac{\partial \ln \bar{q}(\mathbf{x},t)}{\partial x_{.l}}u_{x.l}(\mathbf{x},t)\right] + D(\mathbf{x},t) + \frac{\partial \mathrm{lnA}(\mathbf{x},t)}{\partial t} \\
&\simeq -\mathrm{div}(\mathbf{u_x}(\mathbf{x},t)) - \sum_{l=1}^{L}\left[\frac{\partial \ln \frac{\bar{\rho}(\mathbf{x},t)}{A(\mathbf{x},t)}}{\partial x_{.L}}u_{x.l}(\mathbf{x},t)\right] + D(\mathbf{x},t) + \frac{\partial \mathrm{lnA}(\mathbf{x},t)}{\partial t} \\
&= -\mathrm{div}(\mathbf{u_x}(\mathbf{x},t)) - [\mathbf{u_x}(\mathbf{x},t)]^T\frac{\partial \ln \bar{\rho}(\mathbf{x},t)}{\partial \mathbf{x}} + [\mathbf{u_x}(\mathbf{x},t)]^T\frac{\partial \mathrm{lnA}(\mathbf{x},t)}{\partial \mathbf{x}} \\
&\quad + D(\mathbf{x},t) + \frac{\partial \mathrm{lnA}(\mathbf{x},t)}{\partial t}. \qquad [S11.33]
\end{aligned}$$

Hence, when $\frac{\partial \ln \bar{\rho}(\mathbf{x},t)}{\partial t} \simeq 0$ and $\frac{\partial \ln \bar{\rho}(\mathbf{x},t)}{\partial \mathbf{x}} \simeq \mathbf{0}$ both hold, $D(\mathbf{x},t)$ is given by

$$D(\mathbf{x},t) \simeq \mathrm{div}[\mathbf{u_x}(\mathbf{x},t)] - [\mathbf{u_x}(\mathbf{x},t)]^T\frac{\partial \mathrm{lnA}(\mathbf{x},t)}{\partial \mathbf{x}} - \frac{\partial \mathrm{lnA}(\mathbf{x},t)}{\partial t}. \qquad [S11.34]$$
